# Unravelling an Efficient O_2_/Reductant-Dependent Catalytic Route in Heme Peroxygenases for Sustainable Applications

**DOI:** 10.1101/2024.05.25.595845

**Authors:** Di Deng, Zhihui Jiang, Lixin Kang, Langxing Liao, Xiaodong Zhang, Yuben Qiao, Binju Wang, Aitao Li

**Affiliations:** State Key Laboratory of Biocatalysis and Enzyme Engineering, Hubei Key Laboratory of Industrial Biotechnology, School of Life Sciences, Hubei University, #368 Youyi Road, Wuhan, 430062, P.R. China; State Key Laboratory of Physical Chemistry of Solid Surfaces and Fujian Provincial Key Laboratory of Theoretical and Computational Chemistry, College of Chemistry and Chemical Engineering, Xiamen University, Xiamen, 361005, P. R. China

## Abstract

Heme peroxygenases are attractive biocatalysts for incorporating oxygen into the organic molecules using H_2_O_2_ as oxygen source under mild conditions. However, their practical applications are hindered by irreversible oxidative inactivation caused by exogenous H_2_O_2_ usage. Herein, we report a novel catalytic pathway in heme peroxygenases that relies on O_2_ and small-molecule reductants such as ascorbate acid (AscA), dehydroascorbic acid (DHA), gallic acid (GA), or pyrogallol (PA) to drive reactions. For reactions of unspecific peroxygenase (UPO) with either AscA or DHA, experimental and computational studies revealed that DHAA (the hydrated form of DHA) is the actual co-substrate responsible for activating oxygen to generate oxyferryl heme (compound I, Cpd I) as the oxygenation species. Subsequently, we demonstrate the universality of this O_2_/reductant-dependent route across various heme peroxygenases, highlighting its biological significance as monooxygenases. Compared to the conventional H_2_O_2_-dependent process, this innovative route can efficiently eliminate the excessive production of H_2_O_2_, thereby preventing the heme destruction and related enzyme inactivation. Finally, scale-up reactions were performed for the preparations of chiral, value-added products with unprecedented productivity, underscoring the great synthetic capabilities of the developed peroxygenase technology, which paves the way for sustainable and practical applications in various chemical transformations.

## Introduction

Heme-containing peroxygenases are widely distributed in all types of living organisms such as animals, plants, fungi, and prokaryotes[1–2]. These enzymes play essential roles in the oxidative metabolism of diverse exogenous and endogenous substrates. Additionally, they are involved in the biosynthesis of numerous physiologically important compounds and secondary metabolites, thus providing host organisms with adaptive advantages for the colonization of specific ecological and/or nutritional niches [3–7].

**Figure 1.**
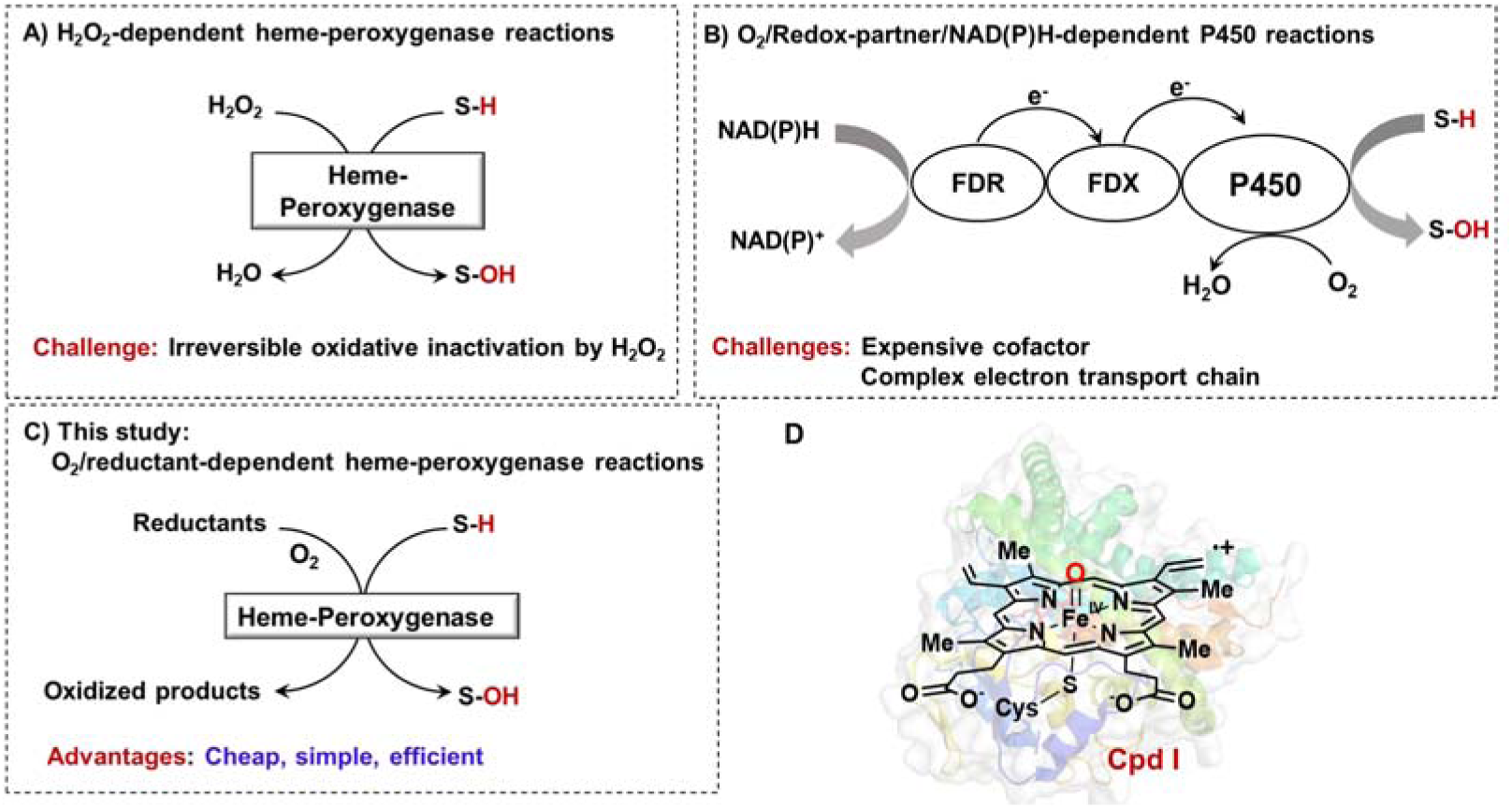
Different catalytic routes of heme-containing oxygenases for generation of oxyferryl heme (compound I, Cpd I) as oxygenation species. **A**. The heme-containing peroxygenases (UPO and P450 peroxygenase) catalyzed reactions *via* the H_2_O_2_-dependent route. **B**. The conventional P450 monooxygenases catalyzed reactions via O_2_/redox-partner/NADPH-dependent route. **C**. The heme-containing peroxygenases catalyzed reactions *via* the O_2_/reductant-dependent route. **D**. Structure of the oxygen-transferring heme species (Compound I, Cpd I).

Among the heme-containing peroxygenases, unspecific peroxygenases (UPOs) has gained significant interest as “dream biocatalysts” due to their ability to introduce oxygen into organic molecules containing inert C-H bonds using hydrogen peroxide (H_2_O_2_) as the oxidant (**Figure 1A**) [1,8–9]. In addition to UPOs, certain cytochrome P450s, such as CYP152 peroxygenases (OleT_JE_, P450_SPα_ and P450_BSβ_), have also been classified as H_2_O_2_-dependent peroxygenases. These enzymes can catalyze hydroxylation or decarboxylation of fatty acids using H_2_O_2_ (**Figure 1A**) [9–13]. Unlike the well-known P450 monooxygenases that require redox partners (ferredoxin and ferredoxin reductases) and reducing agents NAD(P)H for the activation of molecular oxygen (**Figure 1B**) [14,15], both UPOs and P450 peroxygenases exhibit a much simpler catalytic architecture. They can directly generate the reactive oxyferryl heme (compound I, Cpd I) as oxygenation species (**Figure 1D**) from H_2_O_2_, eliminating the need for complex electron transfer chains [1]. Although both UPOs and P450 peroxygenases exhibit obvious advantages as selective oxyfunctionalization catalysts, the catalytic reactions with exogenous H_2_O_2_ suffer from the irreversible oxidative inactivation [16], which largely limits their industrial applications in selective oxyfunctionalization chemistry.

In this study, we have discovered a new catalytic route in heme peroxygenases. Unlike the conventional H_2_O_2_-dependent or O_2_/redox-partner/NADPH-dependent routes, this newly uncovered pathway relies on the activation of O_2_ in the presence of small molecular reductants such as ascorbate acid (AscA), dehydroascorbic acid (DHA), gallic acid (GA) or pyrogallol (PA) (**Figure 1C**). These reductants are naturally occurring in fresh plant tissue or lignified tissue [17]. Extensive experimental and computational studies were conducted to elucidate this unusual mechanism of oxygen activation in both UPO and P450 peroxygenase [18,19]. Furthermore, such O_2_/reductant-dependent route has been demonstrated to be universal across various heme peroxygenases. Finally, the synthesis of chiral, value-added products on a larger scale was conducted to showcase the synthetic capabilities of this developed peroxygenase technology.

## Results and discussions

### Identification of reductant for UPO activity reconstitution

To verify whether UPO can utilize O_2_ as the source of oxygen for oxidation reactions, we investigated the well-known r*Aae*UPO from *Agrocybe aegerita* (PaDa-I variant, r*Aae*UPO) [19] for the hydroxylation of ethylbenzene (**1a**) in the presence of various reducing agents (**Figures 2A and 2B**). The results showed that only the reaction containing r*Aae*UPO, substrate and ascorbate (AscA) resulted in the desired product (*R*)-1-phenylethanol (**1b**) (**Figure 2C**). To further explore if the reaction is an O_2_-dependent process, we conducted the assay under both anaerobic or aerobic conditions, observing that the reaction proceeds efficiently under aerobic conditions but weakly under anaerobic conditions (**Figure 2D**). This suggests that the oxidation reactions catalyzed by r*Aae*UPO are dependent on both AscA and O_2_. Furthermore, the ^18^O_2_ labeling experiment confirmed that the oxygen in the oxidized product originated from the ^18^O_2_ (**Figure 2E**). Finally, we evaluated the catalytic performance of r*Aae*UPO for the hydroxylation of **1a** in the presence of AscA. Impressively, a remarkable total turnover number of 632, 100 ± 37, 073 and over 60 mM of product was obtained (**Figure 2F**), which marks the highest record to date for a free UPO catalyzed reaction.

**Figure 2.**
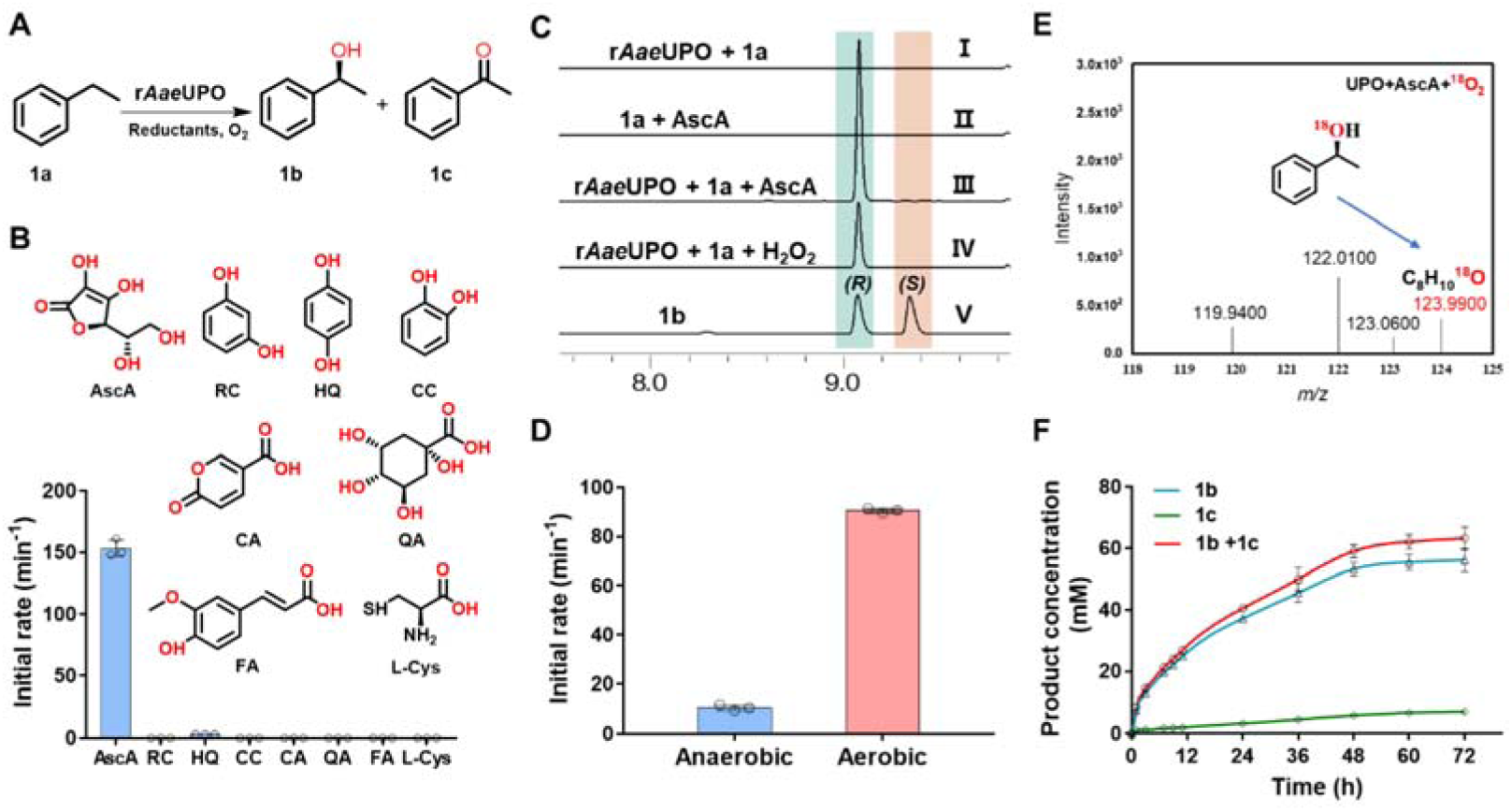
Unspecific peroxygenase r*Aae*UPO-catalyzed benzylic hydroxylation of ethylbenzene (**1a**) *via* the O_2_/reductant-dependent route. **A.** Reaction scheme of r*Aae*UPO-catalyzed benzylic hydroxylation of **1a**. **B.** The reaction activity test of r*Aae*UPO-catalyzed hydroxylation of **1a** in the presence of different reductants. Reactions were performed in potassium phosphate buffer (1 mL, 100 mM, pH 8.0, 30 °C) containing r*Aae*UPO (100 nM), **1a** (10 mM), 5% (v/v) acetonitrile, reductants (20 mM) for 1 h. Reductants used are as follows: Ascorbate (AscA), Resorcinol (RC), Hydroquinone (HQ), Catechol (CC) Coumalic acid (CA), Qinic acid (QA), ferulic acid (FA) and L-Cysteine (L-Cys). **C.** Chiral GC analysis of r*Aae*UPO-catalyzed hydroxylation of **1a** involving AscA as reductant. (□) r*Aae*UPO + **1a**, (□) **1a** + AscA, (□) r*Aae*UPO + **1a** + AscA, (□) r*Aae*UPO + **1a** + H_2_O_2_, (□) standard of (*R*)-1-phenylethanol and (*S*)-1-phenylethanol. **D.** r*Aae*UPO-catalyzed hydroxylation of **1a** by AscA under aerobic and anaerobic conditions for 1 h. Reactions were performed in potassium phosphate buffer (1 mL, 100 mM, pH 8.0, 30 °C) containing r*Aae*UPO (100 nM), **1a** (10 mM), 5% (v/v) acetonitrile, AscA (10 mM). **E.** Probing the origin of oxygen introduced into the **1a** by r*Aae*UPO using labeled ^18^O. **F.** Time course of r*Aae*UPO-catalzyed hydroxylation of **1a** with AscA under aerobic conditions. Reactions were performed in potassium phosphate buffer (100 mM, pH 8.0, 30°C) containing r*Aae*UPO (100 nM), AscA (400 mM), **1a** (100 mM), 5% (m/v) 2-hydroxypropyl-β-cyclodextrin. The data shown in B, D and F are presented as mean value□±□SD (standard deviations) of three biological replicates.

### Identification of real co-substrate for UPO-catalyzed reactions in presence of AscA

It was known that AscA can be smoothly oxidized in aqueous solution to form dehydroascorbic acid (DHA), which then undergoes hydrolysis to produce 2,3-diketo-L-gulonic acid (DKG) [20, 21]. Additionally, DHA is primarily present as the hydrated form (DHAA) in water [21] (**Figure 3A**). Therefore, we proposed that the further oxidized products (DHA or its hydrated form DHAA, or DKG) could potentially act as the actual co-substrates responsible for the reactions. To test this hypothesis, the catalytic activity (initial reaction rate) of r*Aae*UPO was compared by adding AscA, DHA or DKG as the reductant. The results showed that the initial reaction rate reached 144 min^-1^ in the presence of DHA, which is 58% faster than that observed with AscA. No activity was detected when DKG was used. These findings tend to support the hypothesis that DHA (or DHAA) may be the actual reducing co-substrate responsible for the UPO-catalyzed oxidative reactions (**Figure 3B**). Subsequently, the experiments were conducted under both anaerobic and aerobic conditions with the addition of DHA. Like the case of AscA, significant catalytic activity was only achieved under aerobic conditions (**Figure 3C**), indicating that both DHA and O_2_ are essential for the reaction.

To further confirm the critical role of DHA in UPO-catalyzed reactions, DL-Dithiothreitol (DTT), a stronger reductant capable of reducing DHA back to AscA (and also inhibiting the oxidation of AscA to DHA) [22], was added into the reactions containing either AscA or DHA. For the reactions containing 10 mM AscA, it was observed that increasing amounts of DTT resulted in stronger inhibition of the reaction. Complete inhibition occurred when equal amounts of DTT (10 mM) were added (**Figure 3D**). When using DHA as co-substrate, the addition of DTT led to a remarkable decrease in enzyme activity (**Figure 3E**) as the DTT reduced DHA back to AscA. Thus, these experimental findings provide further compelling evidence to support our initial hypothesis that DHA (or DHAA), rather than AscA, serves as the actual co-substrate responsible for the r*Aae*UPO catalyzed reactions.

Next, to investigate whether the consumption of DHA is dependent on r*Aae*UPO, the consumption rate of DHA was monitored in a buffer solution with and without r*Aae*UPO. It was observed that the presence of r*Aae*UPO led to a six-fold enhancement in the rate of DHA consumption, compared to the reaction without enzyme (**Figure 3F**). By contrast, the AscA consumption was found to be considerably lower as compared to DHA (**Figure 3F**). This finding suggests that DHA (or DHAA) may readily enter the active pocket of r*Aae*UPO, whereas AscA may not. Thus, these results suggest a direct interaction between UPO and the reducing agent DHA (or DHAA). To further check if H_2_O_2_ can be generated in the buffer solution in presence of AscA or DHA, control experiments with or without addition of r*Aae*UPO have been conducted. In both scenarios, it was found that the amount of H_2_O_2_ produced is negligible compared to the product formed under the same reaction conditions, indicating the oxidative reactions are essentially independent on the exogenous H_2_O_2_ generated in the buffer solution (**Figure 3G**).

**Figure 3.**
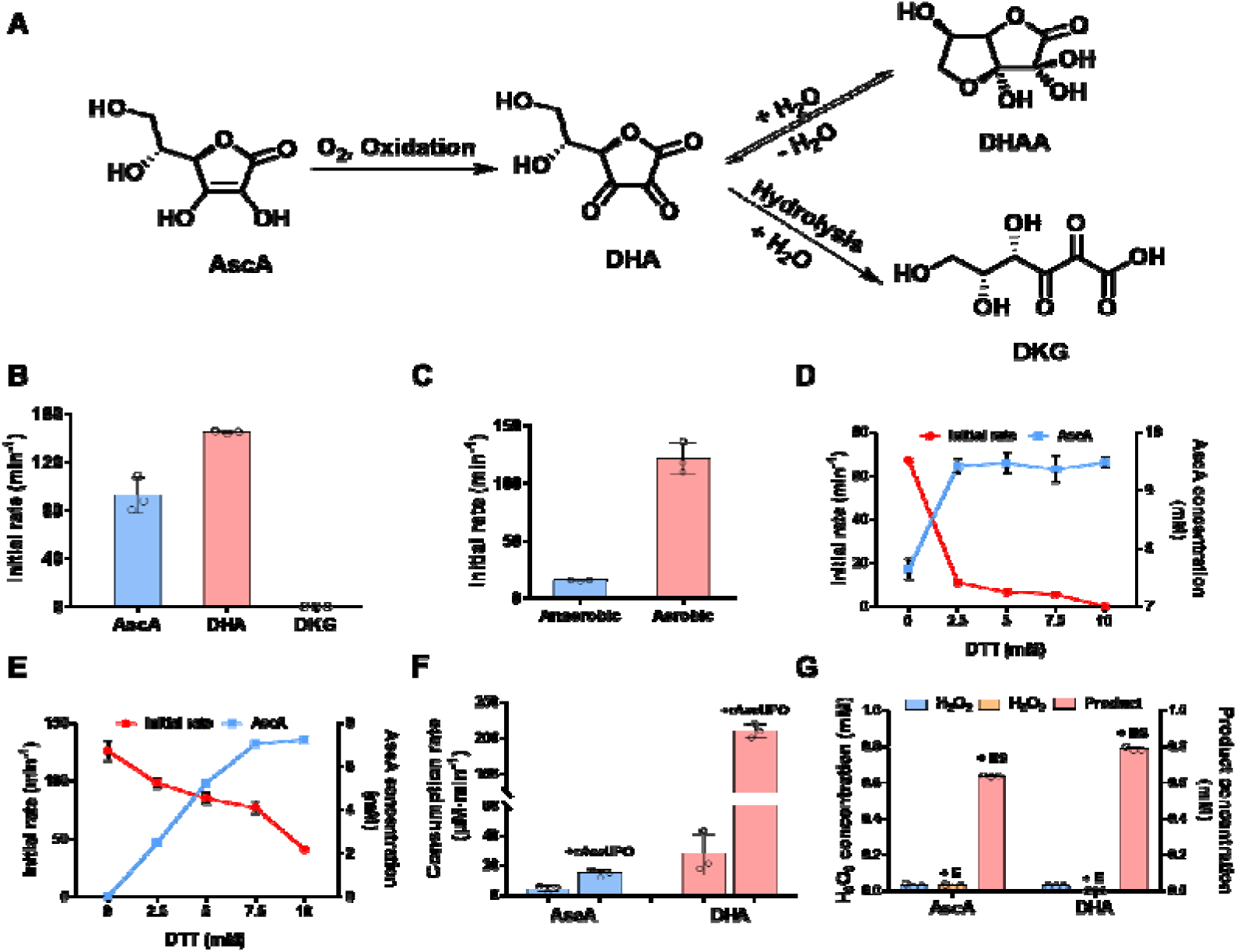
Identification of DHA as the actual co-substrate for r*Aae*UPO catalyzed oxidation reactions. A. The autooxidation of AscA to form DHA (or its hydrated hemiacetal DHAA), followed by hydrolysis to give DKG in aqueous solutions. B. Comparison of the reaction rate by adding AscA, DHA or DKG as reducing agent. C. r*Aae*UPO catalyzed oxidation reactions by DHA under aerobic and anaerobic conditions. D. The effect of DTT on r*Aae*UPO-catalyzed oxidation of 1a using 10 mM AscA as reducing agent. E. The effect of DTT on r*Aae*UPO-catalyzed oxidation of 1a using 10 mM DHA as reducing agent. For the determination of initial reaction rate, all the reactions were performed in potassium phosphate buffer (1 mL, 100 mM, pH 8.5, 30 °C) containing r*Aae*UPO (100 nM), 1a (10 mM), 5% (v/v) acetonitrile, AscA (10 mM), DHA (10 mM), DKG (10 mM) or DTT (0 ∼ 10 mM) for 1 h. F. The consumption rate of DHA and AscA in buffer solution (0.1 M, pH 8.0) with and without r*Aae*UPO addition (800 nM enzyme, 5 mM DHA for 5 min reaction, 5 mM AscA for 60 min reaction). G. Determination of H_2_O_2_ production (with or without enzyme addition) and product (1b) formation under the same reaction conditions. H_2_O_2_ or product (1b) was detected when 10 mM AscA or DHA was incubated at 30 °C in buffer solution (0.1 M, pH 8.0) for 1 h. The data shown in B, C, D, E, F and G are presented as mean value□±□SD (standard deviations) of three biological replicates.

### Mechanistic study of the O_2_ activation by UPO in presence of DHA

To unravel the catalytic mechanism of UPO in presence of DHA, extensive computational studies were conducted. It is worth noting that DHA has been previously described as a poor one-electron reductant [23], which aligns with our calculations (**Supplementary Figure 1**). Furthermore, DHA lacks redox-active OH groups, suggesting it may not act as a reducing agent for O_2_ activation. Thus, the hydrated hemiacetal form of DHA, DHAA, emerged as a potential candidate (**Figure 4A**). Firstly, density functional theory (DFT) calculations demonstrated that the hydration of DHA to generate DHAA is an exothermic process, with a ΔG of -2.3 kcal/mol (**Supplementary Figure 2**). This suggests that DHA is likely dominated by the hydrated form in an aqueous environment, which is in accordance with the previous finding [21]. In addition, our MD simulations show that DHAA can be stably bound in the active site of UPO (**Supplementary Figure 3**), where DHAA is anchored by H-bonding interactions with E196, R189 and T192 (**Figure 4B**). This observation was further supported by MM/GBSA calculations (**Supplementary Table 1**), showing that DHAA exhibited a high binding affinity in the active site of UPO (ΔG= - 27.9 kcal/mol). For a comparison, MD simulations were also performed for the docked AscA. However, AscA failed to stably bind in the active site and escaped after ∼1 ns simulation. Based on these findings, we employed DHAA as the reductant for the subsequent mechanistic study of O_2_ activation by UPO.

Figure 4 displays the QM/MM predicted mechanism of O_2_ activation by UPO. Since the current O_2_ activation process may involve multi-reference characters from heme, Fe and oxygen, the DFT functional of OLYP was selected [24]. Our QM/MM calculations show that the reactant complex of UPO/DHAA/O_2_ has an octuplet ground state (S=7/2), where two single electrons are located on the O_2_ moiety, while the other five single electrons are located on Fe, respectively. The low-spin state (S=1/2), in which two single electrons are located on the O_2_ moiety and three single electrons are located on Fe (^2^RC), is 1.8 kcal/mol higher in energy than the high-spin octuplet state. We first investigated the transfer of H1 atom to the O_2_ moiety, which could be coupled to the O_2_ binding and thus generate an •OOH radical species. Our QM/MM calculations show that such HAT from the high-spin RC is highly unfavorable kinetically (**Supplementary Figure 4)**. Instead, such HAT reaction is favorable from the low-spin ^2^RC, involving a barrier of 18.4 kcal/mol (^2^RC→^2^TS1). The reaction generates an •OOH radical that is bound to the Fe center in → ^2^IM1 (Fe---O1 distance: 1.89 Å). Starting from ^2^IM1, the proton transfer from Ox of DHAA to Glu196 is coupled to the electron transfer from DHA to Fe(III)-•OOH radical species, leading to Cpd 0 in IM2. Meanwhile, a cleavage of the C-C bond of DHAA occurs during this PCET process. It is seen that such a PCET process is quite facile, with a small barrier of only 0.3 kcal/mol (^2^IM1→^2^TS2).

Starting from ^2^IM2, we have compared two competing pathways, one involves the proton transfer from the protonated Glu196 to the distal O of Cpd 0, which triggers the O-O cleavage and affords the active species of Cpd I. This pathway involves a barrier of 5.1 kcal/mol (^2^IM2→^2^TS3). In the alternative pathway, the proton transfer from the protonated Glu196 to the proximal O of Cpd 0 leads to the Fe(III)-OOH intermediate, which involves a barrier of 6.6 kcal/mol at OLYP (^2^IM2→^2^TS3’). Given the GGA functional of OLYP may underestimate the barrier of O-O cleavage of Cpd 0, we have tested the hybrid functional of B3LYP, which has been extensively employed to study the O-O cleavage of Cpd 0 [25–27] With the functional of B3LYP, the calculated barriers are 4.2 kcal/mol for the Fe(III)-H_2_O_2_ formation and 9.7 kcal/mol for Cpd I formation (Figure 4B), suggesting that the formation of Fe(III)-H_2_O_2_ is kinetically favored over that of the direct Cpd I generation. Notably, the Fe(III)-H_2_O_2_ intermediate is well anchored in the active site by H-bonding interactions with Glu196 and dehydrogenated DHAA [2-(4-Hydroxy-2-oxooxolan-3-yl)oxy-2-oxoacetic acid]. In addition, the formation of Cpd I is highly favored thermodynamically over that of H_2_O_2_ (Figure 4A). Due to these factors, the Fe(III)-H_2_O_2_ intermediate would preferably convert to Cpd I *via* the O-O heterolysis mechanism (Fe(III)-H_2_O_2_→Cpd 0→Cpd I) (see proposed catalytic cycle in **Supplementary Figure 5**). Such mechanism is quite similar to that of H_2_O_2_ activation by P450_SPα_ [28]. Further calculations indicate that the dehydrogenated DHAA can undergo the hydrolysis to yield oxalic acid and L-threonic acid-1,4-lactone (ThrOL) within the enzyme binding pocket (**Supplementary Figure 6A**). MD simulations show that the byproduct of oxalic acid can readily diffuse out of the active, which is coupled to the entry of the substrate (**Supplementary Figure 6C-D**). Such finding agrees with our experimental data that the large amount of oxalic acid was detected with GC-MS analysis (**Supplementary Figure 7**). Subsequent QM/MM calculations show that the DHA-mediated oxygen activation by P450_SPα_ follows the similar mechanism as the UPO/DHA/O_2_ system (**Supplementary Figure 8**), suggesting that other analogous enzymes (*Mro*UPO, P450_BSβ_, *Cfu*CPO, OleT_JE_) may follow the common mechanism as demonstrated in r*Aae*UPO and P450_SPα_.

Thus, our calculations support that DHAA can serve as an efficient reductant for oxygen activation by UPOs, which aligns with our experimental findings. Such reactions of the UPO/DHAA/O_2_ system resemble LPMOs, where the presence of O_2_ and reducing agents (e.g. ascorbate acid) can lead to the “in-situ” H_2_O_2_ formation [29–31]. Nevertheless, the H_2_O_2_ intermediate generated from the UPO/DHAA/O_2_ system is ligated to Fe(III) and well anchored by the surrounding H-bonding networks, enabling its efficient conversion to Cpd I and avoiding its release from the active site. This is also consistent with our control experiments that almost no H_2_O_2_ was detected during the reaction with either AscA or DHA (Figure 3G).

**Figure 4.**
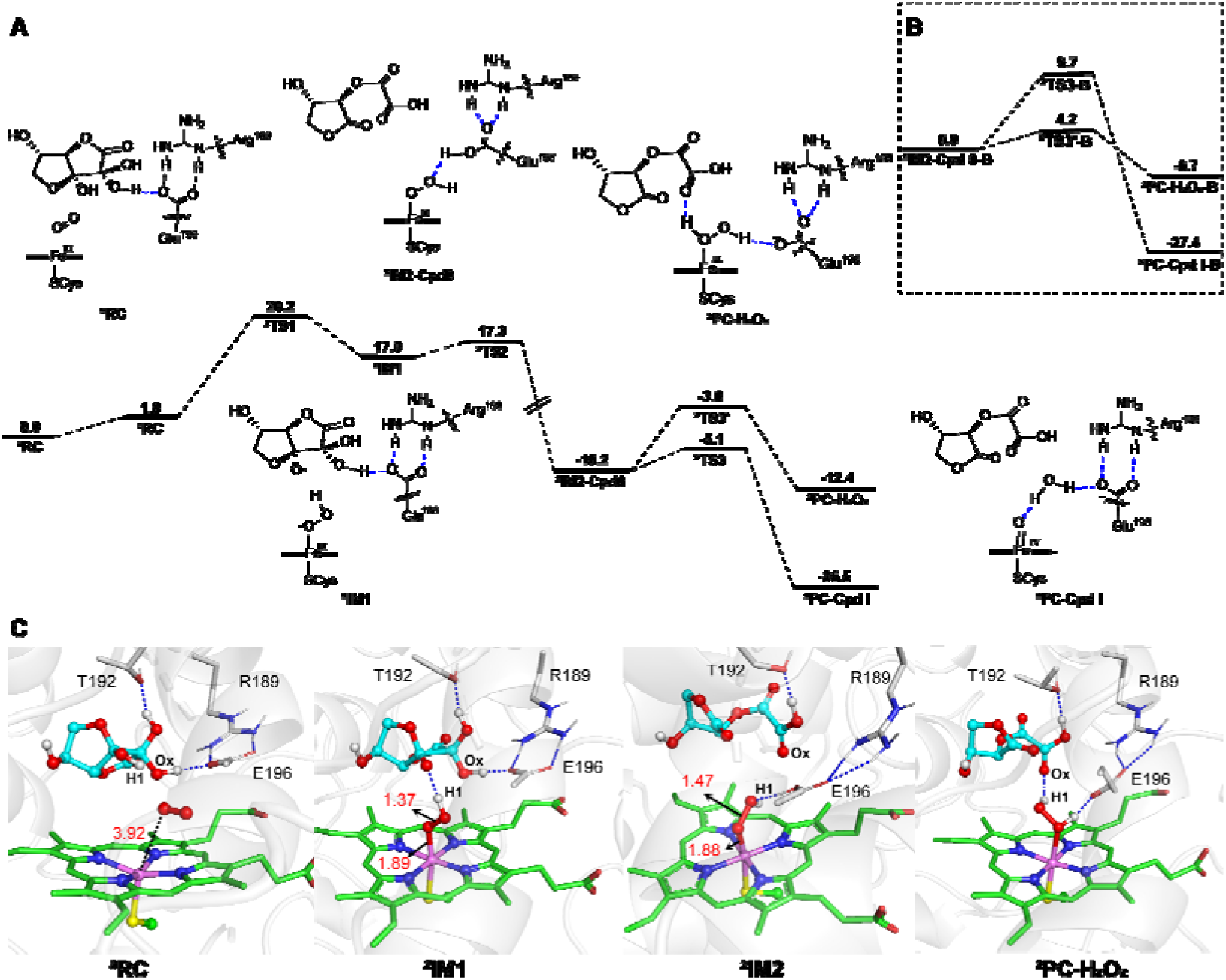
Mechanistic study of the O_2_ activation by UPO in presence of DHAA. A. QM(OLYP/def2-TZVP)/MM calculated mechanism and relative profile (in kcal/mol) for Cpd I and Fe(III)-H_2_O_2_ formation from the UPO/DHAA/O_2_ system. B. QM(B3LYP-D3/def2-TZVP)/MM calculated energy profile (in kcal/mol) for Cpd I formation vs Fe(III)-H_2_O_2_ formation from the Cpd 0 intermediate. C. QM(OLYP/def2-SVP)/MM-optimized geometries of key species involved in the reaction.

### Identification of Gallic acid and Pyrogallol as Co-substrates for UPO

To investigate the potential of other reducing agents as co-substrates for UPO-catalyzed oxidation reactions, in addition to DHA and AscA found in fresh plant tissue, we hypothesized that polyphenolic compounds abundant in lignified tissue could also serve a similar role [17]. To test this hypothesis, we individually examined polyphenolic compounds such as gallic acid (GA) and pyrogallol (PA) in the r*Aae*UPO-catalyzed hydroxylation of **1a** (Figure 5A). As excepted, both GA and PA exhibited excellent catalytic activity as co-substrates, with the initial reaction rate of 265 min^-1^ and 141 min^-1^, respectively (Figure 5B), which are comparable or even superior to those observed with DHA as co-substrate. Consistent with previous findings, it was found that the reaction could only proceed under the aerobic conditions for both GA and PA (Figure 5B). Moreover, the addition of UPO enzyme significantly accelerated the consumption rate of both GA and PA (Figure 5C). These findings collectively support the notion that GA and PA can also function as reducing agents in UPO-catalyzed oxidation reactions.

**Figure 5.**
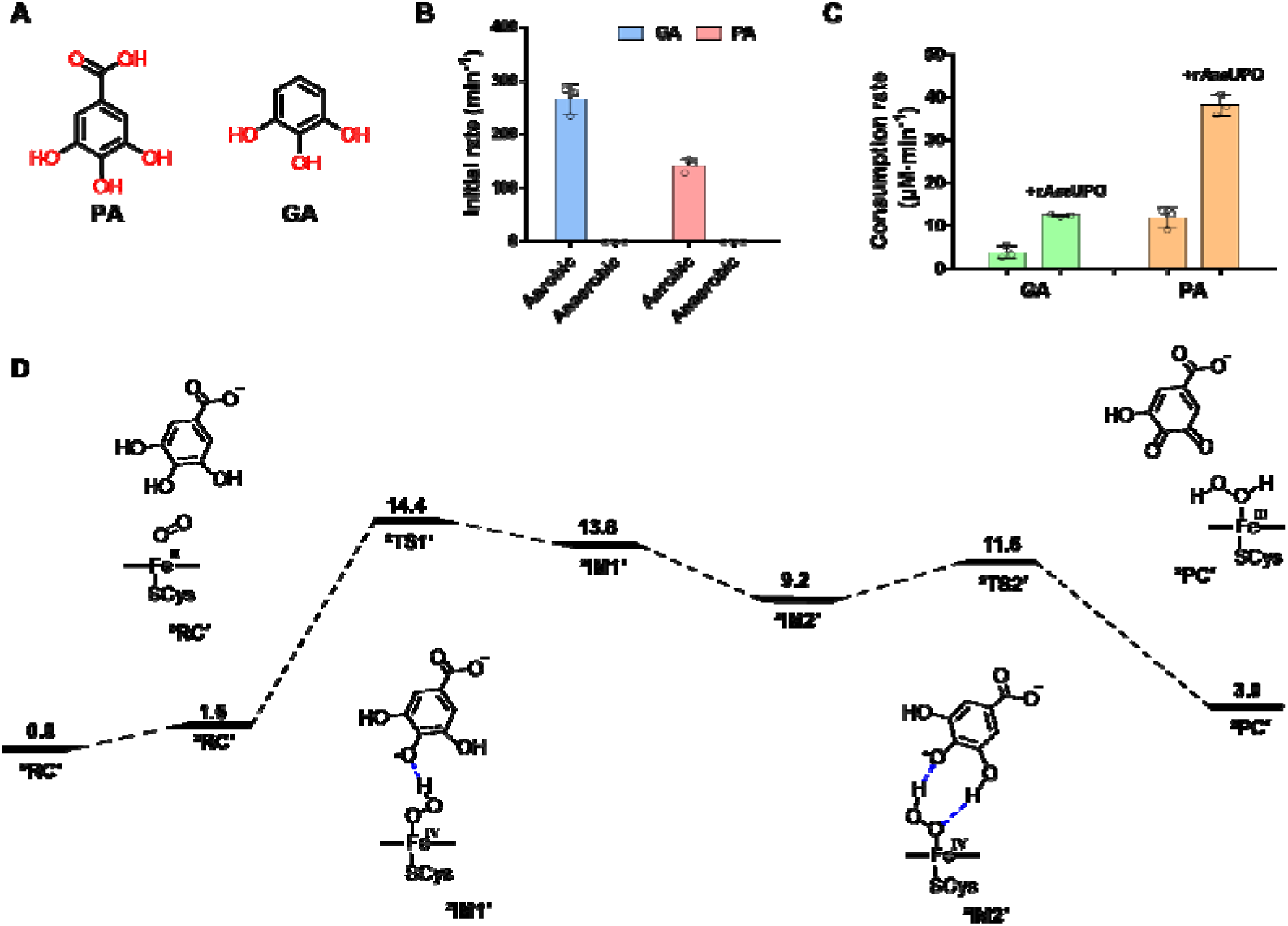
Identification of GA and PA as co-substrates for r*Aae*UPO-catalyzed benzylic hydroxylation of 1a. A. Structural formulas for GA and PA. B. Enzymatic reactions mediated by GA or PA under aerobic and anaerobic conditions with a reaction time of 1 h. Reactions were performed in potassium phosphate buffer (1 mL, 100 mM, pH 8.0, 30 °C) containing r*Aae*UPO (100 nM), 1a (10 mM), 5% (v/v) acetonitrile, GA or PA (10 mM). C. The consumption rate of GA and PA in buffer solution (0.1 M, pH 7.0) with and without r*Aae*UPO addition (800 nM enzyme, 6 mM GA for 60 min reaction, 5 mM PA for 10 min reaction). D. QM(OLYP/def2-TZVP)/MM calculated energetics for the UPO-mediated O_2_ activation in the presence of GA. The calculated reaction energy (ΔE) and energy barriers (ΔE^≠^) are given in kcal mol^-1^. The geometries of species involved in reactions can be found in Supplementary Figure 9. The data shown in B and C are presented as mean value□±□SD (standard deviations) of three biological replicates.

Next, computational studies were conducted to elucidate the mechanism of UPO-mediated O_2_ activation in the presence of polyphenolic reductants, using GA as an example. Similar to the reaction with DHAA, the O_2_ binding is coupled to the H-abstraction from GA, affording the OOH radical species that coordinates with Fe(III) (^2^IM1’ in Figure 5D). Then, one OH group of the GA radical can form the H-bonding interaction with the •OOH radical species, leading to the formation of a more stable intermediate ^2^IM2’. We found that the •OOH radical readily performs hydrogen atom abstraction (HAA) from the adjacent OH group of the phenol radical, resulting in the formation of Fe(III)-H_2_O_2_ intermediate. We assume that the Fe(III)-H_2_O_2_ intermediate may further convert to the active species of Cpd I *via* the O-O heterolysis [32]. It is worth mentioning that excess H_2_O_2_ can be aerobically generated in buffer solution containing GA [33–34], which may rationalize why the use of GA can lead to the activity inhibition for UPO, a phenomenon commonly observed in the H_2_O_2_-dependent actions of peroxidases [31, 35] (**Supplementary Figure 10**). Nevertheless, the experiments showed that the amount of product formed is 1.5-fold higher than the amount of H_2_O_2_ generated in buffer solution (**Supplementary Figure 11**), suggesting that some of the H_2_O_2_ (in the form of Fe(III)-H_2_O_2_) used for catalysis still originates from the O_2_ activation within the enzyme binding pocket.

### Generality of O_2_/reductant-dependent catalytic route in heme peroxygenases

To demonstrate the universality of developed O_2_/reductant-dependent routes in heme-containing peroxygenases, various reductants including AscA, DHA, GA, and PA were investigated in five more representative peroxygenases. These included the “short” type *Mro*UPO (from *Marasmius rotula*)[36] involved hydroxylation of cyclohexane, *Cfu*CPO (from *Caldariomyces fumago*)[37] mediated sulfoxidation of phenyl methyl sulfide, as well as three P450 peroxygenases (P450_SPα_ from *Sphingomonas paucimobilis*[12], P450_BSβ_ from *Bacillus subtilis*[10] and OleT_JE_ from *Jeotgalicoccus sp.* ATCC 8456[1]) catalyzed hydroxylation or decarboxylation of lauric acid. As shown in Figure 6, reductants-fueled UPOs exhibited significantly higher activity compared to H_2_O_2_-dependent reactions (Figures 6A and **6B**). In the case of *Cfu*CPO that showed relatively high H_2_O_2_ tolerance, only DHA and GA displayed improved catalytic performance relative to the H_2_O_2_-dependent process (Figure 6C). However, for P450_SPα_ and P450_BSβ_-catalyzed α or β-hydroxylation of lauric acid, all the reductant-fueled reactions exhibited much higher activity than the H_2_O_2_-dependent process (Figures 6D and **6E**). Notably, GA and PA achieved nearly 100% substrate conversion. While in the decarboxylation reaction catalyzed by P450 OleT_JE_, most reductant-fueled reactions outperformed H_2_O_2_, although GA displayed relatively poor activity (though still better than H_2_O_2_), which requires further investigation (Figure 6F). Thus, we have demonstrated the widespread feasibility of the O_2_/reductant-dependent catalytic route in heme-containing peroxygenase, showcasing excellent catalytic performance. We propose that this O_2_/reductant-dependent route may also be applicable to numerous other heme-containing enzymes.

**Figure 6.**
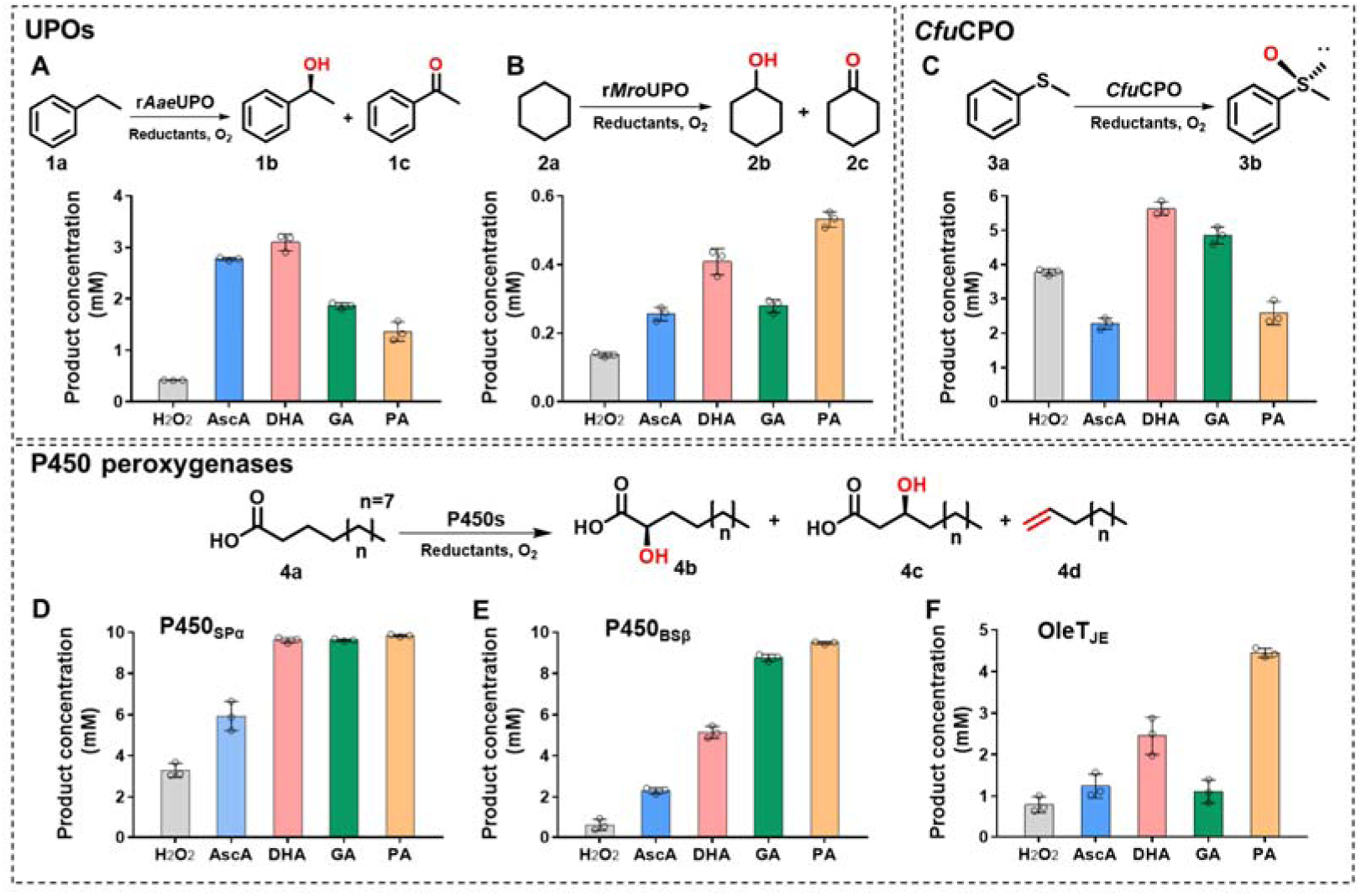
Different UPOs and P450 peroxygenases catalyzed oxidative reactions using different reducing agents. A. The reaction of r*Aae*UPO-catalyzed benzylic hydroxylation of 1a using different reducing agents. B. The reaction of *Mro*UPO-catalyzed hydroxylation of cyclohexane 2a using different reducing agents. C. The reaction of *Cfu*CPO-catalyzed sulfoxidation of phenyl methyl sulfide 3a. D, E and F. The reactions of P450_SPα_, P450_BSβ_ and P450OleT_JE_ catalyzed α or β hydroxylation and decarboxylation of lauric acid 4a. All Reactions were performed in potassium phosphate buffer (1 mL, 100 mM, pH 8.0, 30°C) containing different enzymes (0.1 μM r*Aae*UPO; 0.2 μM *Mro*UPO; 0.15 μM *Cfu*CPO; 2 μM P450_SPα_; 2 μM P450_BSβ_; 2 μM OleT_JE_), substrates (each 10 mM), reducing agents (each 20 mM) and 5% (v/v) cosolvents (acetonitrile for 1a; acetone for 2a; methanol for 3a; DMSO for 4a) for 20 h. The reaction with addition of 20 mM H_2_O_2_ was employed as control. The data shown in A, B, C, D, E and F are presented as mean value□±□SD (standard deviations) of three biological replicates.

### Further comparison of O_2_/reductant-dependent route with H_2_O_2_-dependent process

To further demonstrate the superiority of the developed O_2_/reductant-dependent route over the conventional H_2_O_2_-dependent process, spectroscopic analysis was conducted on the enzyme r*Aae*UPO treated with varying concentrations of H_2_O_2_ or AscA. When r*Aae*UPO was treated with exogenous H_2_O_2_, UV-vis spectrum showed a gradual decrease in absorption intensity of Soret band (ferric resting state) and the emergence of gas bubbles (**Supplementary Figure 12**). This is consistent with the heme-bleaching and the generation of O_2_ caused by excessive H_2_O_2_ in a previous study [38] (see proposed heme-bleaching mechanism in **Supplementary Figure 13**). By contrast, the UPO/DHAA/O_2_ system affords a single Fe(III)-H_2_O_2_ intermediate that is well bound in the active site, in which the binding and reaction of an additional H_2_O_2_ molecule is avoided. As such, the reaction is well controlled toward the Cpd I generation and the subsequent substrate oxidation, thus effectively preventing the irreversible heme damage and related enzyme inactivation.

Furthermore, due to the strong oxidizing nature of H_2_O_2_, the catalytic system with exogenous H_2_O_2_ may not be suitable for substrates susceptible to oxidations by H_2_O_2_. To demonstrate this, quercetin (**5a**), a substrate prone to oxidation by H_2_O_2_, has been tested in both r*Aae*UPO/O_2_/reductant and H_2_O_2_-dependent systems (Figure 7A). In the O_2_/reductant dependent system with either AscA or DHA, **5a** can be selectively hydroxylated at C6-position to produce quercetagetin (**5b**) (Figures 7B **and 7C, Supplementary Figure 14**). By contrast, no C6-hyrdoxylated product can be formed when exogenous H_2_O_2_ was used, which can result in the degradation **5a** to form unknown by products (Figure 7B). Similar things occurred for reductants (PA or GA) that generate excess H_2_O_2_ in buffer solution. These findings provide more convincing support that the O_2_/(AscA/DHA)-dependent catalytic route eliminates the exogenous H_2_O_2_ usage, which is particularly suitable for substrates susceptible to oxidations by H_2_O_2_. It is worth noting that the product quercetagetin (**5b**) possesses diverse biological activities, such as antioxidant and anti-inflammatory properties, making it a valuable additive in the livestock and poultry breeding industry [39–40]. Therefore, our study once again demonstrates the superiority of the O_2_/reductant-dependent catalytic route over the conventional H_2_O_2_-dependent process.

**Figure 7.**
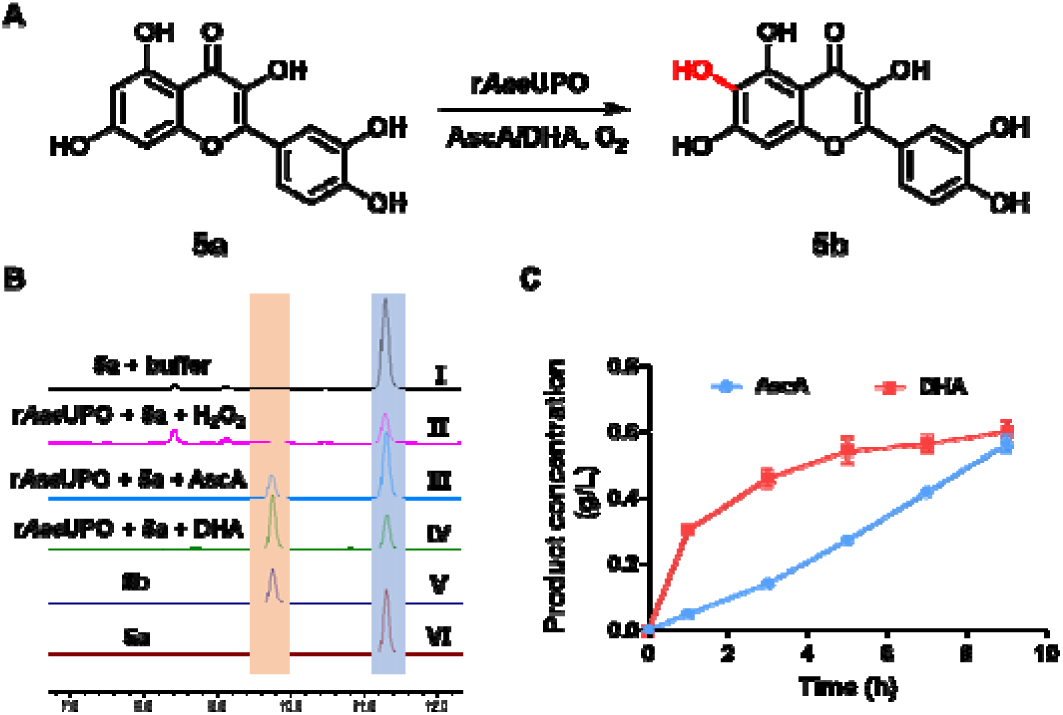
Comparison of r*Aae*UPO-catalyzed biotransformation of quercetin **5a** to **5b** between O_2_/reductant-dependent route and H_2_O_2_-dependent process. **A**. Reaction scheme of r*Aae*UPO-catalyzed selective C6-hydroxylation of quercetin **5a** to **5b**. **B**. Representative HPLC chromatograms of r*Aae*UPO-catalyzed hydroxylation of **5a** in the presence of AscA, DHA or H_2_O_2_. (□) Potassium phosphate buffer (100 mM, pH 8.0) + **5a**, (□) r*Aae*UPO + **5a** + H_2_O_2_, (□) r*Aae*UPO + **5a** + AscA, (□) r*Aae*UPO + **5a** + DHA, (□) standards of quercetagetin (**5b**) and (□) quercetin (**5a**). **C**. Time course of r*Aae*UPO-catalzyed hydroxylation of **5a** with AscA or DHA as a reductant, respectively. Reactions were performed in potassium phosphate buffer (100 mM, pH 8.0, 37°C) containing r*Aae*UPO (1 μM), AscA or DHA (50 mM), **5a** (1.5 g/L), 5% (m/v) 2-hydroxypropyl-β-cyclodextrin, 10% acetone.

### Scale-up reactions for product preparation

To demonstrate the potential for industrial applications, we conducted a scale-up production of the high value-added products using the reductant-fueled biocatalytic system. Our focus was on two specific reactions: the r*Aae*UPO catalyzed benzylic hydroxylation of **1a** to produce (*R*)-1-phenylethanol (**1b**) and the P450_SPα_-catalyzed α-hydroxylation of **4a** to produce α-OH lauric acid (**4b**). We chose AscA as the reductant due to its lower cost and greater availability compared to DHA (**Supplementary Figure 15**). It is important to note that both products have a wide range of applications in industries such as food, cosmetics, and pharmaceuticals [41, 42].

To conduct the r*Aae*UPO catalyzed hydroxylation of **1a**, we performed the reaction in a 5 L fermenter containing 1 L reaction mixture consisting of 200 mM **1a**, 40 g AscA, and 0.25 μM r*Aae*UPO. After reaction, 9.9 g/L of enantiomerically pure (*R*)-1-phenylethanol (**1b)** and 1.8 g/L of acetophenone (**1c**) were obtained, which corresponds to 395,600 catalytic turnovers. These results represent the highest reported product titer for (*R*)-1-phenylethanol production, surpassing the productivity achieved with adjusted H_2_O_2_ feed rate (5 g/L) [43]. In addition, after purification, 7.72 g of **1b** with 31.6% isolated yield and 1.33 g of **1c** were obtained with purities exceeding 98%. It is worth noting that the unreacted substrate can be easily separated and recycled for next batch. While for the P450_SPα_ catalyzed α-hydroxylation of **4a**, we also conducted the reaction in a 5 L fermenter containing 1 L reaction mixture consisting of 10 g **4a** (with an additional 2 g added at 72 h), 30 g AscA, and 1.5 μM P450_SPα_. Remarkably, the substrate conversion reached >99%, resulting in 11.82 g of α-OH lauric acid (**4b**) with an isolated yield of 91%. These results achieved a new record for both the product titer of α-hydroxy lauric acid and the catalytic turnovers of P450_SPα_ (TTN 43,760), outperforming previous reported values by more than six times [44]. It should be noted that the reductant AscA is cheap and readily available bulk chemical, with the price that is two orders of magnitude lower than that of products synthesized (**1b** or **4b**). These results highlight the significant synthetic potential of the reductant-fueled peroxygenase technology, particularly for the synthesis of high value-added compounds.

**Figure 8.**
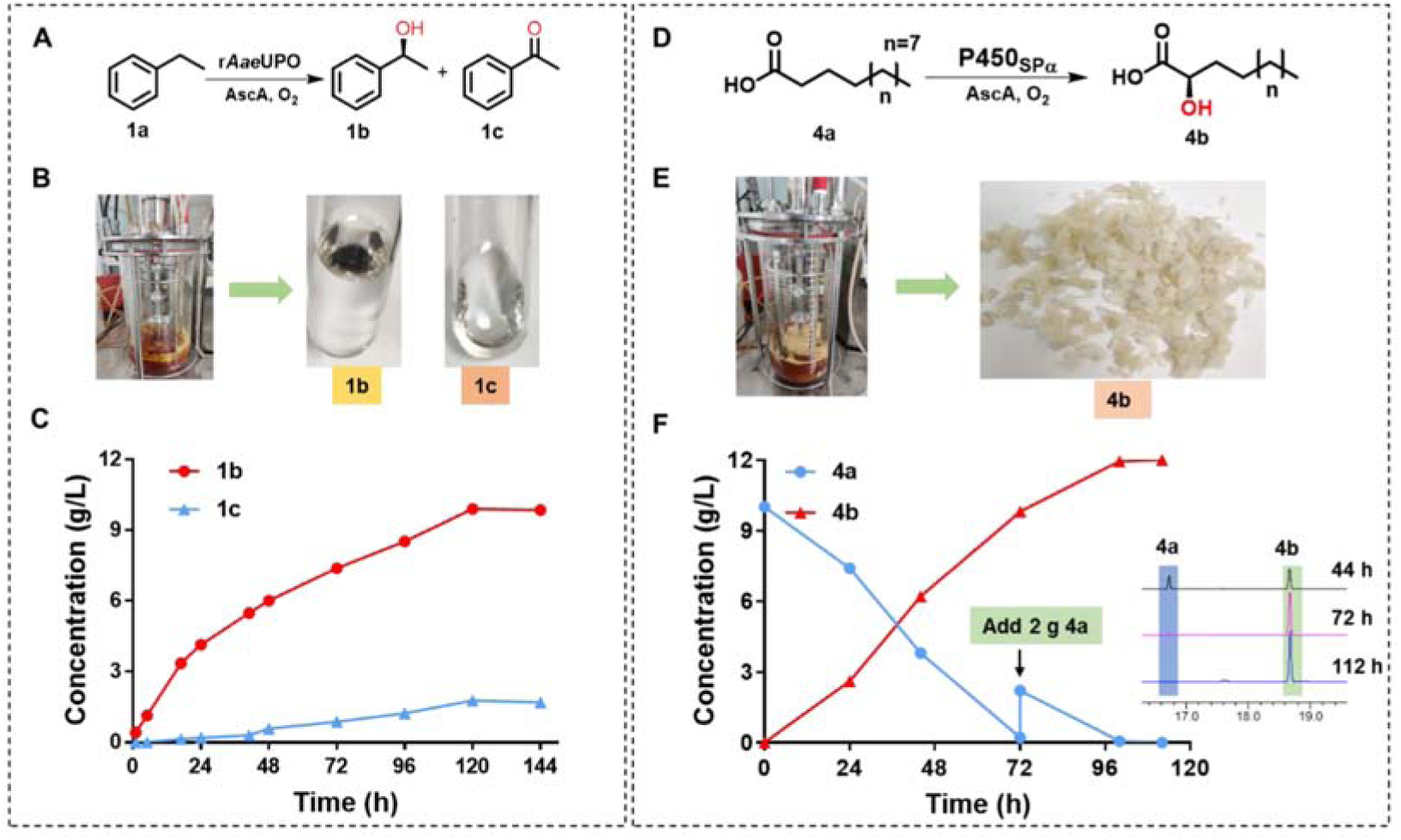
Scale-up reactions for preparations of (*R*)-1-phenylethanol (**1b**) and α-OH lauric acid (**4b**) with the reductant-fueled biocatalytic system. Left (**A**, **B**, **C**): Preparative-scale reaction for the production of **1b** by r*Aae*UPO-catalyzed hydroxylation of **1a** using AscA as the reductant. Reaction was performed in 1 L potassium phosphate buffer (100 mM, pH 8.0, 30 °C) containing r*Aae*UPO (0.25 μM), **1a** (200 mM), AscA (40 g), 10% (m/v) HP-β-CD with pH maintained at 7.0 ∼ 8.0. Right (**D**, **E**, **F**): Preparative-scale reaction for the production of **4b** by P450_SPα_-catalyzed hydroxylation of **4a** using AscA as the reductant. Reaction was performed in 1 L potassium phosphate buffer (100 mM, pH 8.0, 37 °C) containing P450_SPα_ (1.5 μM, OD_600_ is about 40), substrate (12 g), AscA (30 g), 30% (m/v) HP-β-CD and 10% DMSO with pH value maintained at 7.0 ∼ 8.0.

### Conclusions

In summary, we reported a highly efficient but previously undisclosed O_2_/reductant-dependent catalytic route in heme-containing peroxygenases. In this biocatalytic process, reductants such as DHA or AscA present in fresh plant tissue, or polyphenolic compounds like GA or PA abundant in lignified tissue, can serve as co-substrates to initiate oxidative reactions. For reactions of UPO with either AscA or DHA, experimental and computational studies revealed that DHAA (the hydrated form of DHA) is the actual co-substrate responsible for activating oxygen to generate oxyferryl heme (compound I, Cpd I) as the oxygenation species. In this new route, the controlled Fe(III)-H_2_O_2_ intermediate is formed in the binding pocket of UPO, thereby eliminating the excessive H_2_O_2_ production and thus preventing the heme destruction and subsequent enzyme inactivation. Unlike AscA or DHA, the use of reducing agents GA or PA led to the excess H_2_O_2_ production of buffer solution (outside the enzyme pocket), resulting in the inhibition of enzyme activity to some extent. Interestingly, we found that the newly discovered O_2_/reductant-dependent catalytic route is also applicable to other types of heme-containing peroxygenases (including short type UPO, CPO and CYP152). Furthermore, our findings also demonstrate that the developed system utilizing AscA/DHA as reductants system can achieve the efficient and selective oxidation of substrates that are susceptible to oxidation by H_2_O_2_, as exemplified in the biotransformation of quercetin. Collectively, these results also suggest that heme peroxygenases can function as monooxygenases, depending on the availability of O_2_ and external electron donors (or reductants) in the surrounding environment. We anticipate that the O_2_-dependent route will have far-reaching implications in future biocatalysis, as well as in the physiological metabolism of fatty acids, lignin degradation, and plant infection by pathogenic oomycetes.

Finally, to demonstrate the practical feasibility of the developed biocatalytic system, the scale-up reactions were conducted for r*Aae*UPO catalyzed hydroxylation of ethylbenzene and P450_SPα_ mediated α-hydroxylation of lauric acid, respectively. Remarkably, we achieved the highest productivity record for the production of (*R*)-1-phenylethanol and α-OH lauric acid, both of which are widely sought after in the food, cosmetic, and pharmaceutical industries. Moving forward, our future endeavors will focus on the practical applications of this advanced technology to produce more high-value products on an industrial scale. The potential application of advanced mutagenesis methods in directed evolution to enhance and reverse stereoselectivity[45] would further strengthen the capabilities of our newly developed oxidative system.

## Methods

### Construction of recombinant expression cells

DNA fragments of enzyme genes and the linear plasmid backbone were amplified by PCR using primers with 15 to 20 bp homologous arms to facilitate subsequent recombination. Full length of enzyme genes was assembled *via* overlap PCR and cloned into linear vector with the help of T5 exonuclease for generating 15 bp or 20 bp sticky ends to promote recombination efficiency. The reaction mixtures (5 μL), consisting of linear vector, enzyme genes, buffer 4.0 (New England Biolabs) and T5 exonuclease, were incubated in ice-water for 5 min. Subsequently, 50 μL competent cells (*E. coli* DH5α) were quickly added for transformation, followed by plating on LB agar or low salt LB agar containing appropriate antibiotics. The resulting transformants were picked and their DNA was sequenced for confirmation. For protein expression and whole-cell biocatalyst preparation, plasmids containing P450 peroxygenase genes were transformed into *E. coli* BL21 cells. On the other hand, plasmids containing unspecific peroxygenase genes were linearized with the restriction enzyme *Pme* I at 37 L for 1 h and then transformed into *P. pastoris* X33 cells for protein expression. The detailed information about the synthetic gene sequences, plasmids, and corresponding expression hosts are listed in **Supplementary Tables 2-3.**

Typical procedure for *E. coli* protein expression and preparation of whole-cell catalysts. Constructed *E. coli* cells were inoculated into 4 mL LB medium containing antibiotics (50 μg mL^−1^ kanamycin), and cultured at 37 °C, 220 rpm for 6 h. Following this, precultures (4 mL) were transferred into 400 mL TB medium with appropriate antibiotics in 1 L shaking flasks and cultured at 37 °C, 220 rpm for 2 ∼ 3 h until the OD_600_ reached 0.6 ∼ 0.8. At this point, IPTG was added to give a final concentration of 0.2 mM. The temperature was then lowered to 16 °C and maintained for 18∼20 h to allow for protein expression. The cells were harvested by centrifugation at 5000 × *g*, 4 °C for 10 min, washed with 100 mM potassium phosphate buffer (pH 8.0) and used as whole-cell biocatalysts in the subsequent reactions.

### Typical procedure for *P. pastoris* protein expression in shaking flask

To initiate the growth of constructed *P. pastoris* cells, they were inoculated into 100 mL shaking flasks containing 50 mL of BMGY medium and incubated at 28 °C, 250 rpm for 48 h. Once the optical density at 600 nm (OD_600_) reached approximately 20, the supernatant was centrifuged, and the cells were collected. These cells were then resuspended by adding 25 mL of BMMY medium and induced at 220 rpm at 25 °C in 100 mL shaking flasks. To induce protein expression, 1% (v/v) methanol was added every 24 hours, and this process was repeated for 4 to 5 days. Finally, the supernatant was collected for further use.

### Typical procedure for *P. pastoris* high density culture in 5 L fermenter

The strains exhibiting the highest activity in shaking flasks were chosen for large-scale cultivation using a 5 L fermenter, which was initially filled with 2.5 L of basal salts medium. After sterilization, 4.35 mL/L of PTM1 trace salts and 1 mL of antifoam were added to the medium. The pH was adjusted to 5.5 using a 25% ammonium hydroxide solution and maintained at this level throughout the entire process. Fermentation was initiated by inoculating 0.2 L of *P. pastoris* preculture, which had been grown on YPD medium in multiple 1 L baffled shaking flasks at 220 rpm and 28 °C for 36 h. Following the *Pichia* Fermentation Process Guidelines provided by Invitrogen, the batch was carried out at 28 °C and 500 rpm. When all the glycerol was consumed, a 50% (*w/v*) glycerol feed containing 12 mL/L of PTM1 trace salts was introduced to maintain a dissolved oxygen (DO) concentration above 30%. Once the OD_600_ reached above 200, the glycerol supplementation was halted for a 30-minute period to induce starvation in *P. pastoris*. Subsequently, the methanol feed was initiated upon observing a spike in the DO concentration. The rate of methanol supplementation was carefully controlled to maintain the DO at approximately 20%. At this stage, the temperature was adjusted to 25 °C. Regular sampling was performed to monitor wet biomass, OD_600_, and NBD activity. After approximately 5 days of methanol induction, the fermentation process was terminated, and the supernatant of the fermentation broth was collected for further purification.

### Typical procedure for protein purification

The *E. coli* crude cell extract was obtained by disrupting the cells using a high-pressure homogenizer. The resulting cell lysates were then centrifuged at 4 °C for 30 minutes at 17,000 × g to remove any cell debris. The filtrate was obtained by passing the supernatant through a 0.22 μM filter. Similarly, the *P. pastoris* supernatant was also centrifuged and filtered to obtain a filtrate. The obtained filtrate was then applied to a Ni-NTA column using a fast protein liquid chromatography (FPLC) system from GE Healthcare. The column was equilibrated with potassium phosphate buffer A (100 mM, pH 8.0 with 0.5 M NaCl), and the target protein was eluted using a gradient of 10 to 500 mM imidazole. The eluent containing the protein of interest was collected and concentrated using Amicon filters with a 10 kDa cut-off. It was then desalted using a HiTrap desalting column and potassium phosphate buffer (100 mM, pH 8.0). All purified proteins were stored at -80 °C after being quick-frozen with liquid nitrogen for further assays. The purification of OleT_JE_ was performed following previously reported methods [46].

### Derivatization

After reaction, the mixtures were acidified to below pH 2.0 using 4 M HCl and extracted with ethyl acetate. The organic phase was then dried over anhydrous Na_2_SO_4_. To remove Na_2_SO_4_, the obtained mixtures (products in ethyl acetate) were centrifuged at 13,680 × g for 10 minutes, and 300 µL of the resulting supernatant solutions were transferred to fresh 1.5 mL tubes. After evaporation of the ethyl acetate, the resulting solid was dissolved in 30 µL of N-methyl-N-(trimethylsilyl) trifluoroacetamide and 60 µL of pyridine. The derivatization reactions were carried out at 65°C for 1 hour, and the resulting mixtures were used for GC analysis with an SH-Rtx-5 column to analyze lauric acid and its hydroxylation products.

### Scale-up reaction for converting 1a to 1b and 1c

The biotransformation of **1a** to **1b** and **1c** was upscaled using r*Aae*UPO in 5-L fermenters (T&J Bio-engineering, Shanghai, China). First, 100 g HP-β-CD was dissolved in 1 L potassium phosphate buffer (0.1 M, pH 8.0), followed by addition of 200 mM **1a**, 20 g AscA and 0.25 μM r*Aae*UPO. Subsequently, an additional 5 g AscA was added after 48 h reaction and then every 24 h thereafter. Reaction was carried out at 30L, 500 rpm in 5-L fermenter, and pH was maintained around 7.0∼8.0 by the addition of KOH solution. The reaction was monitored by gas chromatography analysis. After reaction was completed, ethyl acetate (1 L) was added for product extraction. The resulting mixture was then subjected to centrifugation, and the organic phase was evaporated to obtained the crude product that was further purified by SiO_2_ column chromatography with ethyl acetate and petroleum ether (1:3) as mobile phase. The purified products were finally subjected to Nuclear magnetic resonance (NMR) analysis for structure identification.

### Scale-up reaction for converting 4a to 4b

The biotransformation of **4a** to **4b** was upscaled in 1-L fermenters (T&J Bio-engineering, Shanghai, China) using recombinant *E. coli* expressing P450_SPα_. The recombinant *E. coli* expressing P450_SPα_ was cultured as described above, the harvested whole-cell catalyst was re-suspended in 900 mL potassium phosphate buffer (0.1 M, pH 8.0) to reach an OD_600_ about 40 (*ca*. 1.5 μM concentration of P450_SPα_). Subsequently, 300 g HP-β-CD was dissolved in 900 mL cell resuspension in a bioreactor, followed by addition of 10 g **4a** (dissolved in 100 mL DMSO) and 10 g AscA. Afterwards, 5 g AscA was added every 24 h. Reaction was carried out at 37L, 500 rpm in 5-L fermenter, and pH was maintained around 7.0∼8.0 by the addition of KOH solution. The reaction was monitored by gas chromatography analysis. An additional 2 g **4a** was supplemented when the initial amount of **4a** was completely converted at 72 h. After reaction, the extraction, separation and identification of the products were conducted according to the procedure described above.

### GC and HPLC analysis

The GC measurements were conducted using a Shimadzu GC-2030/FID equipped with various columns. The reactions were stopped at intervals by adding ethyl acetate containing dodecane (2.5 mM) as an internal standard. After extraction and centrifugation, the organic phase was dried using anhydrous sodium sulfate and analyzed *via* gas chromatography with CP-Chirasil-DEX CB or SH-Rtx-WAX column. All concentrations reported were based on calibration curves obtained from authentic standards, and GC chromatograms were shown in **Supplementary Figures** 16-19. The HPLC measurements were performed on a Shimadzu LC-2030C system with a Shimadzu SPD-M20A Photo Diode Array detector. The analytical detection of AscA, GA, and PA required the addition of 4 M HCl to lower the pH below 2 before detection. HPLC chromatograms were shown in **Supplementary Figures 20-23**. Specific analysis methods were described in **Supplementary Table 4**.

### NMR analysis

NMR chromatograms were shown in **Supplementary Figures 24-26**. **1b:** ^1^H NMR (400 MHz, CH_3_OH-*d*_4_): δ 7.37 (m, 1H), 7.35 (m, 1H), 7.32 (m, 1H), 7.30 (m, 1H), 7.22 (tt, *J* = 7.2, 1.5 Hz, 1H), 4.82 (q, *J* = 6.5 Hz, 1H), 1.4 (d, *J* = 6.5 Hz, 3H); ^13^C NMR (100 MHz, CH_3_OH-*d*_4_): 147.5, 129.3 × 2, 128.1, 126.5 × 2, 70.8, 25.6. **1c:** ^1^H NMR (400 MHz, DMSO-*d*_6_): δ 7.97 (m, 1H), 7.95 (m, 1H), 7.63 (tt, *J* = 7.4, 1.4 Hz, 1H), 7.53 (m, 1H), 7.51 (m, 1H), 2.57 (s, 3H); ^13^C NMR (100 MHz, DMSO-*d*_6_): 198.0, 137.0, 133.3, 128.8 × 2, 128.3 × 2, 26.8. **4b:** ^1^H NMR (400 MHz, CH_3_OH-*d*_4_): δ 4.12 (ddd, *J* = 12.3, 7.8, 4.5, Hz, 1H), 1.69–1.80 (m, 1H), 1.58–1.69 (m, 1H), 1.22–1.37 (overlap, 14H), 0.90 (t, *J* = 7.0 Hz, 3H); ^13^C NMR (100 MHz, CH_3_OH-*d*_4_): 178.0, 71.4, 35.4, 33.1, 30.7 × 2, 30.6, 30.5 × 2, 26.1, 23.7, 14.5.

### System setup and MD simulations

The initial structure was obtained from the crystal structure of r*Aae*UPO (PDB: 5OUX)[47], *Mro*UPO (PDB: 5FUJ), *Cfu*CPO (PDB: 1CPO) [48], OleT_JE_ (PDB: 4L54) [49], P450_BSβ_ (PDB: 2ZQX) [50], and P450_SPα_ (PDB: 3VM4) [51]. The protonation states of titratable residues (His, Glu, Asp) were calculated using PROPKA [52]. All dockings were performed using the AutoDock Vina tool [53–54] and Partial atomic charges of the ligands were set using the RESP approach at the B3LYP/def2-TZVP theory level [55]. The protein was treated by Amber ff14SB force field and general AMBER force field (GAFF) [56] and the ferric resting state were parametrized using the “MCPB.py” modeling tool [57–58]. Subsequently, sodium ions were added to the surface of the protein and the entire system was solvated in a rectangular box with TIP3P waters (minimum distance was 20 Å from the protein surface). After the setup of the system, MD simulations were performed according to the following steps. (1) Each system was totally minimized by the combined steepest descent and conjugate gradient methods. (2) Each system was annealed from 0 to 300 K for 50 ps with the NVT ensemble. (3) These systems were subjected to equilibration of 1 ns in an NPT ensemble at a target temperature of 300 K and a target pressure of 1 bar. The target temperature and target pressure were maintained with the Langevin thermostat [59] and the Berendsen barostat [60]. (4) The enzyme complexes were equilibrated for 4 ns under the NPT ensemble. (5) A productive MD for 50 ns was performed in an NPT ensemble. During the simulations, the covalent bonds involving hydrogen atoms were constrained with the SHAKE method [61] and the long-range electrostatic interactions were treated using the Particle Mesh Ewald (PME) method [62]. Finally, the binding free energies of reductants were calculated from snapshots of the last 4 ns MD trajectory using the MM/GBSA [63], with an interval of 10 ps. All MD simulations were performed using Amber 18 software package [64]. More details can be found in SI.

### QM/MM calculation

Based on the MD simulations and clustering analysis of complex (**Supplementary Figure 27-28**, the representative conformations were selected in simulation trajectory. The QM region included the substrate, the Heme-Fe(III) complex the axial ligand of Cys36, oxygen, residue R189 and residue E196 (**Supplementary Figure 29**). In the reaction with DHAA, since E196 acted as a proton acceptor to accept the proton of DHAA and then transferred the proton to Fe(III)-OOH, we selected the second most populated conformation with a close distance between E196 and Fe atom for QM/MM calculation (**Supplementary Figure 27**). For the system with the substrate GA, the active site structure of the three most populated structures were quite similar (**Supplementary Figure 28**). The QM/MM calculations were conducted using ChemShell [65–67], which applied ORCA [68–72] for QM region and DL_POLY [73] for MM region. Electronic embedding scheme [74] was employed to account the polarizing effect of the enzyme environment in the QM region, while the hydrogen link atoms with the charge-shift model was used to deal with the QM/MM boundary. The QM part of the system was treated by the pure functional OLYP [75–76] with the all-electron basis set of def2-SVP, whereas the MM part was modeled at the classical level using the same parameters as in the MD simulations. Energies were further corrected with the large all-electron basis set def2-TZVP. The transition states were determined as the highest energy structure from the adiabatic scan, which was further optimized using the P-RFO optimizer [77] without any restraint.

### Density functional theory (DFT) calculation

DFT calculations were carried out using Gaussian 16 software [78]. The geometry optimizations and frequency calculations were carried out at the B3LYP [76, 79–81] level with the SMD continuum solvation model [82]. And the dispersion corrections computed with Grimme’s D3 method were included in DFT calculations [83–84]. The energies were further refined with the larger basis set def2-TZVP for all atoms.

## Reporting summary

Further information on research design is available in the Nature Portfolio Reporting Summary linked to this article.

## Data availability

The data that support the findings of this study are available within the main text and its Supplementary Information file. Data are also available from the corresponding author upon request.

## Acknowledgments

This study was supported by the National Key Research and Development Program of China (2019YFA09005000), by the National Natural Science Foundation of China (No. 32371552, 22122305 and 21702052) and Research Program of State Key Laboratory of Biocatalysis and Enzyme Engineering. We also thank Dr. Xuexia Xu and Dr. Xin Liu for helpful discussions and supports.

## Author Contributions

A.L. and B. W. conceived and supervised the project, D. D., Z. J., L. K., L. L., X. Z. and Y. Q. performed the experiments and analyzed the data; A.L., B. W., D. D. and Z. J. wrote the manuscript; all authors checked and modified the manuscript.

## Competing Interests

The authors declare no competing interests.

## Additional information

Supplementary information is available in the online version of the paper. Correspondence and requests for materials should be addressed to A.L. or B. W. Peer review information: XXX thanks the anonymous reviewer(s) for their contribution to the peer review of this work. Peer reviewer reports are available.

## Notes

### Competing Interest Statement

The authors have declared no competing interest.

## References

1. Sigmund, M. C. & Poelarends, G. J. Current state and future perspectives of engineered and artificial peroxygenases for the oxyfunctionalization of organic molecules. Nat. Catal. 3, 690–702 (2020).

2. Hobisch, M. Holtmann, D. Gomez de Santos, P. Alcalde, M. Hollmann, F. & Kara, S. Recent developments in the use of peroxygenases-Exploring their high potential in selective oxyfunctionalisations. Biotechnol. Adv. 51, 107615 (2021).

3. Kumari, R. Singh, A. & Yadav, A. N. Fungal enzymes: Degradation and detoxification of organic and inorganic pollutants. Recent Trends in Mycological Research 2, 99–125 (2021).

4. Hanano, A., et al. Plant seed peroxygenase is an original heme-oxygenase with an EF-hand calcium binding motif. J. Biol. Chem. 281, 33140–33151 (2006).

5. Fuchs, C. & Schwab, W. Epoxidation, hydroxylation and aromatization is catalyzed by a peroxygenase from *Solanum lycopersicum*. *J. Mol. Catal.*, B Enzym. 96, 52–60 (2013).

6. Podust, L. M. & Sherman, D. H. Diversity of P450 enzymes in the biosynthesis of natural products. Nat. Prod. Rep. 29, 1251–1266 (2012).

7. Aratani, Y. Myeloperoxidase: Its role for host defense, inflammation, and neutrophil function. Arch Biochem Biophys 640, 47–52 (2018).

8. Kim, J. Hollmann, F. & Park, C.B. Lignin as a multifunctional photocatalyst for solar powered biocatalytic oxyfunctionalization of C-H bonds. Nat. Synth. 1, 217–226 (2022).

9. Grogan, G. Hemoprotein catalyzed oxygenations: P450s, UPOs, and progress toward scalable reactions. JACS Au 1, 1312–1329 (2021).

10. Zhang, K. et al. Biocatalytic enantioselective βLhydroxylation of unactivated C−H bonds in aliphatic carboxylic acids. Angew. Chem. Int. Ed. 61, e202204290 (2022).

11. Pickl, M., et al. Mechanistic studies of fatty acid activation by CYP152 peroxygenases reveal unexpected desaturase activity. ACS Catal. 9, 565–577 (2018).

12. Ramanan, R. Dubey, K. D. Wang, B. Mandal, D. & Shaik, S. Emergence of function in P450-proteins: A combined quantum mechanical/molecular mechanical and molecular dynamics study of the reactive species in the H_2_O_2_-dependent cytochrome P450_SPα_ and its regio- and enantioselective hydroxylation of fatty acids. J. Am. Chem. Soc. 138, 6786–6797 (2016).

13. Jiang, Y., et al. Unexpected reactions of α, β-unsaturated fatty acids provide insight into the mechanisms of CYP152 peroxygenases. Angew. Chem. Int. Ed. 60, 24694–24701 (2021).

14. Urlacher, V. B. & Girhard, M. Cytochrome P450 monooxygenases in biotechnology and synthetic biology. Trends Biotechnol. 37, 882–897 (2019).

15. Li, S. Du, L. & Bernhardt, R. Redox partners: function modulators of bacterial P450 enzymes. Trends Microbiol. 28, 445–454 (2020).

16. Burek, B. O. Bormann, S. Hollmann, F. Bloh, J. Z. & Holtmann, D. Hydrogen peroxide driven biocatalysis. Green Chem. 21, 3232–3249 (2019).

17. Kracher, D. et al. Extracellular electron transfer systems fuel cellulose oxidative degradation. Science 352, 1098–1101 (2016).

18. Ullrich, R. Nuske, J. Scheibner, K. Spantzel, J. & Hofrichter, M. Novel haloperoxidase from the agaric basidiomycete *Agrocybe aegerita* oxidizes aryl alcohols and aldehydes. Appl. Environ. Microbiol. 70, 4575–4581 (2004).

19. Molina-Espeja, P. Garcia-Ruiz, E. Gonzalez-Perez, D. Ullrich, R. Hofrichter, M. & Alcalde, M. Directed evolution of unspecific peroxygenase from *Agrocybe aegerita*. Appl. Environ. Microbiol. 80, 3496–3507 (2014).

20. Njus, D. Kelley, P. M. Tu, Y. J. & Schlegel, H. B. Ascorbic acid: The chemistry underlying its antioxidant properties. Free Radic. Biol. Med. 159, 37–43 (2020).

21. Deutsch, J. C. Dehydroascorbic acid. J. Chromatogra. 881, 299–307 (2000).

22. ChebroluK. K. Jayaprakasha, G. K. Yoo, K. S. Jifon, J. L. & Patil, B. S. An improved sample preparation method for quantification of ascorbic acid and dehydroascorbic acid by HPLC. Lwt-food Sci. Technol. 47, 443–449 (2012).

23. Stepnov, A. A. Christensen, I. A. Forsberg, Z. Aachmann, F. L. Courtade, G. & Eijsink, V. G. H. The impact of reductants on the catalytic efficiency of a lytic polysaccharide monooxygenase and the special role of dehydroascorbic acid. FEBS Lett. 596, 53–70 (2021).

24. Yu, H.S. Zhang, W. Verma, P. He, X. & Truhlar, D. G. Nonseparable exchange–correlation functional for molecules, including homogeneous catalysis involving transition metals. Phys. Chem. Chem. Phys. 17, 12146–12160 (2015).

25. Zheng, J., Wang, D., & Thiel, W. QM/MM study of mechanisms for compound formation in the catalytic cycle of cytochrome P450cam. J. Am. Chem. Soc., 128, 13204–13215 (2006).

26. Derat, E., Shaik, S., Rovira, C., Vidossich, P., & Alfonso-Prieto, M. The effect of a water molecule on the mechanism of formation of compound 0 in horseradish peroxidase. J. Am. Chem. Soc., 129, 6346–6347 (2007).

27. Vidossich, P., Fiorin, G., Alfonso-Prieto, M., Derat, E., Shaik, S., & Rovira, C. On the role of water in peroxidase catalysis: a theoretical investigation of HRP Compound I formation. J. Phys. Chem. B, 114, 5161–5169 (2010).

28. Isaksen, et al. A C4-oxidizing lytic polysaccharide monooxygenase cleaving both cellulose and cello-oligosaccharides. J. Biol. Chem., 289, 2632–2642 (2014).

29. Kittl, R., Kracher, D., Burgstaller, D., Haltrich, D., & Ludwig, R. Production of four *Neurospora crassa* lytic polysaccharide monooxygenases in *Pichia pastoris* monitored by a fluorometric assay. Biotechnol. Biofuels, 5, 1–14 (2012).

30. Stepnov A. A, Christensen I. A, Forsberg Z., Aachmann F. L, Courtade G., & Eijsink V. G H. The impact of reductants on the catalytic efficiency of a lytic polysaccharide monooxygenase and the special role of dehydroascorbic acid. FEBS Lett., 596, 53–70 (2022).

31. Bissaro, B., et al. Oxidative cleavage of polysaccharides by monocopper enzymes depends on H_2_O_2_. Nat. Chem. Biol. 13, 1123–1128 (2017).

32. Wang, B. Zhang, X. Fang, W. Rovira, C. & Shaik, S. How do metalloproteins tame the fenton reaction and utilize •OH radicals in constructive manners? Acc. Chem. Res. 55, 2280–2290 (2022).

33. Tulyathan, V. Boulton, R. B. & Singleton, V. L. Oxygen uptake by gallic acid as a model for similar reactions in wines. J. Agric. Food Chem. 37, 844–849 (1989).

34. Grzesik, M. Bartosz, G. Stefaniuk, I. Pichla, M. Namiesnik, J. & Sadowska-Bartosz, I. Dietary antioxidants as a source of hydrogen peroxide. Food Chem. 278, 692–699 (2019).

35. Valderrama, B. Ayala, M. & Vazquez-Duhalt, R. Suicide Inactivation of Peroxidases and the Challenge of Engineering More Robust Enzymes. Chem. Biol. 9, 555–565 (2002).

36. Carro, J. et al. Modulating fatty acid epoxidation vs hydroxylation in a fungal peroxygenase. ACS Catal. 9, 6234–6242 (2019).

37. Getrey, L. Krieg, T. Hollmann, F. Schrader, J. & Holtmann, D. Enzymatic halogenation of the phenolic monoterpenes thymol and carvacrol with chloroperoxidase. Green Chem. 16, 1104–1108 (2014).

38. Karich A, Scheibner K, Ullrich R. & Hofrichter M. Exploring the catalase activity of unspecific peroxygenases and the mechanism of peroxide-dependent heme destruction. J. Mol. Catal. B: Enzym. 134, 238–246 (2016).

39. Wu F, et al. Quercetagetin alleviates zearalenone-induced liver injury in rabbits through Keap1/Nrf2/ARE signaling pathway. Front Pharmacol. 14, 1271384 (2023).

40. Wu F, Wang H, Li S, Wei Z, Han S, Chen B. Effects of dietary supplementation with quercetagetin on nutrient digestibility, intestinal morphology, immunity, and antioxidant capacity of broilers. Front. Vet. Sci. 9, 1060140 (2022).

41 . Zhou, Y., et al. Characterization of enzymes specifically producing chiral flavor compounds (*R*)-and (*S*)-1-phenylethanol from tea (*Camellia sinensis*) flowers. Food Chem. 280, 27–33 (2019).

42. Bertolini, V., et al. Synthesis of α-hydroxy fatty acids from fatty acids by intermediate α-chlorination with TCCA under solvent-free conditions: A way to valorization of waste fat biomasses. ACS Omega 6, 31901–31906 (2021).

43. Tonin, F., et al. Pilot-scale production of peroxygenase from *Agrocybe aegerita*. Org. Process Res. Dev. 25, 1414–1418 (2021).

44. Giuriato, D., et al. Design of a H_2_O_2_-generating P450_SPα_ fusion protein for high yield fatty acid conversion. Protein Sci. 31, e4501 (2022).

45. Qu, G., et al. The crucial role of methodology development in directed evolution of selective enzymes. Angew. Chem. Int. Ed. 59, 13204–13231 (2020).

46. Jiang, Y. Li, Z. Wang, C. Zhou, Y. J. Xu, H. & Li, S. Biochemical characterization of three new alpha-olefin-producing P450 fatty acid decarboxylases with a halophilic property. Biotechnol Biofuels 12, 79 (2019).

47. Ramirez-Escudero, M. Molina-Espeja, P. Gomez de Santos, P. Hofrichter, M. Sanz-Aparicio, J. & Alcalde, M. Structural insights into the substrate promiscuity of a laboratory-evolved peroxygenase. ACS Chem. Biol. 13, 3259–3268 (2018).

48. Sundaramoorthy, M. Terner, J. & Poulosl, T. L. The crystal structure of chloroperoxidase: a heme peroxidase-cytochrome P450 functional hybrid. Structures 3, 1367–1377 (1995).

49. Belcher, J., et al. Structure and biochemical properties of the alkene producing cytochrome P450 OleT_JE_ (CYP152L1) from the *Jeotgalicoccus* sp. 8456 Bacterium. J. Biol. Chem. 289, 6535–6550 (2014).

50. Shoji, O., et al. Understanding substrate misrecognition of hydrogen peroxide dependent cytochrome P450 from *Bacillus subtilis*. J. Biol. Inorg. Chem. 15, 1331–1339 (2010).

51. Fujishiro, T., et al. Chiral-substrate-assisted stereoselective epoxidation catalyzed by H_2_O_2_-dependent cytochrome P450_SPα_. Chem-Asian J. 7, 2286–2293 (2012).

52. Søndergaard, C. R. Olsson Mats H. M. Rostkowski, M. & Jensen, J. H. Improved treatment of ligands and coupling effects in empirical calculation and rationalization of pKa values. J. Chem. Theory Comput. 7, 2284–2295 (2011).

53. Trott, O. & Olson, A. J. AutoDock Vina: Improving the speed and accuracy of docking with a new scoring function, efficient optimization, and multithreading. J. Comput. Chem. 31, 455–461 (2009).

54. Pettersen, E. F., et al. UCSF Chimera—A visualization system for exploratory research and analysis. J. Comput. Chem. 25, 1605–1612 (2004).

55. Bayly, C. I. Cieplak, P. Cornell, W. D. & Kollman, P. A. A well-behaved electrostatic potential based method using charge restraints for deriving atomic charges: The RESP Model. J. Phys. Chem. C. 97, 10269–10280 (1993).

56. Maier, J. A. Martinez, C. Kasavajhala, K. Wickstrom, L. Hauser, K. E. & Simmerling, C. ff14SB: Improving the accuracy of protein side chain and backbone parameters from ff99SB. J. Chem. Theory. Comput. 11, 3696–3713 (2015).

57. Li, P. & Merz, K. M. MCPB.py: A python based metal center parameter builder. J. Chem. Inf. Model. 56, 599–604 (2016).

58. Li, P. & Merz, K. M. Metal ion modeling using classical mechanics. Chem. Rev. 117, 1564–1686 (2017).

59. Izaguirre, J. A. Catarello, D. P. Wozniak, J. M. & Skeel, R. D. Langevin stabilization of molecular dynamics. J. Chem. Phys. 114, 2090–2098 (2001).

60. Berendsen, H. J. C. Postma, J. P. M. Gunsteren, W. F. DiNola, A. & Haak, J. R. Molecular dynamics with coupling to an external bath. J. Chem. Phys. 81, 3684–3690 (1984).

61. Krutler, V. Gunsteren, W. F. & Hnenberger, P. H. A fast SHAKE algorithm to solve distance constraint equations for small molecules in molecular dynamics simulations. J. Comput. Chem. 22, 501–508 (2001).

62. Darden, T. York, D. & Pedersen, L. Particle mesh Ewald: An N⋅log(N) method for Ewald sums in large systems. J. Chem. Phys. 98, 10089–10092 (1993).

63. Ylilauri, M. & Pentikäinen, O. T. MMGBSA As a tool to understand the binding affinities of filamin–peptide interactions. J. Chem. Inf. Model. 53, 2626–2633 (2013).

64. Case, D. A. et al. AMBER 2018 (University of California, 2018).

65. Lu, Y. et al. Open-source, python-based redevelopment of the ChemShell multiscale QM/MM environment. J. Chem. Theory. Comput. 15, 1317–1328 (2019).

66. Kastner, J. Carr, J. M. Keal, T. W. Thiel, W. & Wander, A. DL-FIND: An open-source geometry optimizer for atomistic simulations. J. Phys. Chem. A 113, 11856–11865 (2009).

67. Metz, S. Kästner, J. Sokol, A. A. Keal, T. W. & Sherwood, P. ChemShell—a modular software package for QM/MM simulations. Wires. Comput. Mol. Sci. 4, 101–110 (2014).

68. Neese, F. The ORCA program system. Wires. Comput. Mol. Sci. 2, 73–78 (2012).

69. Neese, F. Software update: the ORCA program system, version 4.0. Wires. Comput. Mol. Sci. 8, e1327 (2018).

70. Neese F. Wennmohs, F. Becker, U. & Riplinger, C. The ORCA quantum chemistry program package. J. Chem. Phys. 152, 224108 (2020).

71. Neese, F. Software update: The ORCA program system—Version 5.0. Wires. Comput. Mol. Sci. 12, e1606 (2022).

72. Helmich-Paris, B. Souza, B. Neese, F. & Izsák R. An improved chain of spheres for exchange algorithm. J. Chem. Phys. 155, 104109 (2021).

73. Smith, W. Yong, C. W. & Rodger, P. M. DL_POLY: Application to molecular simulation. Mol. Simul. 28, 385–471 (2002).

74. Bakowies, D. & Thiel, W. Hybrid models for combined quantum mechanical and molecular mechanical approaches. J. Chem. Phys. 100, 10580–10594 (1996).

75. Handy, N. C. & Cohen, A. J. Left-right correlation energy. Mol. Phys. 99, 403–412 (2009).

76. Lee, C. Yang, W. & Parr, R. G. Development of the Colle-Salvetti correlation-energy formula into a functional of the electron density. Phys. Rev. B 37, 785–789 (1988).

77. Billeter, S. R. Turner, A. J. & Thiel, W. Linear scaling geometry optimisation and transition state search in hybrid delocalised internal coordinates. Phys. Chem. Chem. Phys. 2, 2177–2186 (2000).

78. Frisch, M. J. et al. *Gaussian 16, Revision A.03* (Gaussian, Inc., 2016).

79. Becke, A. D. Density-functional thermochemistry. II. The effect of the Perdew–Wang generalized-gradient correlation correction. J. Chem. Phys. 97, 9173–9177 (1992).

80. Becke, A. D. Density-functional thermochemistry. III. The role of exact exchange. J. Chem. Phys. 98, 5648–5652 (1993).

81. Weigend, F. & Ahlrichs, R. Balanced basis sets of split valence, triple zeta valence and quadruple zeta valence quality for H to Rn: Design and assessment of accuracy. Phys. Chem. Chem. Phys. 7, 3297–3305 (2005).

82. Marenich, A. V. Cramer, C. J. & Truhlar, D. G. Universal Solvation Model Based on Solute Electron Density and on a Continuum Model of the Solvent Defined by the Bulk Dielectric Constant and Atomic Surface Tensions. J Phys Chem 113, 6378–6396 (2009).

83. Grimme, S. Ehrlich, S. & Goerigk, L. Effect of the damping function in dispersion corrected density functional theory. J. Comput. Chem. 32, 1456–1465 (2011).

84. Grimme, S. Semiempirical GGA-type density functional constructed with a longrange dispersion correction. J. Comput. Chem. 27, 1787–1799 (2006).

