## Supplemental Information for "Unravelling an Efficient O_2_/Reductant-Dependent Catalytic Route in Heme Peroxygenases for Sustainable Applications"

**This file includes**

**Supplementary Methods	3**

**1. General information	3**

**2. Enzyme, strains and plasmids	3**

**3. Typical procedure for NBD activity	3**

**4. Determination of enzyme concentration	3**

**5. Determination of hydrogen peroxide concentration	4**

**6. Molecular Mechanics Generalized Born Surface Area (MM-GBSA) Free Energy Calculations Process	4**

**7. Classical Molecular Dynamics Simulations Process	4**

**Supplementary Tables 1-6	6**

**Supplementary Figures 1-29	16**

**Supplementary References	47**

**Supplementary Methods**

**1. General information**

Unless mentioned otherwise, all chemicals were purchased from HUSHI (Shanghai, China), Macklin and Aladdin and were used without further purification. Columns and column material for enzyme purification were purchased from GE Healthcare.

**2. Enzyme, strains and plasmids**

PaDa-Ⅰ, a mutant of *Aae*UPO (from *Agrocybe aegerita*)^[1]^ and *Mro*UPO (from *Marasmius rotula*)^[2]^, were optimized with *Pichia pastoris* codon and synthesized by Sangon (Shanghai, China). P450_SPα_ (from *Sphingomonas paucimobilis*)^[3]^, P450_BSβ_ (from *Bacillus subtilis*)^[4]^ and OleT_JE_ (from *Jeotgalicoccus sp.* ATCC 8456)^[5]^ were optimized with *E. coli* codon and synthesized by Sangon (Shanghai, China). *Cfu*CPO (from *Caldariomyces fumago*)^[6]^ was purchased from Sigma-Aldrich.

*E. coli* DH5α was used as host for gene cloning. *E. coli* BL21 (DE3) was used as hosts for P450 peroxygenase (OleT_JE_, P450_SPα_ and P450_BSβ_) expression. *P. pastoris* X33 was used as host for unspecific peroxygenase (*Aae*UPO and *Mro*UPO) expression.

The plasmid pRSFDuet-1 (Novagen, Germany) were used as vector for P450 peroxygenase gene cloning and protein expression. The plasmid pPICZ-A (Invitrogen, American) were used as vector for unspecific peroxygenase gene cloning and protein expression.

**3. Typical procedure for NBD activity**

The method is a slight variation on previous reports ^[7]^. The reaction mixture contains in a total of 1 ml following ingredients: 790 μL~880 potassium phosphate buffer (100 mM, pH 7.0), 100 μL 5-nitro-1,3-benzodioxole (NBD; 5 mM in 100% acetonitrile) and 10~100 μL of the enzyme sample. The reaction is started by the addition of 10 μL H_2_O_2_ (100 mM) and followed at 425 nm and room temperature over 15~30 seconds. Peroxygenase activity is calculated using the initial linear increase in absorbance and the extinction coefficient for 4-nitrocatechol at 425 nm (ε_425_ = 9,700 M^-1^ cm^-1^).

**4. Determination of enzyme concentration**

Determination of heme-peroxygenase concentration was by the method of Omura and Sato using an extinction coefficient of 91 mM^-1^ cm^-1^ in the reduced/carbon monoxide-bound minus reduced absorption difference spectrum.

**5. Determination of hydrogen peroxide concentration**

The method is a slight variation on previous reports ^[8]^.10 μL of 25 mM Xylenol Orange and 20 μL of 25 mM Fe (NH_4_)_2_(SO_4_)_2_ containing 2.5 M H_2_SO_4_ were added to 180 μL of the sample. After 30 min incubation at room temperature, absorbance of the samples was measured at 560 nm.

**6. Molecular Mechanics Generalized Born Surface Area (MM-GBSA) Free Energy Calculations Process**

The 4 ns trajectories obtained from the ff14SB MD simulations were analyzed using MMPBSA calculations for DHA, DHAA and AscA in complex. The default dielectric constant value (ε = 1) and the salt concentration of 0.1 M were set. Frames which used for these calculations were collected every 10 ps during the 4 ns MD simulations of the complexes. In particular, AscA did not have a stable binding in *Aae*UPO during the whole 50-ns trajectory, resulting in the inability to calculate binding free energy in this complex system.

**7. Classical Molecular Dynamics Simulations Process**

The complex system first was subjected to 5000 steps of steepest descent and 5000 steps of conjugate gradient minimization, with the protein held fixed by using position restraints with a force constant of 500 kcal·mol^−1^Å^−2^. And an additional 5000 steps of steepest descent and 25000 steps of conjugate gradient minimization was performed to fully optimize the system without restraints. Then, the systems were annealed from 0 to 300 K for 50 ps with the NVT ensemble, during which the constraint of 25 kcal/mol/Å was applied. Then, the density equilibration for 1.0 ns was conducted under the NPT ensemble to obtain a uniform density, where the target temperature of 300 K was kept with the Langevin thermostat and a 2 ps collision frequency, and the 1.0 atm target pressure was maintained with the Berendsen barostat and a pressure relaxation time of 1 ps. Subsequently, all the restraints on the complex systems were removed, and the enzyme complexes were equilibrated for 4 ns under the NPT ensemble. Lastly, a productive MD simulation of 50 ns was carried out under the NPT ensemble. During the simulations, the covalent bonds involving hydrogen atoms were constrained with the SHAKE method, and the integration step was set to 2 fs. A cutoff radius of 8 Å was set for nonbonded interactions, while the long-range electrostatic interactions were treated using the Particle Mesh Ewald (PME) method. All MD simulations were performed using Amber 18 software package.

**Supplementary Tables 1-6**

**Supplementary Table 1.** Binding free energies of AscA-enzyme, DHA-enzyme and DHAA-enzyme complex systems, calculated by MM/GBSA.

| Energy (kcal·mol^–1^) | | *Aae*UPO | *Mro*UPO | OleT_JE_ | *Cfu*CPO | P450_BSβ_ | P450_SPα_ |
| --- | --- | --- | --- | --- | --- | --- | --- |
| ΔG_bind,AscA_ | / | | -8.13 | -13.35 | -0.66 | -14.00 | -2.14 |
| ΔG_bind,DHA_ | -14.39 | | -10.07 | -15.92 | -12.21 | -14.51 | -10.04 |
| ΔG_bind,DHAA_ | -27.89 | | -13.90 | -23.41 | -16.96 | -18.37 | -13.73 |

**Supplementary Table 2. Plasmid and expression host**

| **Plasmid** | **Description** | **Expression Host** |
| --- | --- | --- |
| 1 | pPICZ-A carring *Aae*UPO (PaDa-Ⅰ) | *P. pastoris* X33 |
| 2 | pPICZ-A carring *Aae*UPO, TEV sequence and C-6×HIS | *P. pastoris* X33 |
| 3 | pPICZ-A carring *Mro*UPO | *P. pastoris* X33 |
| 4 | pPICZ-A carring *Mro*UPO and C-6×HIS | *P. pastoris* X33 |
| 5 | pRSFDuet-1 carring P450_SPα_ | *E. coli* BL21 |
| 6 | pRSFDuet-1 carring P450_SPα_ and SUMO | *E. coli* BL21 |
| 7 | pRSFDuet-1 carring P450_BSβ_ | *E. coli* BL21 |
| 8 | pRSFDuet-1 carring P450 OleT_JE_ | *E. coli* BL21 |

TEV sequence means 5’-Gly- Gln- Phe- Tyr- Leu- Asn- Glu-3’.

C-6×HIS means adding 6 HIS to the C-terminal.

**Supplementary Table 3. Synthetic gene sequences**

***Aae*UPO (PaDa-Ⅰ) from *Agrocybe aegerita***

ATGAAGTACTTTCCACTGTTCCCCACCTTGGTTTACGCCGTTGGTGTTGTTGCTTTCCCAGACTACGCTTCTTTGGCTGGTTTGTCCCAACAAGAGTTGGACGCTATCATCCCAACTTTGGAAGCTAGAGAACCAGGTTTGCCACCAGGTCCTTTGGAAAACTCTTCCGCCAAGTTGGTTAACGACGAAGCTCATCCATGGAAGCCACTTAGACCAGGTGATATCAGAGGTCCATGTCCAGGTTTGAACACTTTGGCTTCTCACGGTTACTTGCCAAGAAACGGTGTTGCTACTCCAGCTCAGATCATCAACGCTGTTCAAGAGGGTTTCAACTTCGACAACCAGGCTGCTATCTTCGCTACTTACGCTGCTCATTTGGTCGACGGTAACTTGATCACTGACTTGCTGTCCATCGGTAGAAAGACCAGATTGACTGGTCCAGATCCACCACCACCAGCTTCAGTTGGTGGTTTGAACGAACACGGTACTTTCGAAGGTGACGCTTCCATGACTAGAGGTGATGCTTTCTTCGGTAACAACCACGACTTCAACGAGACTTTGTTCGAGCAGTTGGTCGACTACTCCAACAGATTCGGTGGTGGTAAGTACAACTTGACTGTTGCTGGTGAGCTGAGGTTCAAGAGAATCCAAGACTCCATTGCCACCAATCCAAACTTCAGCTTCGTGGACTTCAGATTCTTCACTGCCTACGGTGAGACTACTTTCCCAGCCAACTTGTTCGTTGACGGTAGAAGAGATGACGGTCAGTTGGATATGGACGCTGCCAGATCATTCTTCCAGTTCTCTAGAATGCCAGACGACTTCTTCAGAGCCCCATCTCCAAGATCTGGTACTGGTGTTGAGGTTGTTGTTCAGGCTCATCCAATGCAGCCAGGTAGAAACGTTGGTAAGATCAACTCCTATACTGTCGACCCAACTTCCTCCGACTTCTCCACTCCATGTTTGATGTACGAGAAGTTCGTCAACATCACCGTCAAGTCTCTGTACCCAAATCCAACCGTCCAGTTGAGAAAGGCCTTGAACACTAACCTGGACTTTCTGTTCCAAGGTGTTGCTGCTGGTTGTACCCAAGTTTTCCCATACGGTAGAGATTAA

***Mro*UPO from *Marasmius rotula***

ATGAAGTTGGCTATTTCTTCTTCTTTGATTGCTTTGGTTTCTGTTACTACTGCTTTGGCTAACTCTCAAGATGTTGTTGATTTCGGTTCTGCTCATCCATGGAAAGCTCCAGGTCCTAATGATTCTAGAGGTCCATGTCCAGGTTTGAACACTTTGGCTAACCATGGTTTCTTGCCTAGAAATGGTAGAAACATTTCTGTTCCAATGATTGTTAAGGCTGGTTTCGAAGGTTATAATGTTCAATCTGATATCTTGATCCTGGCTGGTAAGATTGGTATGTTGACTTCTAGAGAAGCTGATACTATTTCTTTGGAAGATTTGAAACTGCATGGTACTATTGAACATGATGCTTCTTTGTCTAGAGAAGATGTTGCTATTGGTGATAATTTGCATTTCAACGAAGCTATTTTCACTACTTTGGCTAATTCTAACCCAGGTGCTGATGTTTATAACATTTCTTCTGCTGCTCAAGTTCAACATGATAGATTGGCTGATTCTTTGGCTAGAAATCCTAATGTTACTAATACTGATCTGACTGCTACTATTAGATCTTCTGAATCTGCTTTTTTCTTGACTGTTATGTCTGCTGGTGATCCATTGAGAGGTGAAGCTCCAAAGAAGTTCGTTAACGTTTTTTTTAGAGAGGAAAGAATGCCTATTAAGGAAGGTTGGAAGAGATCTACTACTCCTATTACTATTCCATTGTTGGGTCCTATTATTGAAAGAATTACTGAATTGTCCGACTGGAAACCAACTGGTGATAACTGTGGTGCTATTGTTTTGTCTCCTGAATTGTAA

**P450_SPα_ from *Sphingomonas paucimobilis* with SUMO**

ATGTCGGACTCAGAAGTCAATCAAGAAGCTAAGCCAGAGGTCAAGCCAGAAGTCAAGCCTGAGACTCACATCAATTTAAAGGTGTCCGATGGATCTTCAGAGATCTTCTTCAAGATCAAAAAGACCACTCCTTTAAGAAGGCTGATGGAAGCGTTCGCTAAAAGACAGGGTAAGGAAATGGACTCCTTAAGATTCTTGTACGACGGTATTAGAATTCAAGCTGATCAGACCCCTGAAGATTTGGACATGGAGGATAACGATATTATTGAGGCTCACAGAGAACAGATTGGTGGTATGCCGAAAACCCCGCACACCAAAGGTCCGGATGAAACCCTGAGCCTGCTGGCTGATCCGTACCGCTTCATCTCTCGTCAGTGTCAGCGTCTGGGCGCGAACGCGTTCGAATCTCGCTTCCTGCTGAAAAAGACCAACTGCCTGAAAGGCGCGAAAGCGGCGGAAATCTTCTACGACACCACCCGTTTCGAACGTGAAGGCGCTATGCCGGTTGCTATCCAGAAAACCCTGCTGGGCCAGGGCGGTGTTCAGGGCCTGGATGGTGAAACTCACCGTCACCGTAAACAGATGTTCATGGGCCTGATGACCCCGGAACGTGTGCGCGCGCTGGCGCAGCTGTTCGAAGCGGAATGGCGTCGTGCGGTGCCGGGCTGGACCCGCAAAGGCGAAATCGTATTTTATGACGAACTGCACGAACCGCTGACTCGCGCGGTGTGCGCGTGGGCGGGTGTGCCGCTGCCGGACGACGAAGCGGGCAACCGCGCGGGTGAACTGCGCGCGCTGTTCGATGCCGCTGGTAGCGCGTCCCCGCGTCACCTGTGGAGCCGTCTGGCCCGTCGTCGCGTTGACGCGTGGGCGAAACGTATCATCGAAGGTATCCGTGCTGGTTCCATCGGTAGCGGTAGCGGCACCGCGGCGTACGCTATCGCGTGGCACCGCGATCGTCACGATGATCTGCTGAGCCCGCACGTTGCGGCGGTTGAACTGGTTAACGTTCTGCGTCCGACCGTGGCTATCGCGGTGTACATCACCTTCGTGGCACATGCACTGCAGACCTGCTCCGGCATTCGCGCAGCTCTGGTTCAGCAGCCGGATTATGCAGAACTGTTCGTGCAGGAAGTTCGTCGTTTCTACCCGTTCTTCCCGGCGGTGGTTGCGCGCGCAAGCCAGGATTTCGAATGGGAAGGCATGGCGTTCCCGGAAGGCCGTCAGGTTGTTCTGGACCTGTATGGCTCTAACCATGACGCGGCGACCTGGGCGGACCCGCAGGAATTTCGTCCGGAACGTTTCCGTGCGTGGGATGAAGATTCTTTCAACTTCATCCCGCAGGGCGGTGGTGATCACTACCTGGGCCACCGCTGCCCAGGTGAATGGATCGTTCTGGCTATTATGAAAGTTGCGGCTCACCTGCTGGTTAACGCTATGCGTTATGATGTTCCGGACCAGGATCTGAGCATTGATTTCGCACGTCTGCCGGCGCTGCCGAAATCCGGCTTCGTTATGCGTAACGTTCACATCGGCGGTTAA

**P450_BSβ_ from *Bacillus subtilis***

ATGAACGAACAAATCCCACACGATAAATCCCTGGACAACTCTCTGACTCTGCTGAAAGAAGGTTATCTGTTCATCAAAAACCGCACCGAGCGCTATAACTCCGATCTGTTCCAAGCTCGTCTGCTGGGCAAAAACTTCATCTGCATGACCGGTGCAGAAGCAGCGAAAGTCTTCTACGACACGGATCGTTTCCAGCGTCAGAACGCGCTGCCGAAACGTGTTCAAAAATCTCTGTTCGGCGTTAACGCGATCCAGGGTATGGATGGTTCTGCACACATTCACCGCAAAATGCTGTTCCTGTCTCTGATGACCCCACCACACCAGAAACGTCTGGCAGAACTGATGACTGAAGAATGGAAAGCCGCAGTTACCCGCTGGGAAAAAGCGGACGAGGTCGTTCTGTTTGAAGAAGCAAAAGAAATCCTGTGCCGTGTGGCTTGTTATTGGGCTGGTGTACCACTGAAAGAAACGGAAGTGAAAGAACGTGCGGATGACTTCATCGACATGGTCGATGCATTCGGCGCAGTAGGTCCGCGTCACTGGAAAGGTCGTCGTGCACGTCCACGTGCCGAGGAATGGATTGAAGTGATGATCGAAGACGCGCGTGCAGGCCTGCTGAAGACTACCTCTGGTACCGCGCTGCATGAAATGGCGTTCCACACCCAGGAGGACGGTAGCCAGCTGGACTCTCGCATGGCGGCGATCGAACTGATCAACGTTCTGCGTCCTATCGTGGCAATCTCCTACTTCCTGGTATTCTCCGCCCTGGCGCTGCACGAGCATCCGAAATACAAAGAATGGCTGCGTTCCGGTAATAGCCGCGAACGTGAAATGTTCGTGCAGGAAGTGCGTCGTTACTATCCGTTTGGTCCGTTCCTGGGTGCTCTGGTTAAAAAAGACTTCGTTTGGAACAACTGTGAATTCAAAAAAGGCACCTCTGTACTGCTGGATCTGTATGGTACCAACCACGACCCACGTCTGTGGGACCACCCTGATGAATTCCGTCCGGAGCGTTTTGCGGAGCGTGAAGAAAACCTGTTCGATATGATTCCGCAAGGTGGTGGCCACGCAGAAAAAGGCCATCGTTGTCCGGGCGAAGGTATTACTATCGAAGTGATGAAGGCGTCCCTGGACTTCCTGGTTCACCAAATCGAATACGATGTTCCGGAACAGAGCCTGCACTATTCTCTGGCTCGCATGCCGAGCCTGCCGGAATCTGGTTTTGTTATGAGCGGCATCCGTCGCAAAAGCTAA

**OleT_JE_ from *Jeotgalicoccus sp.* ATCC 8456**

ATGGCGACCCTGAAACGTGATAAAGGCCTGGATAACACCCTGAAAGTTCTGAAACAGGGTTACCTGTATACCACCAACCAGCGTAACCGCCTGAACACCAGCGTTTTCCAGACCAAAGCTCTGGGCGGTAAACCGTTCGTTGTGGTGACCGGTAAAGAAGGTGCGGAAATGTTCTACAACAACGATGTTGTTCAGCGTGAAGGTATGCTGCCGAAACGTATCGTTAACACCCTGTTCGGTAAAGGTGCAATCCACACCGTTGATGGTAAAAAACACGTGGACCGTAAAGCACTGTTCATGTCTCTGATGACCGAAGGTAACCTGAACTATGTTCGTGAACTGACCCGTACCCTGTGGCATGCGAACACCCAGCGTATGGAAAGCATGGATGAAGTTAACATCTATCGTGAATCTATCGTGCTGCTGACTAAAGTGGGCACCCGTTGGGCTGGTGTGCAGGCGCCGCCGGAAGATATCGAACGTATTGCAACCGACATGGATATTATGATTGATAGCTTCCGTGCACTGGGTGGTGCGTTCAAAGGTTACAAAGCATCTAAAGAAGCTCGTCGTCGCGTTGAAGATTGGCTGGAAGAACAGATCATTGAAACCCGTAAAGGTAACATCCACCCGCCGGAAGGCACCGCACTGTATGAATTTGCGCACTGGGAAGATTATCTGGGTAACCCGATGGACTCCCGTACCTGCGCAATTGACCTGATGAACACCTTTCGTCCGCTGATCGCAATCAACCGTTTCGTTAGCTTCGGTCTGCACGCGATGAACGAAAACCCGATTACCCGCGAAAAAATCAAATCCGAACCGGATTACGCTTACAAATTCGCGCAGGAAGTTCGTCGTTACTATCCGTTCGTTCCGTTTCTGCCGGGCAAAGCTAAAGTTGATATCGATTTCCAGGGCGTGACCATTCCGGCTGGCGTTGGTCTGGCGCTGGATGTTTATGGCACCACCCACGATGAATCTCTGTGGGATGATCCGAACGAATTTCGTCCGGAACGCTTCGAAACCTGGGATGGTAGCCCGTTCGACCTGATCCCGCAGGGCGGTGGTGATTACTGGACCAACCACCGTTGCGCGGGTGAATGGATCACCGTTATTATTATGGAAGAAACCATGAAATATTTCGCGGAAAAAATCACCTACGATGTGCCGGAACAGGATCTGGAAGTTGATCTGAACTCTATCCCAGGCTACGTTAAAAGCGGTTTCGTTATCAAAAACGTGCGTGAAGTTGTTGATCGTACCTAA

**Supplementary Table 4.** GC and HPLC analytics

| Column | Temperature program/ gradient | Retention time |
| --- | --- | --- |
| GC  CP-Chirasil-DEX CB (Agilent)  (25 m × 0.25 mm, 0.25 µm)  carrier gas: N_2_ | 100 ºC hold 5 min  40 ºC/min to 140 ºC hold 5 min  80 ºC/min to 180 ºC hold 1 min | 3.10 min ethylbenzene  9.18 min (*R*)-1-phenylethanol  9.41 min (*S*)-1-phenylethanol  6.95 min acetophenone  7.45 min dodecane |
| GC  SH-Rtx-WAX  (30 m × 0.25 mm, 0.25 µm)  carrier gas: N_2_ | 50 ºC hold 0 min  5 ºC/min to 110 ºC hold 0 min  80 ºC/min to 240 ºC hold 2 min | 10.14 min cyclohexanol  7.54 min cyclohexanone  5.96 min dodecane |
| GC  SH-Rtx-5  (30 m × 0.25 mm, 0.25 µm)  carrier gas: N_2_ | 80 ºC hold 5 min  10 ºC/min to 280 ºC hold 1 min | 16.74 min lauric acid  18.62 min α-OH of lauric acid  18.79 min β-OH of lauric acid  8.04 min 1-undecene  10.20 min dodecane |
| HPLC  Shimadzu C18  (4.6 × 250 mm, 5 μm)  flow rate: 1.0 mL/min | methanol: “water” (containing 0.2 % glacial acetic acid) = 2:98 | 4.51 min AscA  (245 nm) |
| HPLC  Shimadzu C18  (4.6 × 250 mm, 5 μm)  flow rate: 1.0 mL/min | acetonitrile: water = 70:30 | 6.82 min phenyl methyl sulfide  3.16 min phenyl methyl sulfoxide  (254 nm) |
| HPLC  Shimadzu C18  (4.6 × 250 mm, 5 μm)  flow rate: 1.0 mL/min | methanol: “water” (containing 0.2 % glacial acetic acid) = 44:56 | 3.45 min gallic acid (272 nm)  3.75 min pyrogallic acid (210 nm) |
| HPLC  Shimadzu C18  (4.6 × 250 mm, 5 μm)  flow rate: 1.0 mL/min | acetonitrile: water (containing 0.1 % formic acid) = 10:90;  0~1 min: 10% acetonitrile, 90% water; 1~15 min: acetonitrile increased from 10% to 90%, water reduced from 90% to 10%; 15~18 min: 10% acetonitrile, 90% water; | 9.71 min quercetagetin  11.25 min quercetin |

**Supplementary Table 5.** Spin density population of key atoms for key species involved in the Cpd I formation from the UPO/DHAA/O_2_ system.

| Species | Spin density | | | | | |
| --- | --- | --- | --- | --- | --- | --- |
|  | Fe | Op | Od | Cys-36 | HEM | DHAA |
| ^8^RC | 4.03 | 0.98 | 0.99 | 0.45 | 0.53 | 0.00 |
| ^2^RC | 2.47 | -0.90 | -0.93 | 0.41 | -0.05 | 0.00 |
| ^2^IM1 | 1.37 | 0.36 | 0.19 | 0.18 | -0.11 | -0.99 |
| ^2^IM2 | 0.87 | 0.14 | 0.01 | 0.08 | -0.11 | 0.00 |
| ^2^PC | 1.16 | 0.71 | 0.01 | -0.44 | -0.42 | 0.01 |

**Supplementary Table 6.** Spin density population of key atoms for key species involved in the oxygen activation and hydrogen peroxide formation from the UPO/GA/O_2_ system.

| Species | Spin density | | | | | |
| --- | --- | --- | --- | --- | --- | --- |
|  | Fe | Op | Od | Cys-36 | HEM | GA |
| ^8^RC | 3.95 | 0.98 | 1.01 | 0.42 | 0.63 | 0.00 |
| ^2^RC | 2.60 | -0.98 | -1.01 | 0.42 | -0.03 | 0.00 |
| ^2^IM1 | -0.13 | -0.20 | -0.01 | 0.36 | 0.00 | 0.97 |
| ^2^IM2 | 0.15 | 0.07 | 0.04 | -0.14 | -0.08 | 0.97 |
| ^2^PC | 1.34 | 0.00 | 0.00 | 0.16 | -0.04 | -0.45 |

**Supplementary Figures 1-29**

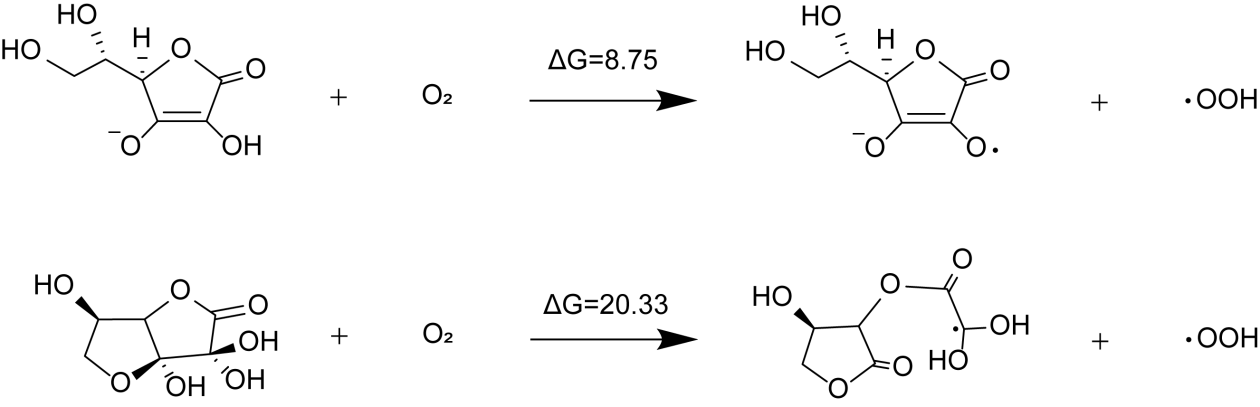

**Supplementary Figure 1.** Calculated reaction free energy (in kcal/mol) for the reduction of O_2_ to •OOH by DHAA and AscA in aqueous solution at the level of B3LYP/def2-TZVP.

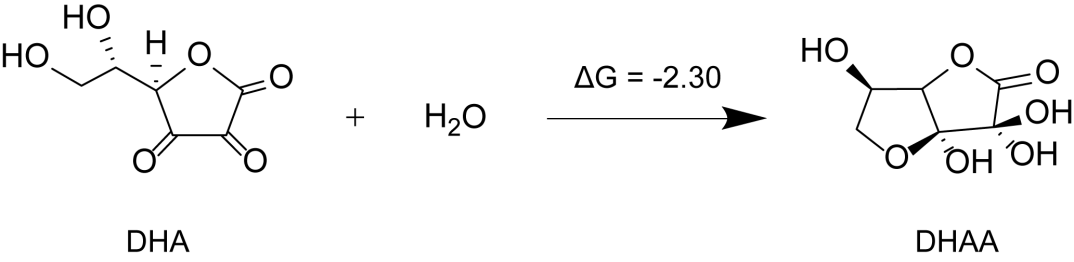

**Supplementary Figure 2.** Calculated reaction free energy (in kcal/mol) for the hydration of DHA to generate DHAA in aqueous solution at the level of B3LYP/def2-TZVP.

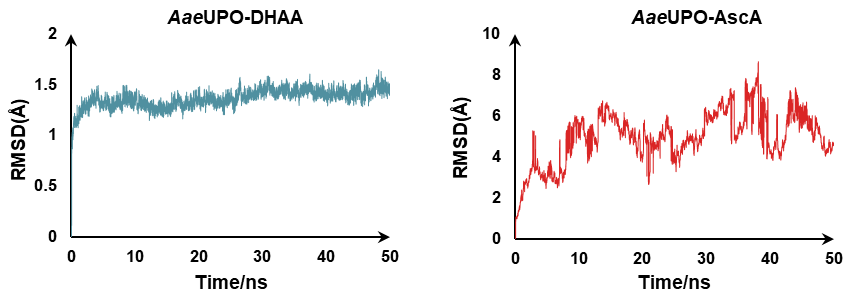

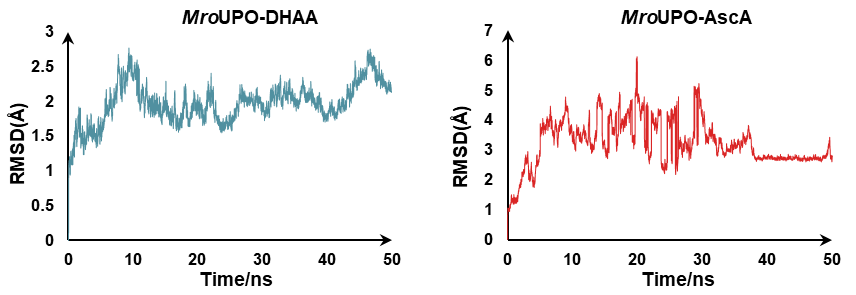

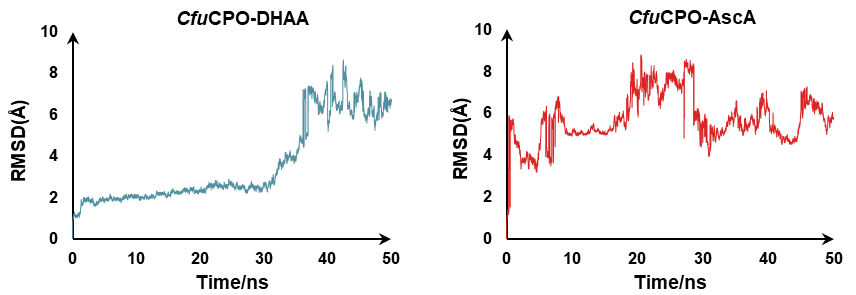

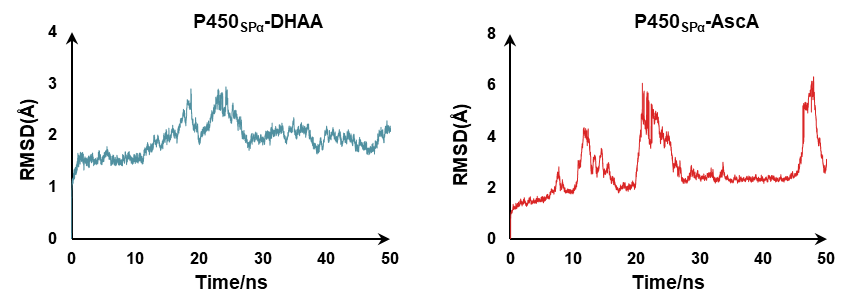

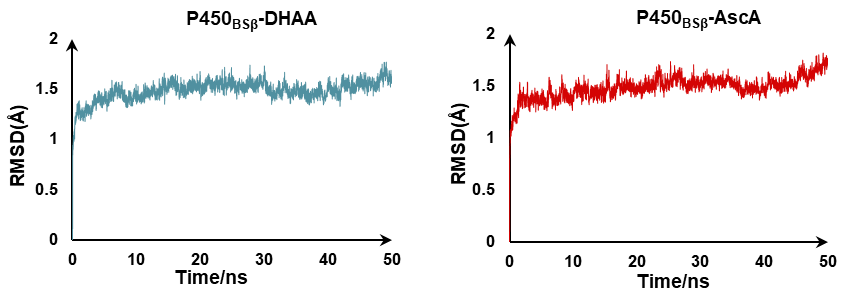

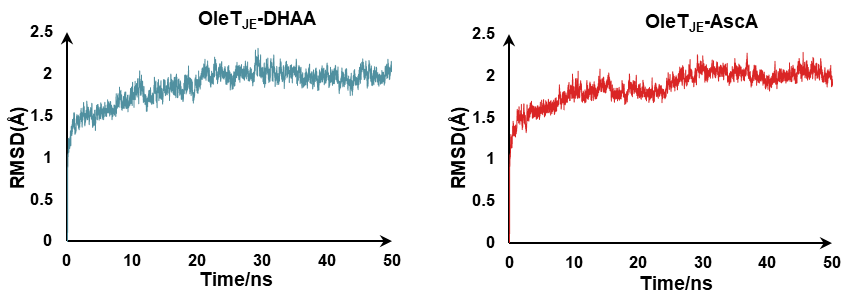

**Supplementary Figure 3.** Time evolution of the root mean square deviations (RMSD) for the proteins and ligands during the MD simulations.

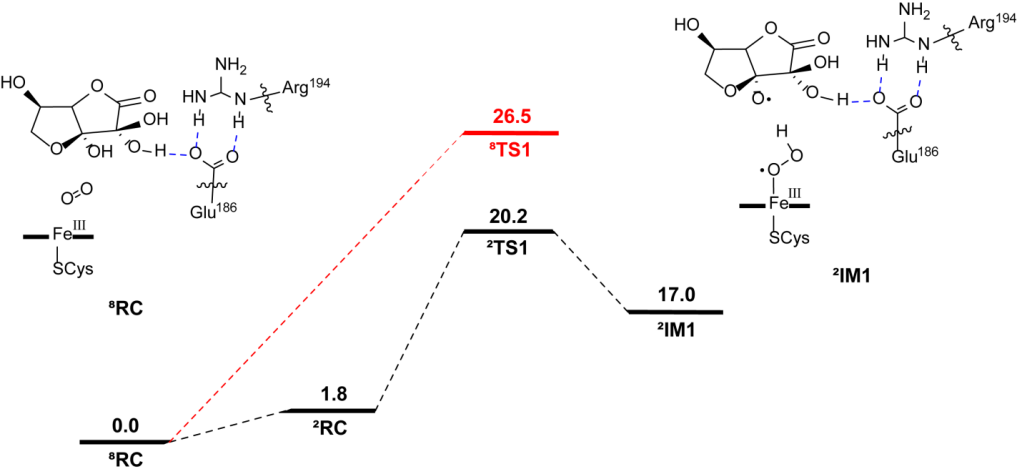

**Supplementary Figure 4.** Comparison of the QM(OLYP/def2-TZVP)/MM calculated energy profiles (in kcal/mol) for HAT reaction from ^8^RC and ^2^RC in the UPO/DHAA/O_2_ system.

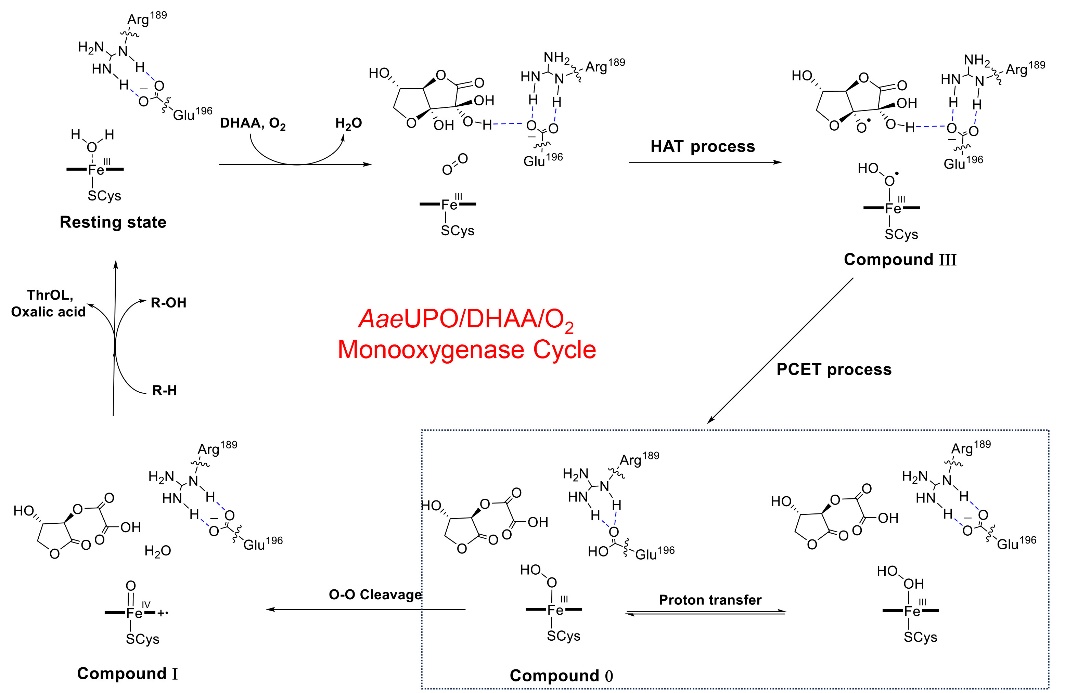

**Supplementary Figure 5.** Proposed catalytic cycle of UPO-catalyzed oxidative oxidations in the presence of O_2_/reductant (DHAA). In this route, no enzyme inactivation with biliverdin formation (heme bleaching) was observed. The catalytic cycle is initiated by the DHAA-mediated O_2_ binding to Fe. This is followed by H-atom transfer from DHAA to the O_2_ moiety to form the compound II. Then, proton transfer from the O atom of DHAA to Glu196 is coupled with the electron transfer from DHA to Fe(III)-•OOH radical species (PCET process), leading to the formation of Cpd 0. Finally, the proton transfers from the protonated Glu196 to the proximal O to generate Fe(III)-H_2_O_2_, or to the distal O to trigger the O-O cleavage to form the active species Cpd I. The formation of Fe(III)-H_2_O_2_ vs. Cpd 0 is reversible, but the further formation of Cpd is highly favorable thermodynamically.

**Supplementary Figure 6.** Mechanistic study on departure of byproduct of DHAA and entry of the substrate **1a**. **A**. QM(OLYP/def2-TZVP)/MM-calculated potential energy profile (in kcal/mol) for hydrolysis reaction. **B**. Proposed mechanisms for hydrolysis of DHAA. **C**. Comparison of the fluctuation of the SUB-Fe distance and the fluctuation of the ThrOL-Fe distance. **D**. The initial and final structures of the molecular dynamics simulation.

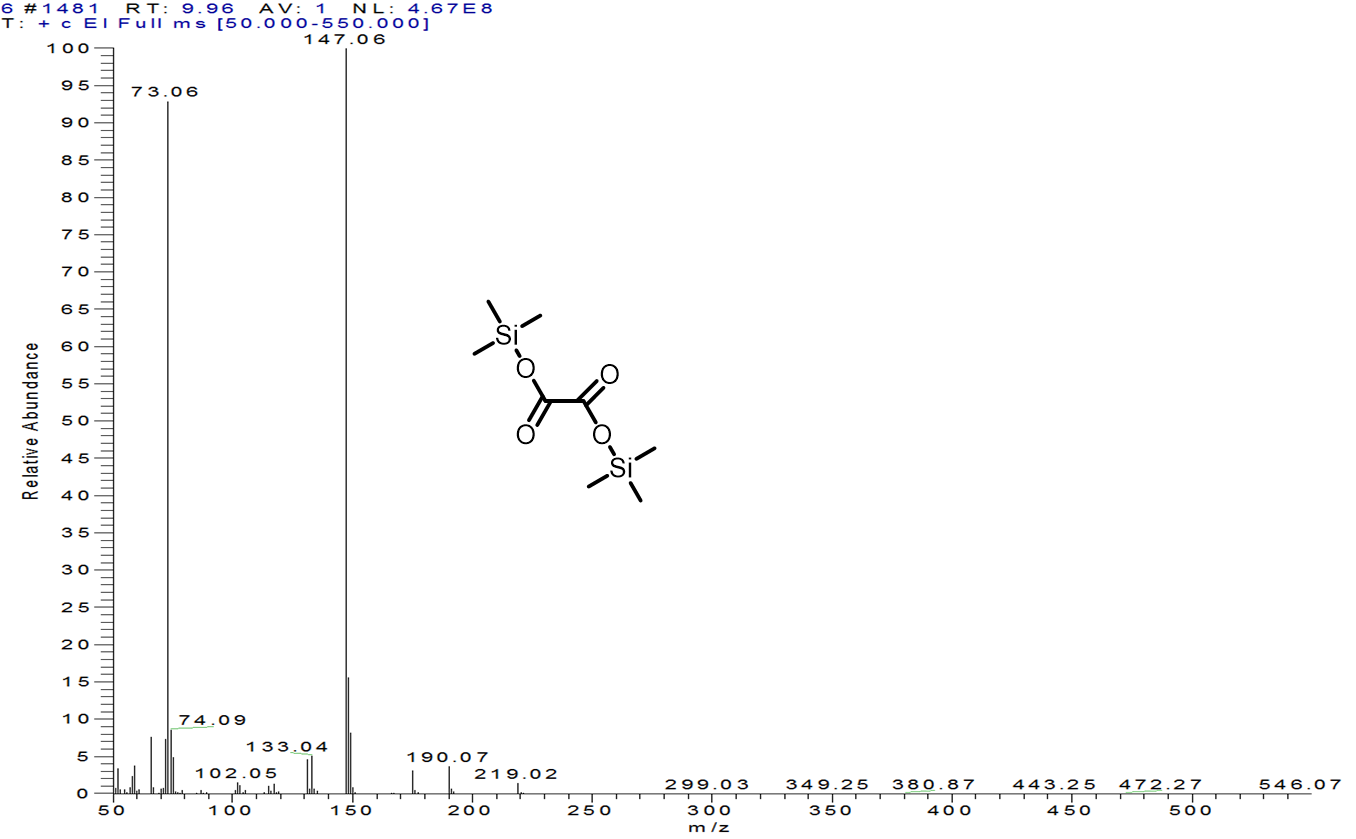

**Supplementary Figure 7.** GC-MS analysis of reaction mixture after the reaction of *Aae*UPO catalyzed hydroxylation of ethyl benzene with addition of AscA. Large amount of oxalic acid was detected in the reaction mixture and the fragmentation pattern was obtained for oxalic acid after derivatization. The GC–MS analysis was performed using the Thermo Scientific GC-MS equipped with a DB-5MS column (30 m×0.25 mm, 0.25 μm).

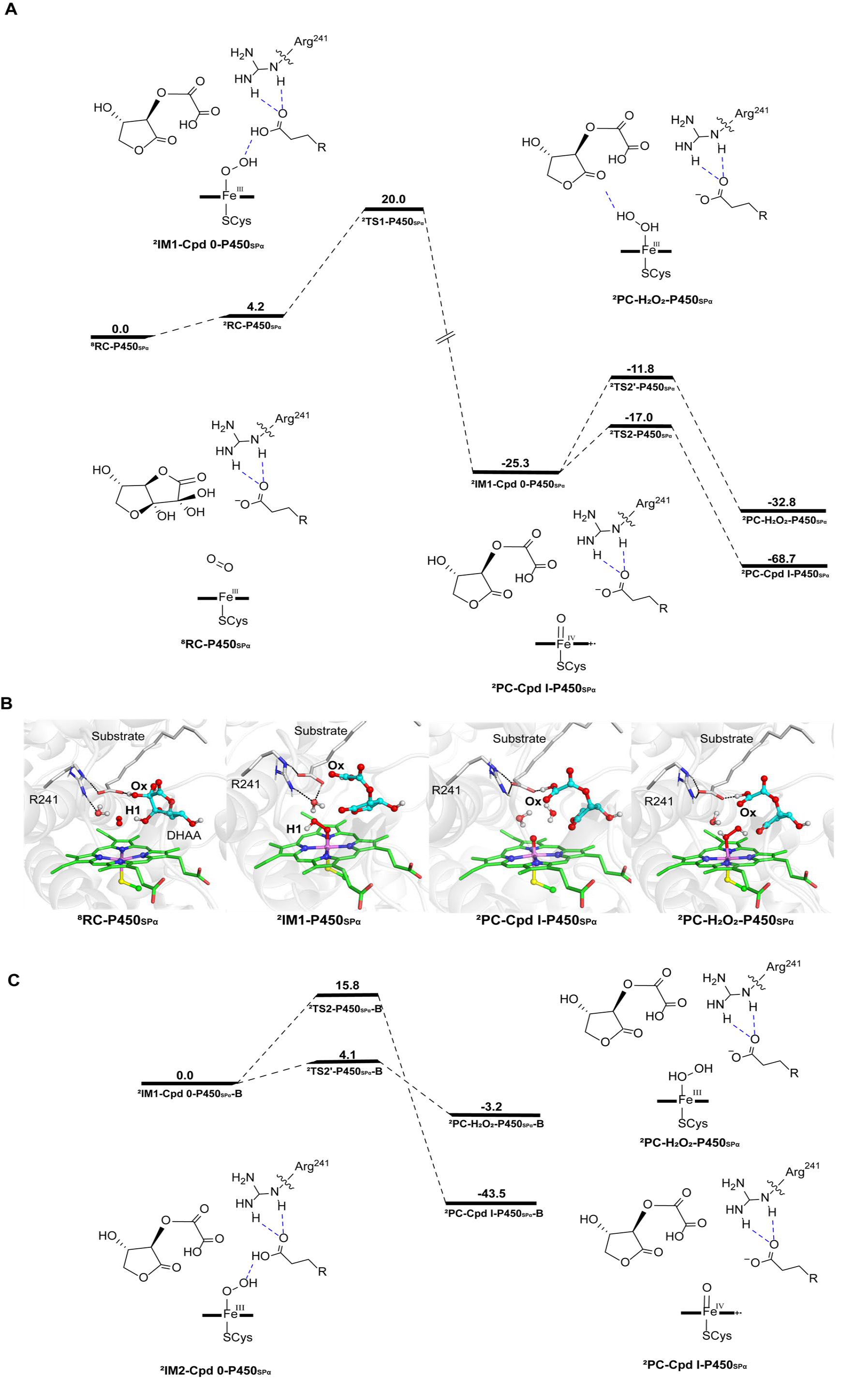

**Supplementary Figure 8.** QM/MM calculations for DHAA-mediated oxygen activation mechanism by P450_SPα_. **A**. QM(OLYP/def2-TZVP)/MM-calculated potential energy profile (in kcal/mol) for Cpd I and Fe(III)-H_2_O_2_ formation from the P450_SPα_/DHAA/O_2_ system. **B**. QM(OLYP/def2-SVP)/MM-optimized geometries of key species from the P450_SPα_/DHAA/O_2_ system. **C**. QM(B3LYP-D3/def2-TZVP)/MM-calculated potential energy profile (in kcal/mol) for Cpd I and Fe(III)-H_2_O_2_ formation from the Cpd 0 species in the P450_SPα_/DHAA/O_2_ system. It is seen that the DHAA binding and the active site architecture are quite similar to those in the UPO/DHAA/O_2_ system. The reaction begins with the O_2_ binding to Fe(III) center. This is followed by H-atom transfer from H1 to O_2_ moiety, which triggers the spontaneous proton-coupled electron transfer (PCET) reaction, resulting in the formation of Cpd 0. In the PCET process, the proton transfer from Ox of DHAA to the substrate carboxyl group is coupled to electron transfer from DHAA to Fe(III)-·OOH radical species. Cpd 0 can further convent the more stable species of Cpd I via the O-O heterolysis.

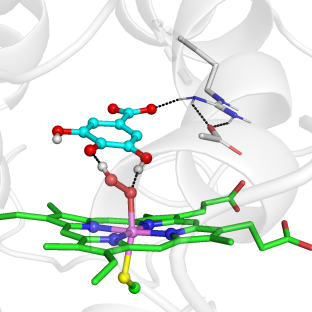

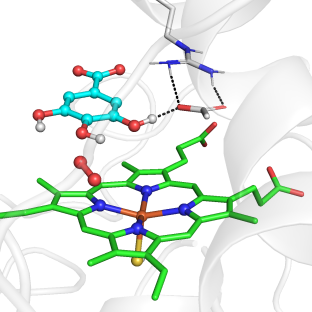

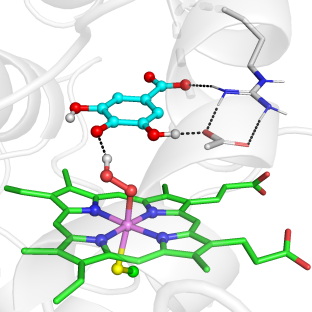

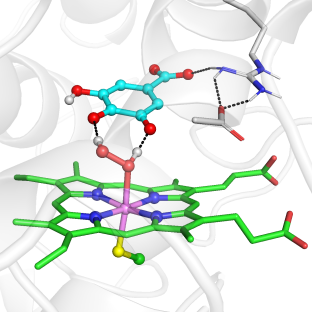

**E196**

**R189**

**^8^RC**

**R189**

**E196**

**R189**

**E196**

**R189**

**^2^IM1**

**^2^IM2**

**^2^PC**

1.37

1.39

1.86

1.43

**E196**

**Supplementary Figure 9.** QM (OLYP/def2-SVP)/MM-optimized geometries of key species involved in the oxygen activation and hydrogen peroxide formation from the UPO/GA/O_2_ system.

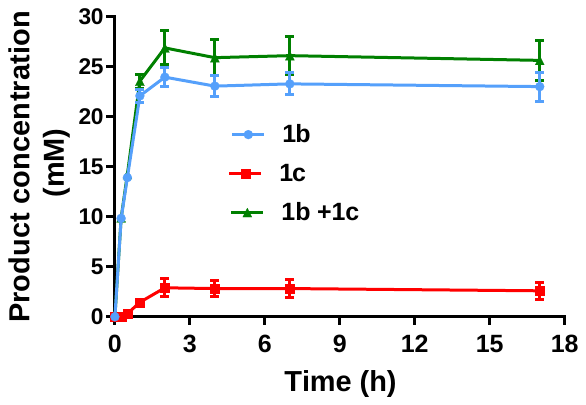

**Supplementary Figure 10.** Time course of r*Aae*UPO catalzyed hydroxylation of 1a with GA under aerobic conditions. Reactions were performed in potassium phosphate buffer (100 mM, pH 8.0, 30°C) containing r*Aae*UPO (100 nM), GA (100 mM), **1a** (50 mM), 5% (m/v) 2-hydroxypropyl-β-cyclodextrin.

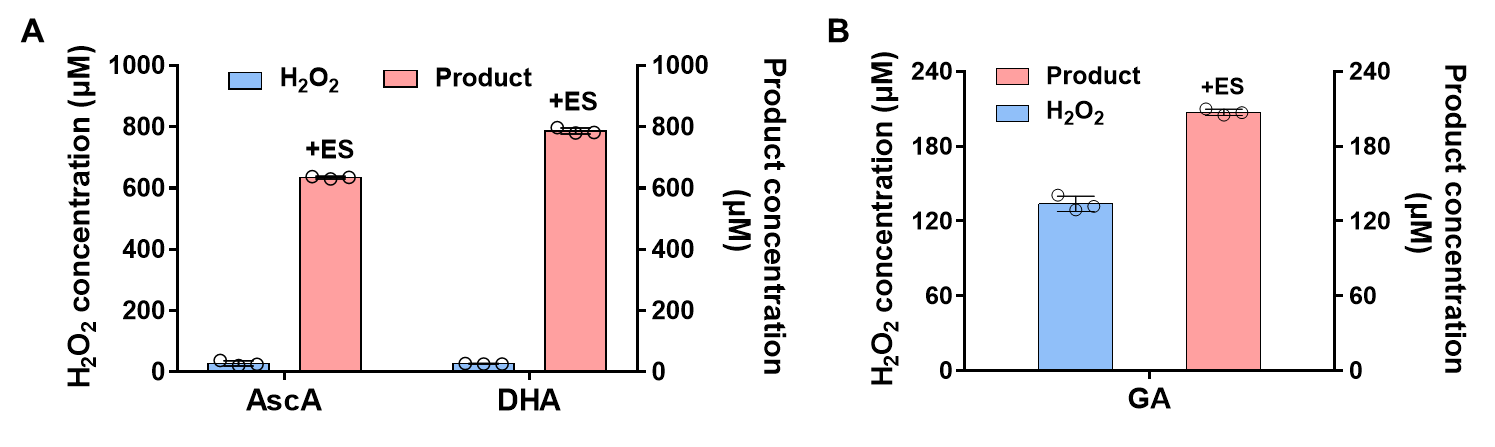

**Supplementary Figure 11.** Comparison of hydrogen peroxide production and product production in the presence or absence of r*Aae*UPO and ethylbenzene. Hydrogen peroxide produced by 1 mM GA incubation at 30 °C in buffer solution (0.1 M, pH 7.0) for 1h and products produced in the presence of r*Aae*UPO and ethylbenzene.

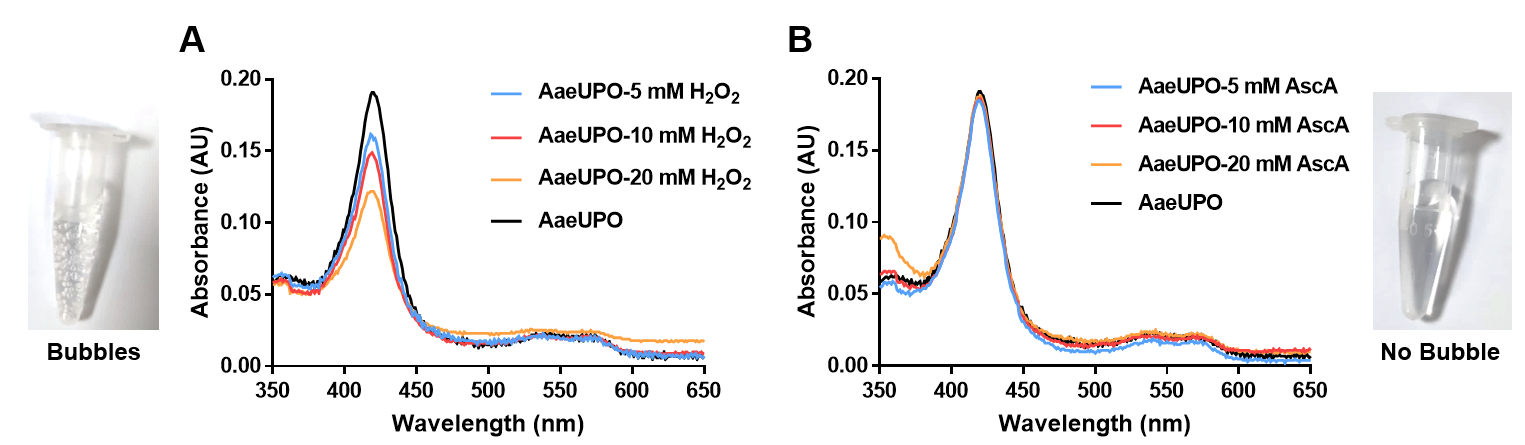

**Supplementary Figure 12.** UV–vis spectra analysis of r*Aae*UPO in phosphate buffer at pH 7. The enzymes were treated with different concentrations (Blue: 5 mM, Red: 10 mM, Orange: 20 mM.) of H_2_O_2_ (**A**) or AscA (**B**). The treatment with H_2_O_2_ led to the formation of O_2_ bubbles and pronounced heme-bleaching, while AscA treatment did not cause the O_2_ formation without heme-bleaching.

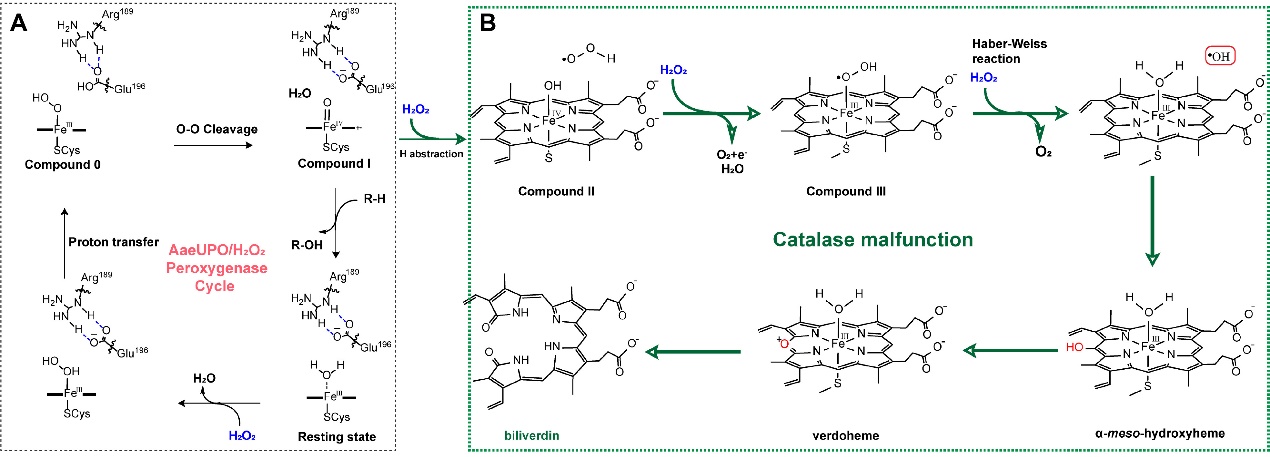

**Supplementary Figure 13.** The catalytic cycle of UPO-catalyzed oxidative reactions with exogenous H_2_O_2_ (**A**), along with the proposed catalase malfunction of UPO that leads to the heme destruction with heme bleaching and verdoheme and biliverdin formation (**B**). The catalytic cycle consists of the following steps: The reaction between a single molecule of H_2_O_2_ and r*Aae*UPO generates the active species of Compound I. Then, subsequent reactions between Compound I and an additional H_2_O_2_ lead to the sequential formation of Compound II and Compound III. Finally, Compound III reacts with the third H_2_O_2_ molecule to produce the hydroxyl radical, which would result in heme bleaching (cleavage of the protoporphyrin ring into biliverdin with Fe release), leading to heme destruction and enzyme inactivation.

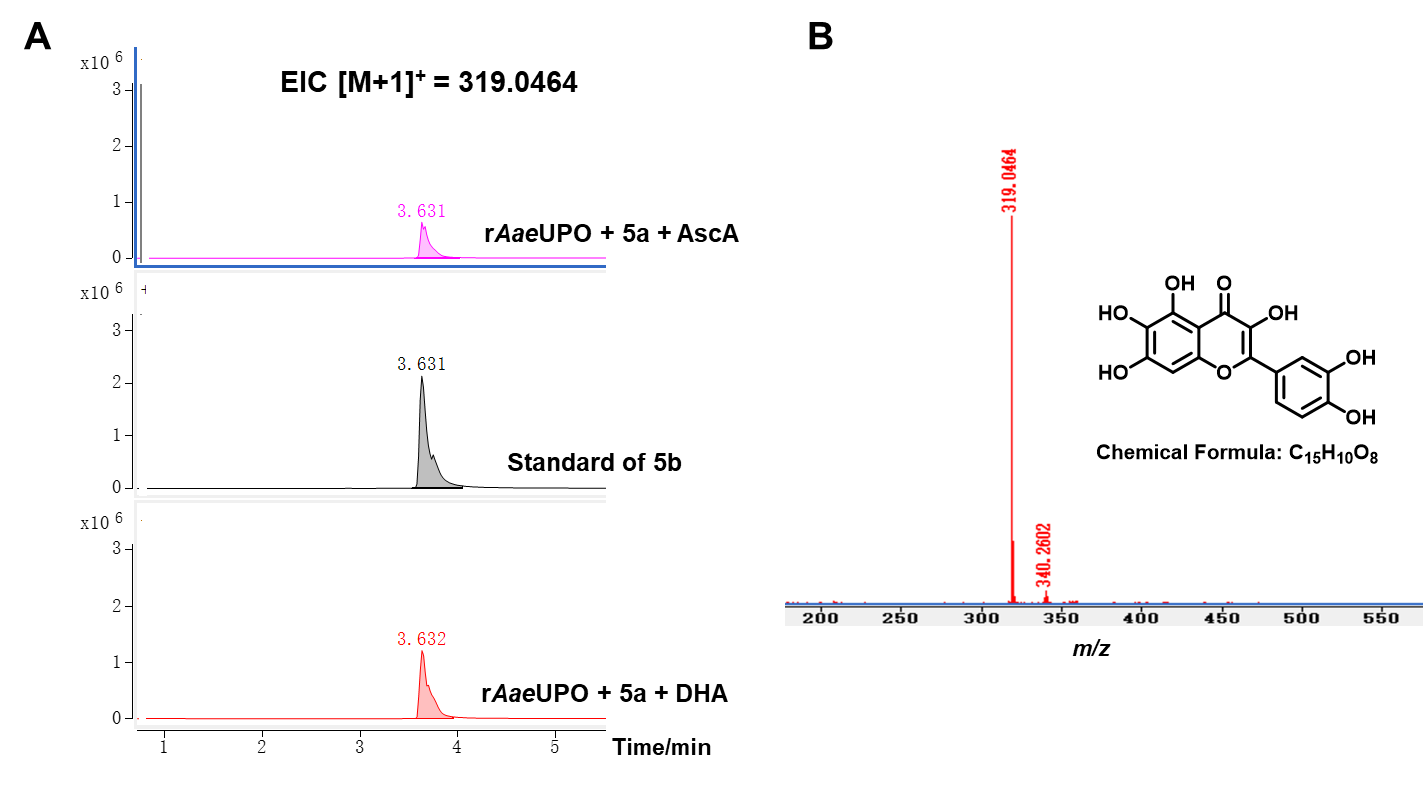

**Supplementary Figure 14.** LC-MS analysis of 5b. **A.** LC chromatograms of r*Aae*UPO-catalyzed hydroxylation of 5a in the presence of AscA or DHA. **B**. MS analysis of scutellarin in the r*Aae*UPO-catalyzed reactions with AscA or DHA. Reactions were performed in potassium phosphate buffer (100 mM, pH 8.0, 37°C) containing r*Aae*UPO (1 μM), AscA or DHA (50 mM), 5a (1.5 g/L), 5% (m/v) 2-hydroxypropyl-β-cyclodextrin, 10% acetone.

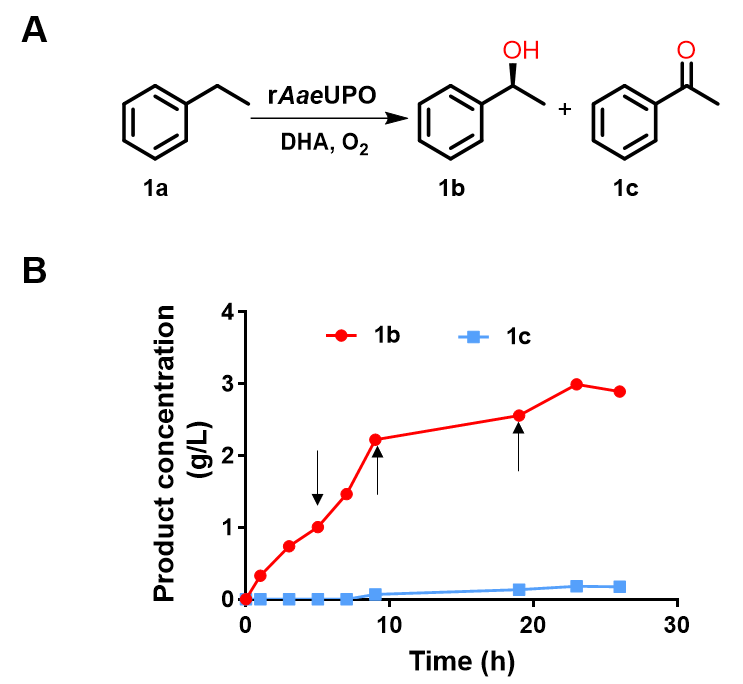

**Supplementary Figure 15.** The r*Aae*UPO-catalzyed hydroxylation of **1a** with DHA as reductants at 100 mL scale. **A**. Reaction scheme of r*Aae*UPO-catalyzed benzylic hydroxylation of **1a** using DHA as reductant. B. Scale-up reaction in 100 mL reaction mixture for the production of **1b** and **1c** by r*Aae*UPO-catalyzed hydroxylation of **1a** using DHA as the co-substrate. Reaction was performed in 100 mL potassium phosphate buffer (100 mM, pH 8.0, 30 °C) containing r*Aae*UPO (0.25 μM), 1a (200 mM), DHA (The reaction started with 50 mM DHA and was subsequently added three times with 50 mM each, for a total of 200 mM. The black arrow indicates the addition of DHA at this time), 10% (m/v) HP-β-CD with pH maintained at 7.0 ~ 8.0. When considering the use of DHA in scaling up the reaction, only a 100 mL reaction was performed due to the limited availability of DHA at a high cost (about 150 $/g). The results showed that only 3 g/L product was achieved, which is lower than when using AscA as the reductant. This could be attributed to the extreme instability of DHA, leading to rapid hydrolysis in buffer solution ^[9]^. As such, only small fraction of DHA enters the binding pocket and used for initiating the desired reactions. In contrast, AscA undergoes slow oxidation to DHA in the buffer solution, subsequently swiftly entering the hydrophobic binding pocket of UPO to generate Cpd I for catalyzing oxidative reactions. Thus, compared with DHA, the utilization of AscA proves more suitable for practical applications.

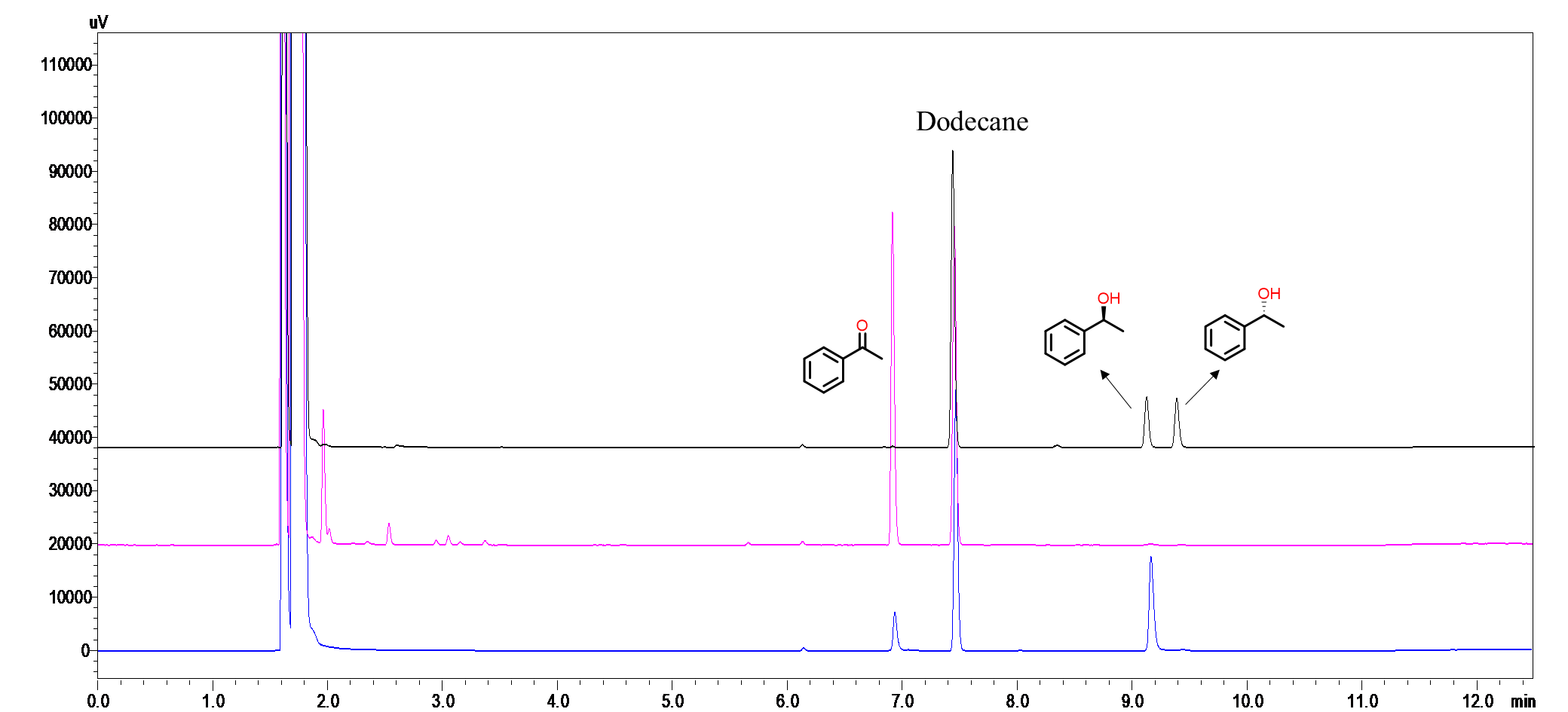

**Supplementary Figure 16.** GC chromatogram of the r*Aae*UPO-catalysed hydroxylation of ethylbenzene. Black: (*R*)-1-phenylethanol and (*S*)-1-phenylethanol standard. Red: acetophenone standard. Blue: samples.

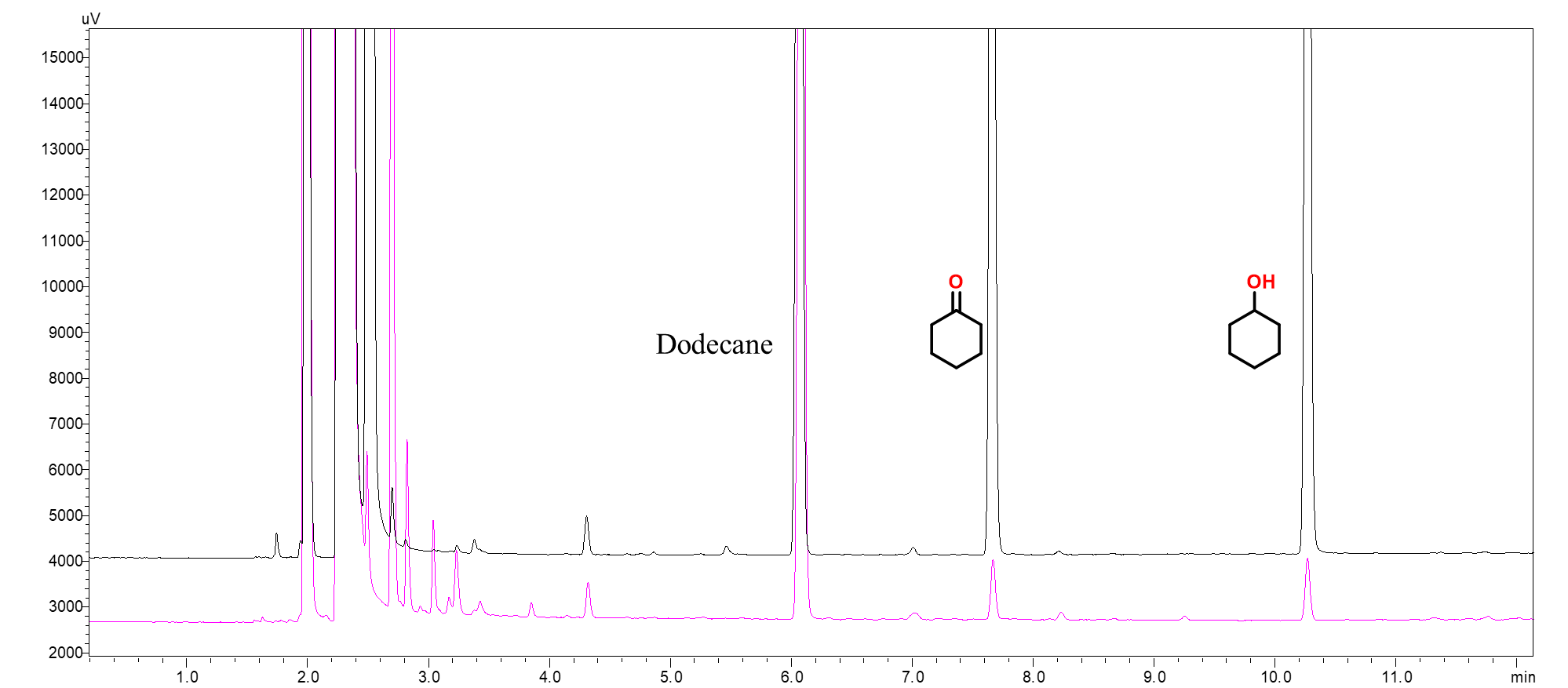

**Supplementary Figure 17.** GC chromatogram of the *Mro*UPO-catalysed hydroxylation of cyclohexane. Black: cyclohexanol standard and cyclohexanone standard. Red: samples

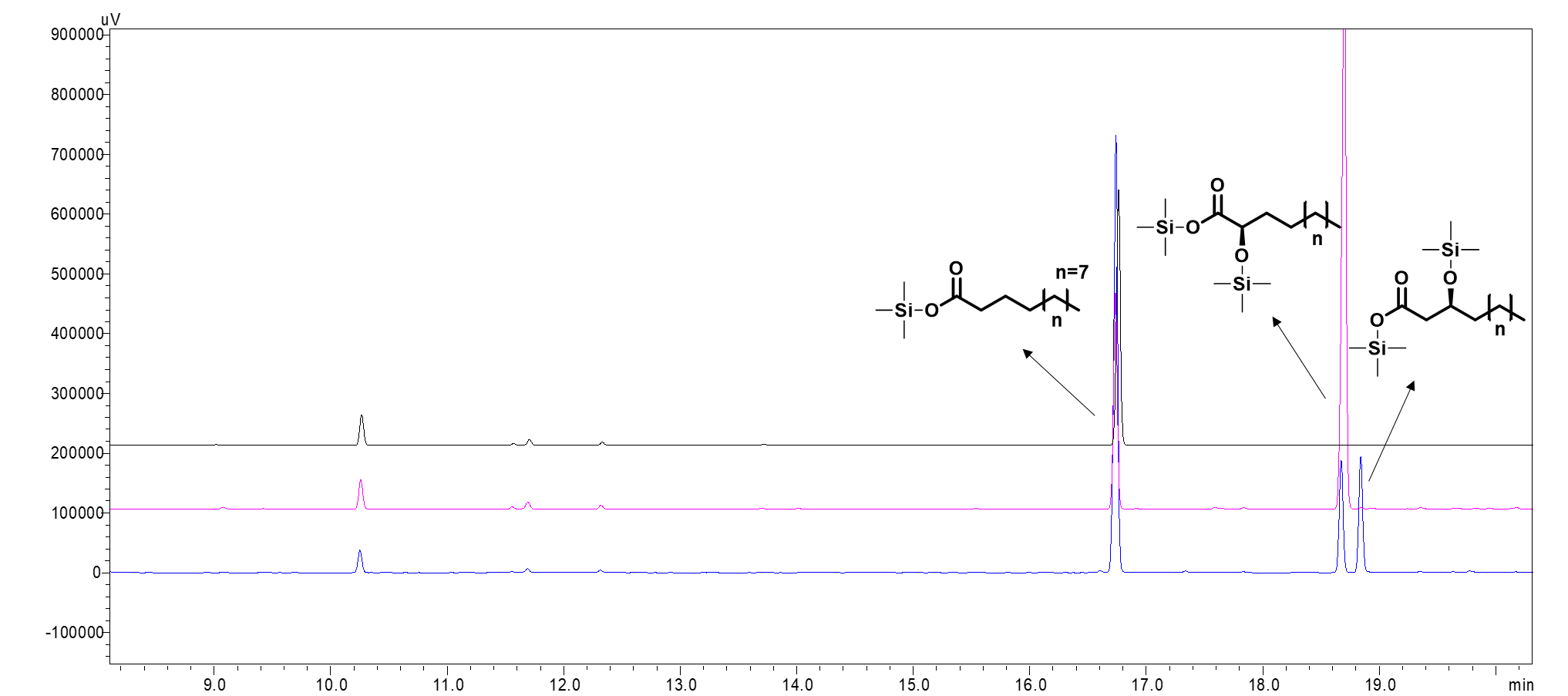

**Supplementary Figure 18.** GC chromatogram of the P450s-catalysed hydroxylation of lauric acid. Black: lauric acid standard. Red: samples of P450_SPα_ after derivatization. Blue: samples of P450_BSβ_ after derivatization.

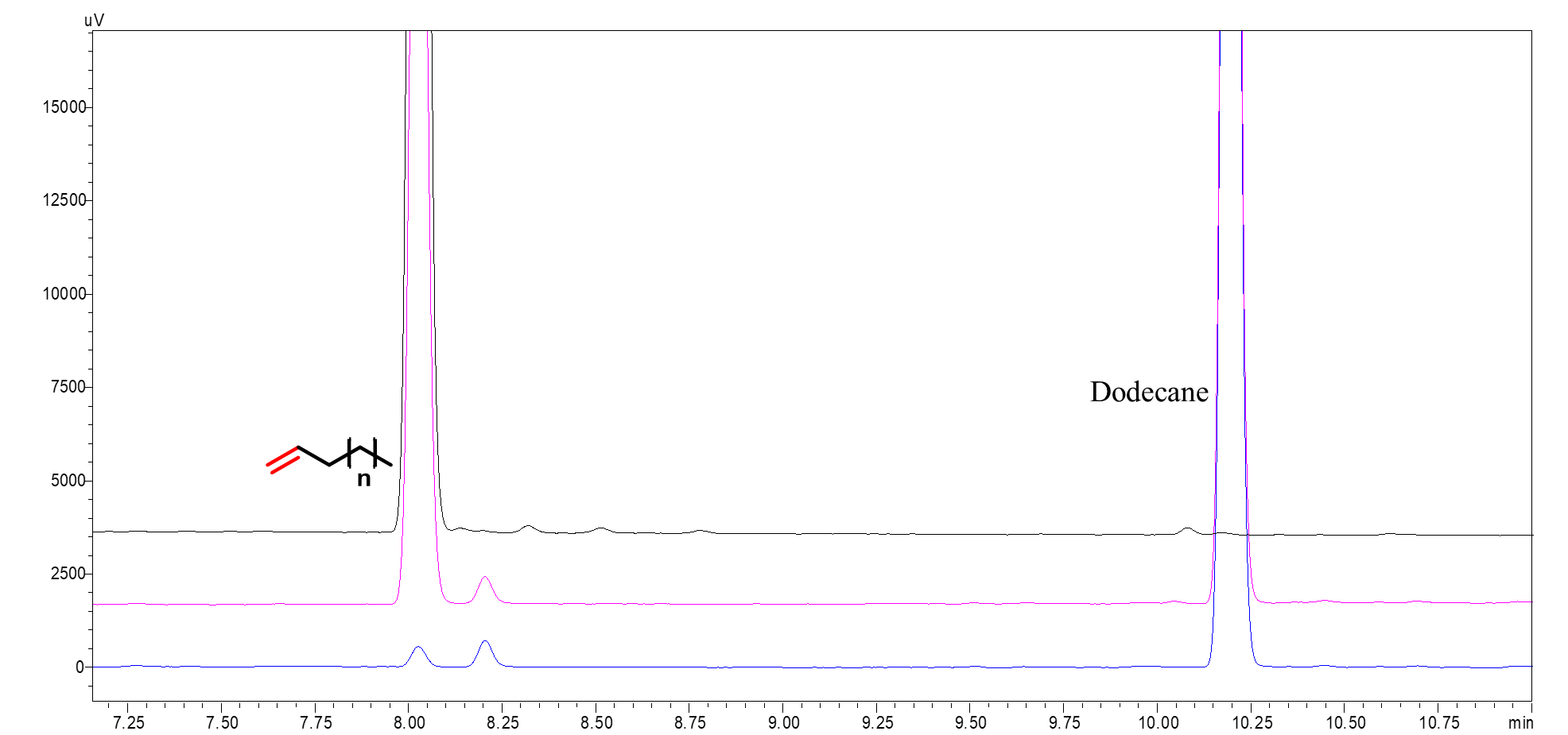

**Supplementary Figure 19.** GC chromatogram of the P450s-catalysed decarboxylation of lauric acid. Black: 1-undecene standard. Red: samples of P450_BSβ_ without derivatization. Blue: samples of P450_SPα_ without derivatization.

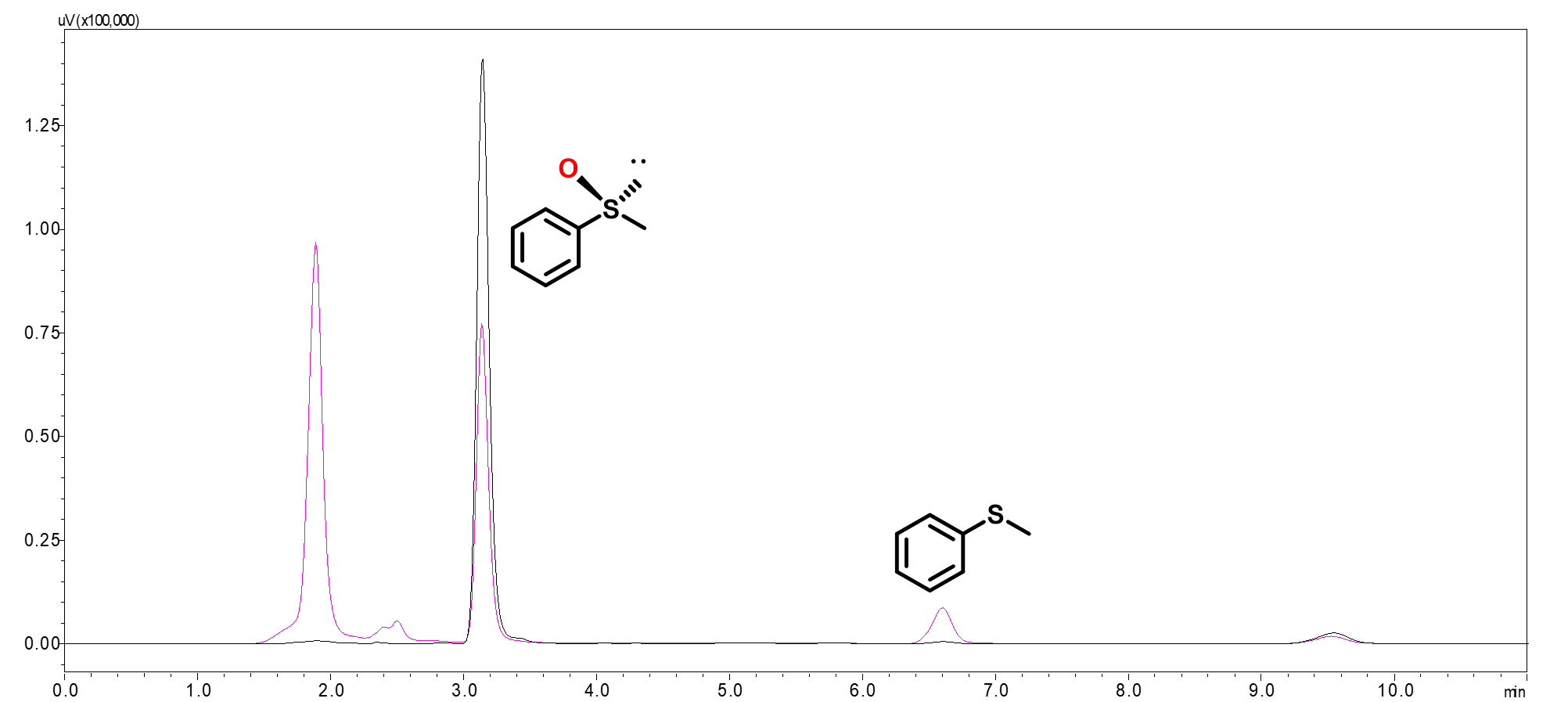

**Supplementary Figure 20.** HPLC chromatogram of the *Cfu*CPO-catalysed hydroxylation of phenyl methyl sulfide. Black: phenyl methyl sulfoxide standard. Red: samples.

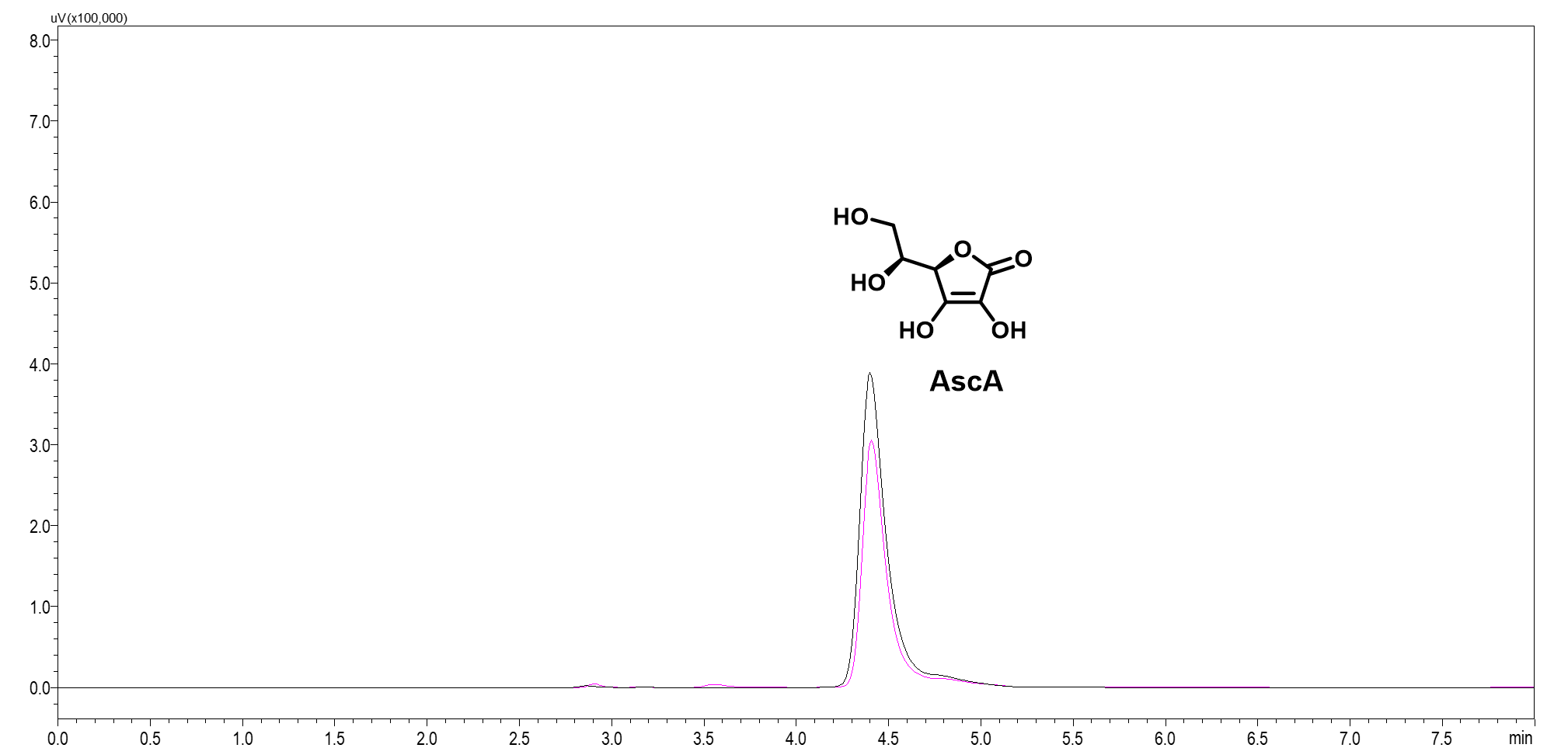

**Supplementary Figure 21.** HPLC chromatogram of the detection of AscA. Black: AscA standard. Red: samples.

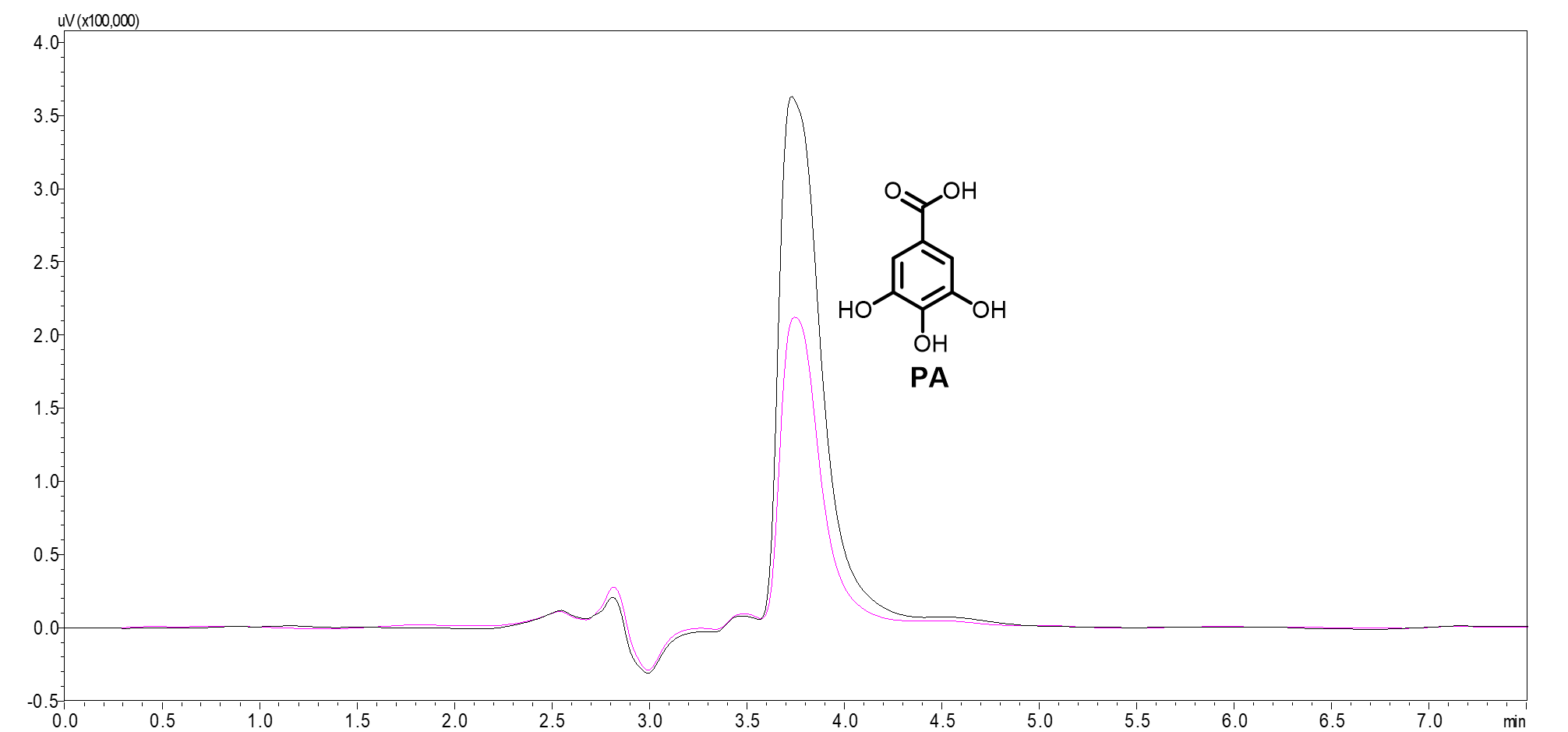

**Supplementary Figure 22.** HPLC chromatogram of the detection of PA. Black: PA standard. Red: samples.

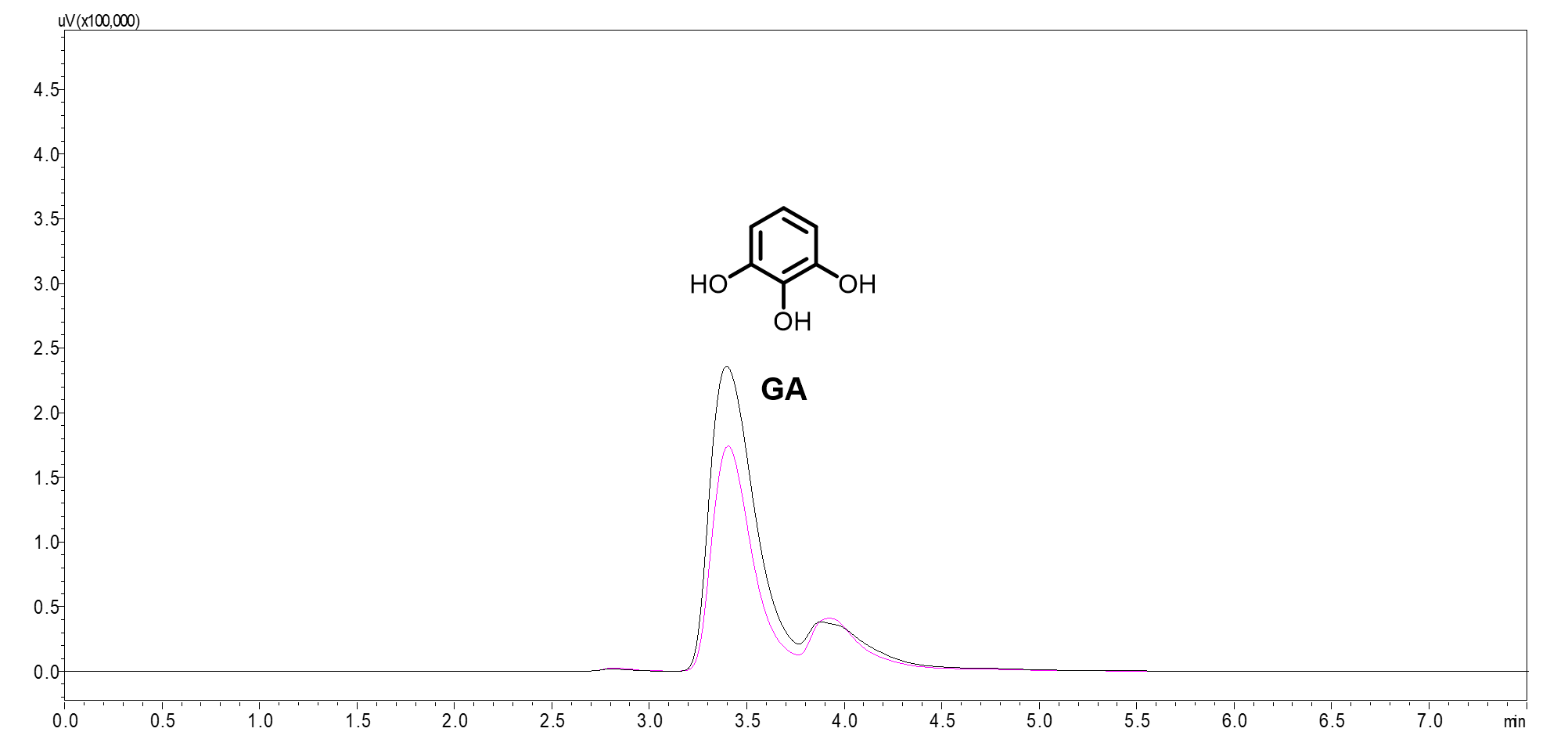

**Supplementary Figure 23.** HPLC chromatogram of the detection of GA*.* Black: GA standard. Red: samples.

**a**

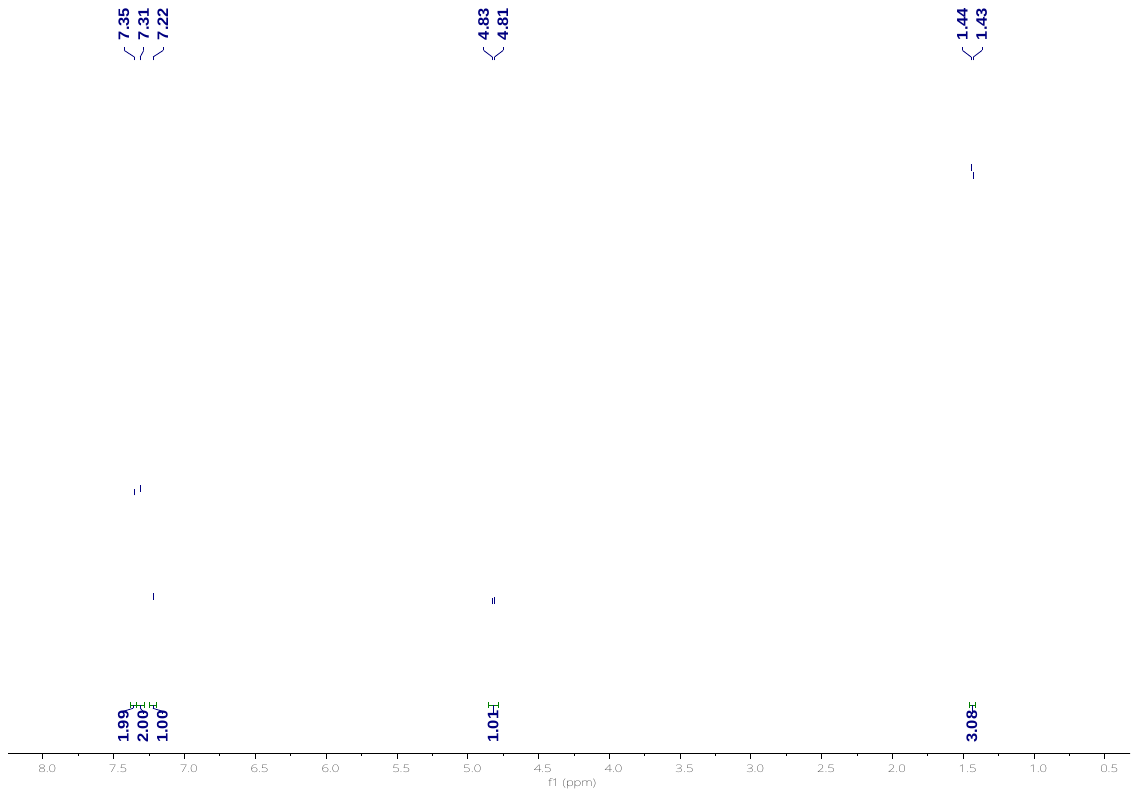

**b**

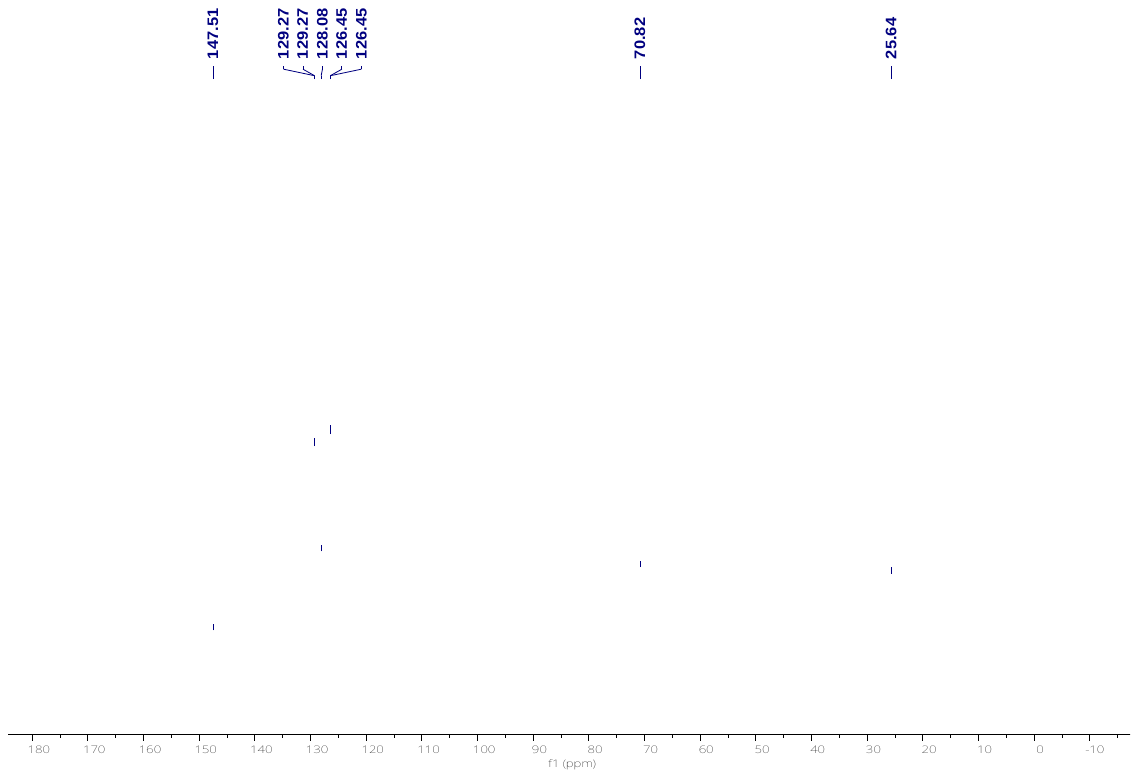

**Supplementary Figure 24.** NMR spectra of 1b. **a.** ^1^H NMR. **b.** ^13^C NMR.

**a**

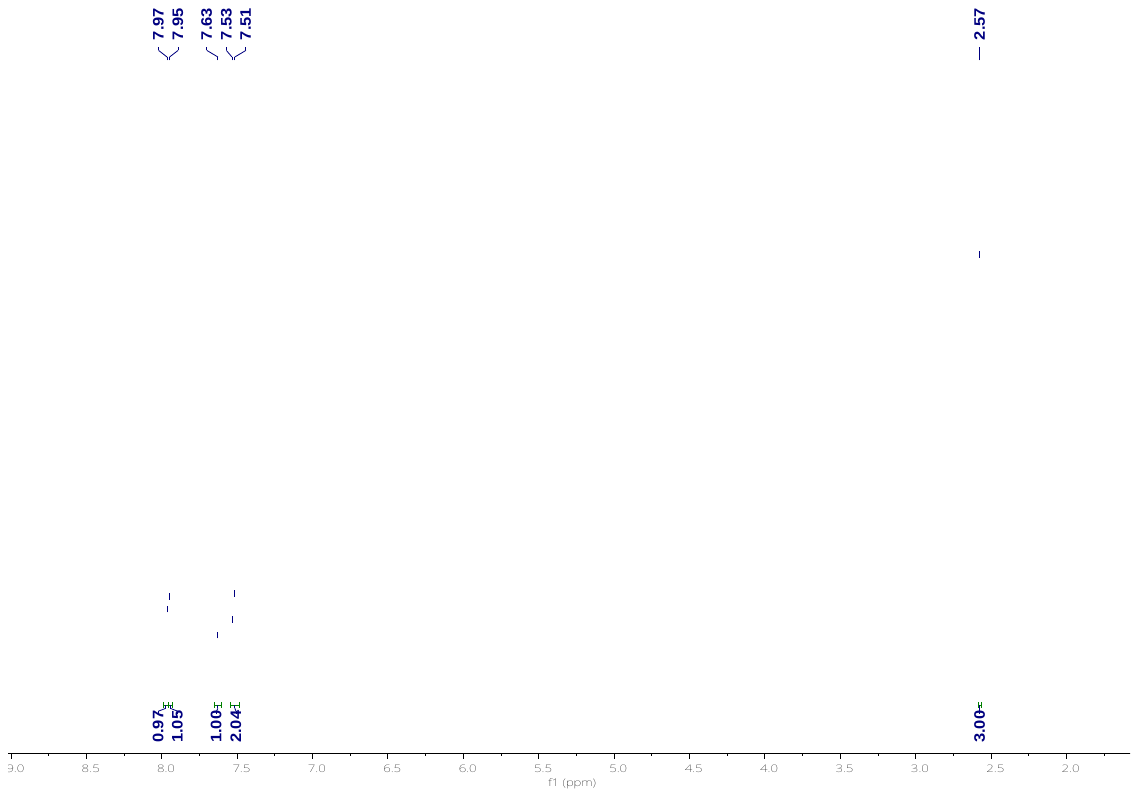

**b**

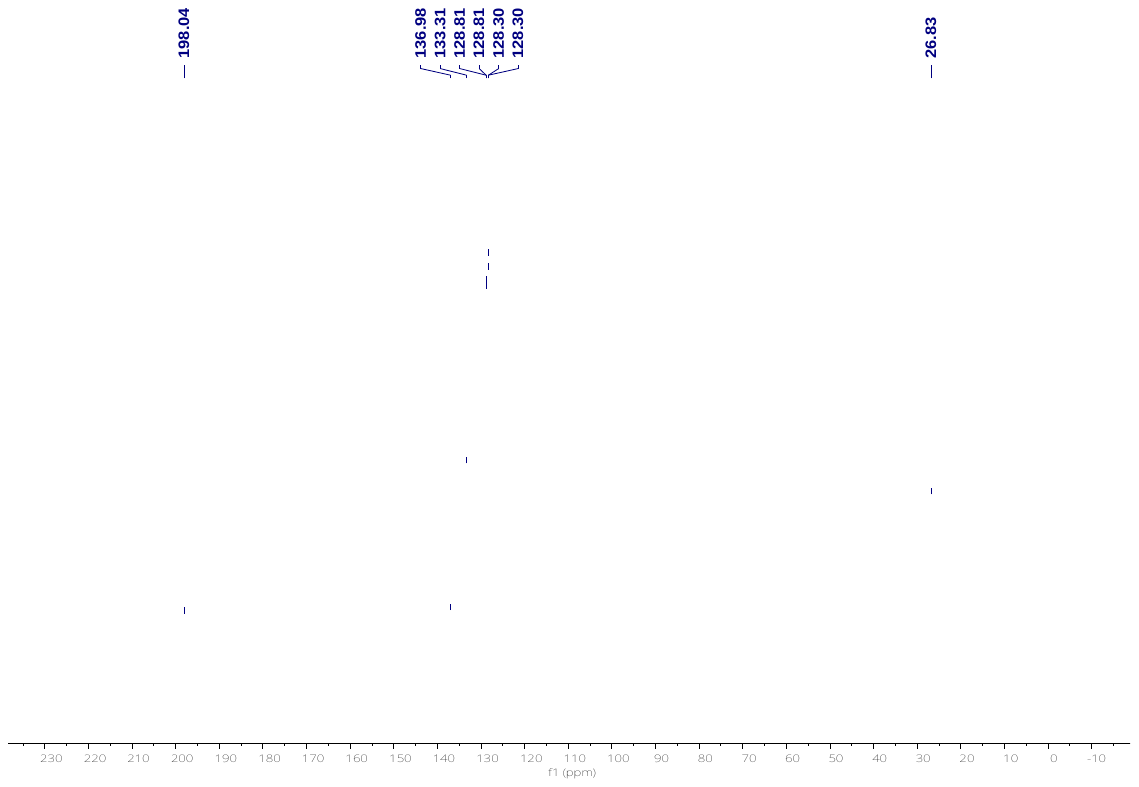

**Supplementary Figure 25.** NMR spectra of 1c. **a.** ^1^H NMR. **b.** ^13^C NMR.

**a**

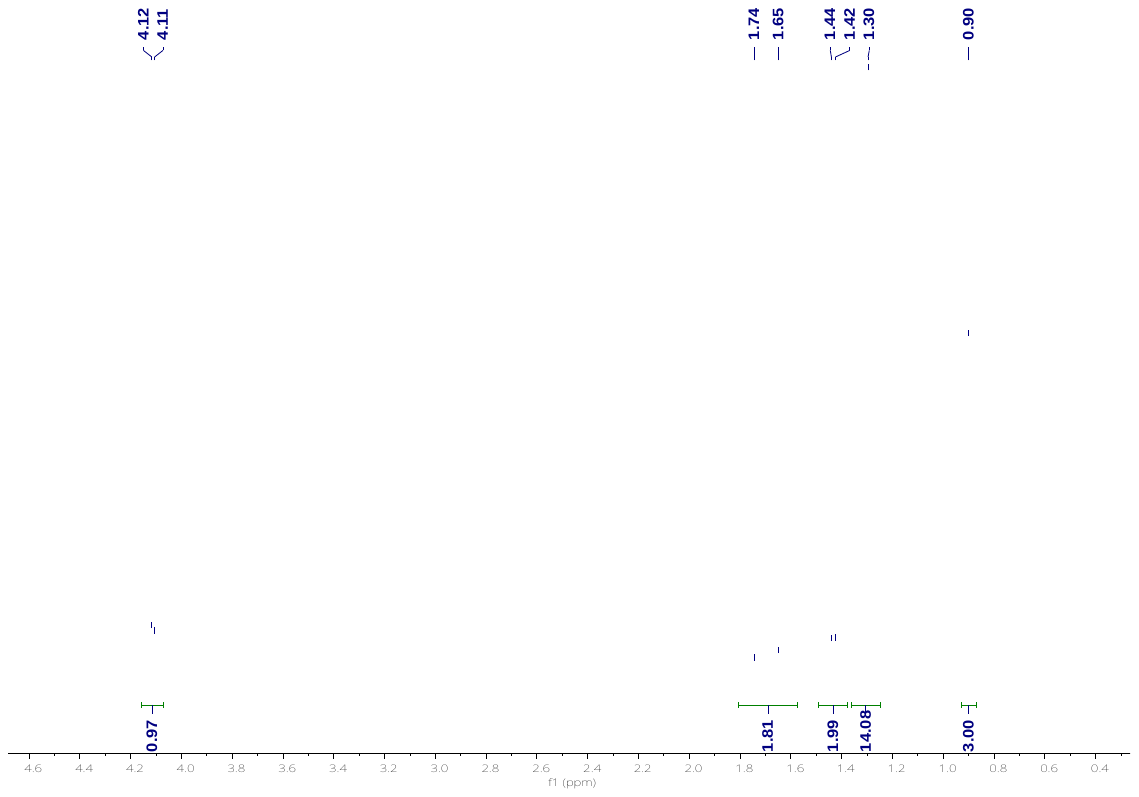

**b**

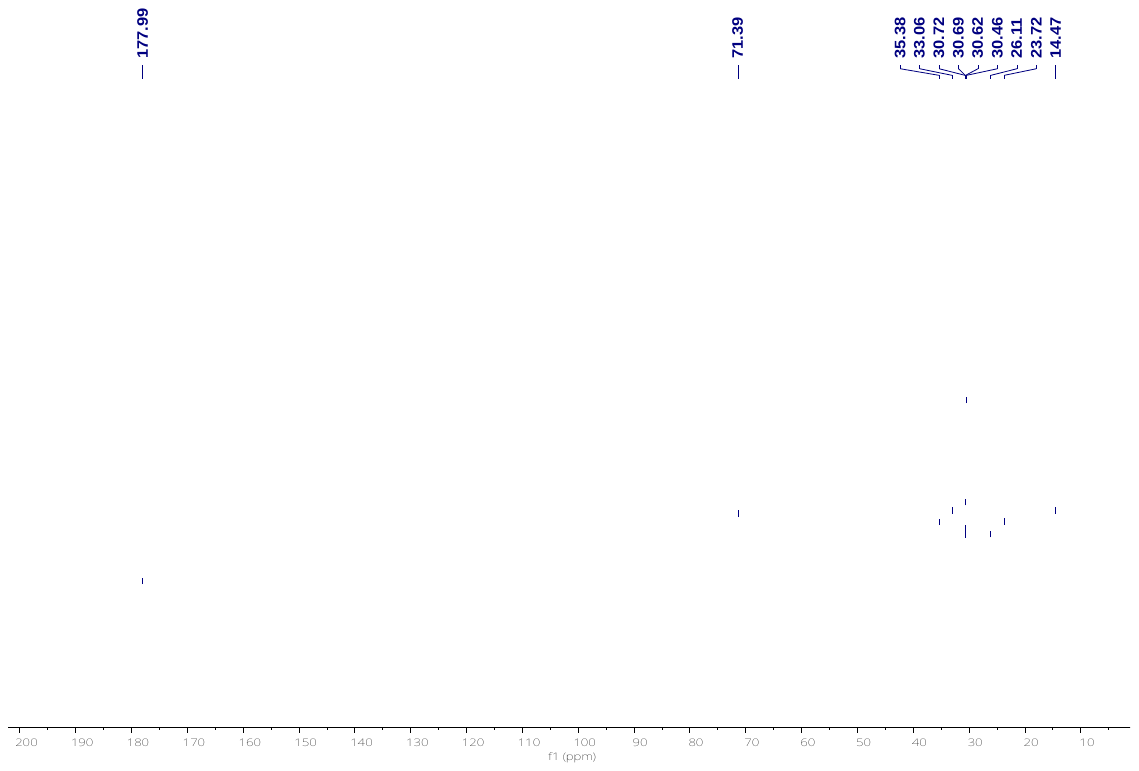

**Supplementary Figure 26.** NMR spectra of 4b. **a.** ^1^H NMR. **b.** ^13^C NMR.

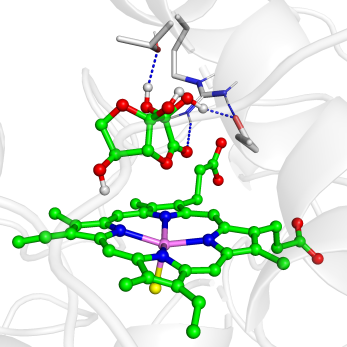

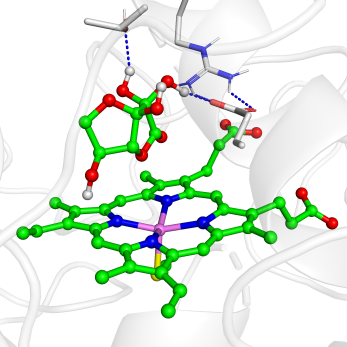

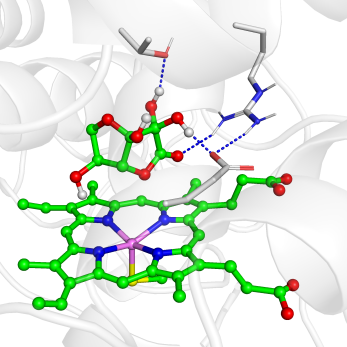

2456/5000

1688/5000

856/5000

**Supplementary Figure 27.** Three most populated structures by clustering of the MD trajectory for the resting state in complex with the substrate DHAA during 50-ns MD simulation. Here we chose the second most populated conformations for the QM/MM calculation of first stage that involves the activation of O_2_ to form the active species of Cpd I.

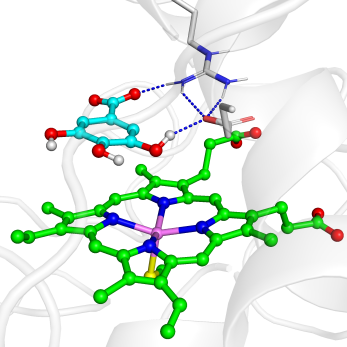

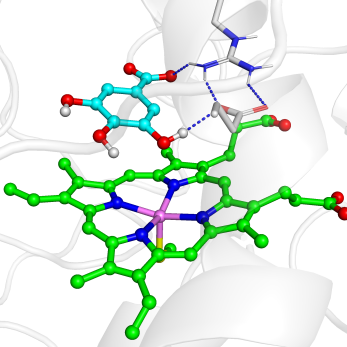

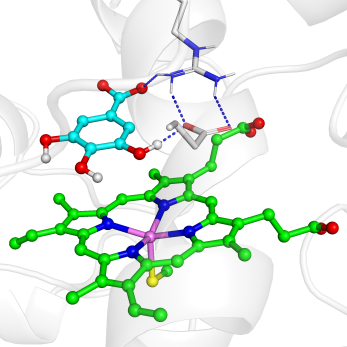

3262/5000

881/5000

857/5000

**Supplementary Figure 28.** Three most populated structures by clustering of the MD trajectory for the resting state in complex with the substrate GA during 50-ns MD simulation. Here we chose the populated conformations for the QM/MM calculation of first stage that involves the oxygen activation and hydrogen peroxide formation.

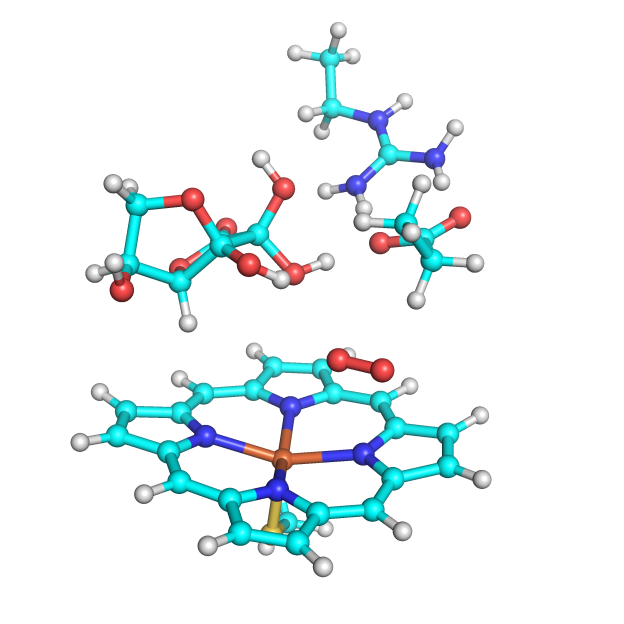

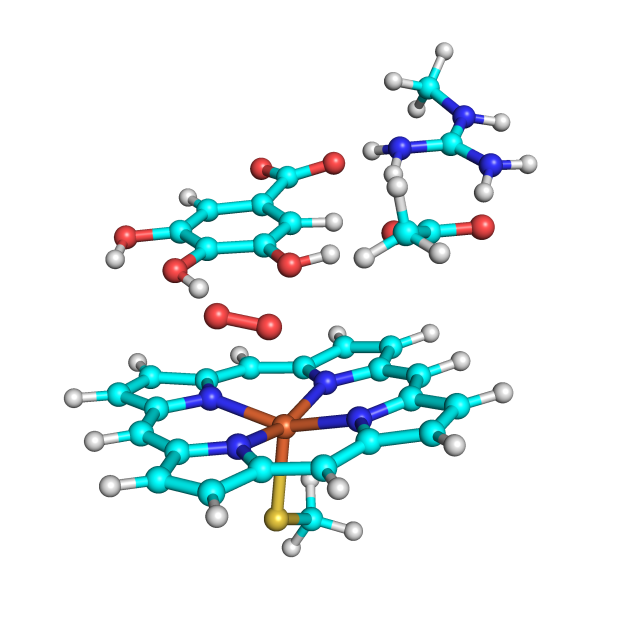

**A**

**B**

**Supplementary Figure 29.** (A) The QM region used for UPO/DHAA/O_2_ system. (B) The QM region used for UPO/GA/O_2_ system.

### Supplementary References

1. Molina-Espeja P, Garcia-Ruiz E, Gonzalez-Perez D, Ullrich R, Hofrichter M, Alcalde M. Directed evolution of unspecific peroxygenase from *Agrocybe aegerita*. *Appl Environ Microbiol* **80**, 3496-3507 (2014).
2. Gröbe G*, et al.* High-yield production of aromatic peroxygenase by the agaric fungus *Marasmius rotula*. *AMB Express* **1**, 31 (2011).
3. Fujishiro T, Shoji O, Nagano S, Sugimoto H, Shiro Y, Watanabe Y. Crystal structure of H_2_O_2_-dependent cytochrome P450_SPα_ with its bound fatty acid substrate: insight into the regioselective hydroxylation of fatty acids at the α position. *J Biol Chem* **286**, 29941-29950 (2011).
4. Lee DS*, et al.* Substrate recognition and molecular mechanism of fatty acid hydroxylation by cytochrome P450 from *Bacillus subtilis*. Crystallographic, spectroscopic, and mutational studies. *J Biol Chem* **278**, 9761-9767 (2003).
5. Sigmund M-C, Poelarends GJ. Current state and future perspectives of engineered and artificial peroxygenases for the oxyfunctionalization of organic molecules. *Nat. Catal.* **3**, 690-702 (2020).
6. Sundaramoorthy M, Terner J, Poulos TL. The crystal structure of chloroperoxidase: a heme peroxidase-cytochrome P450 functional hybrid. *Structure* **3**, 1367-1378 (1995).
7. Soares CM, Gonzalez-Perez D, Molina-Espeja P, Garcia-Ruiz E, Alcalde M. Mutagenic Organized Recombination Process by Homologous In Vivo Grouping (MORPHING) for Directed Enzyme Evolution. *PLoS ONE* **9(3)**: e90919, (2014).
8. Grzesik M, Bartosz G, Stefaniuk I, Pichla M, Namiesnik J, Sadowska-Bartosz I. Dietary antioxidants as a source of hydrogen peroxide. *Food Chem.* **278**, 692-699 (2019).
9. Deutsch, J. C. Dehydroascorbic acid. *J.* *Chromatogra.* **881**, 299-307 (2000).
